# Leveraging mathematical models to predict and control T-cell activation

**DOI:** 10.1101/2025.11.20.689432

**Authors:** Xabier Rey Barreiro, Jose Faro, Alejandro F. Villaverde

**Affiliations:** CITMAga, 15782 Santiago de Compostela, Galicia, Spain; Universidade de Vigo, Department of Systems Engineering and Control, 36310 Vigo, Galicia, Spain; Universidade de Vigo, Department of Biochemistry, Genetics and Immunology, 36310 Vigo, Galicia, Spain; Immunology Research Group, Galicia Sur Health Research Institute (IIS Galicia Sur), SERGAS-UVIGO, Spain

## Abstract

T-cell receptor (TCR)-mediated T-cell activation is a key process in adaptive immune responses. The complexity of this process has led to the development of different mathematical models that seek to describe and predict the conditions of antigen-TCR interactions required for TCR triggering and T-cell activation. These models are characterized by describing different sets of sequential molecular interactions and their kinetics, positing the generation of a final product as a necessary and sufficient condition for T-cell activation. Such modeling could provide an effective tool for simulating antigen recognition by T cells and, consequently, aid in the design of effective therapeutic strategies. However, it is necessary to previously assess the predictive capabilities of the proposed models when fitted to experimental data. To achieve this goal, in this work we examine the parameter identifiability and sensitivity of the published models of TCR-based T-cell activation. For each model, we consider different, often experimentally measured, output quantities and show how their availability affects the results. These analyses allow us to determine the ability of each model to correctly describe different experimental situations, and to establish to what extent these models can be applied to reliably predict and control T-cell activation by specific therapeutic targets.

**Author summary:** The adaptive immune system is a highly specialized defense mechanism present in vertebrates. It produces a response that results from a complex set of interrelated cellular and molecular mechanisms. A key process is antigen-dependent activation of T lymphocytes, for which several mechanisms have been postulated. The corresponding mathematical models allow us to explore competing hypotheses, make quantitative predictions, and eventually aid in the design of strategies to control the immune response. To achieve these goals, it is essential that the models be identifiable and observable, that is, it must be possible to infer their parameters and state from output measurements. Furthermore, their response should exhibit an adequate level of sensitivity to certain key parameters. In this work, we evaluate the degree to which currently existing models possess these qualities. Our findings provide minimum requirements for the design of system identification experiments, allowing us to discard those models that cannot be successfully calibrated with a given experimental setup.

## Introduction

Adaptive immune responses to protein antigens (Ag) are based on interactions of T lymphocytes with target or Ag-presenting cells (APC) that lead to the activation of Ag-specific T cells. This lymphocyte activation is initiated by the binding of T-cell receptors (TCR) to their ligands, Ag-derived short peptides complexed with major histocompatibility complex proteins (pMHC) expressed on the membrane of APCs [1]. TCRs are membrane proteins composed of two chains, TCR*α* and TCR*β*, each with two extracellular globular domains. They bind pMHC ligands by their two membrane-distal or variable domains but lack the capacity to directly transduce a signal intracellularly, owing to their very short intracellular tails. However, they can signal indirectly through their tight, non-covalent association with a membrane invariant multimer termed CD3, composed of two heterodimers (CD3*ϵ*δ, and CD3*ϵγ*) and one homodimer (CD3ζζ) [1]. The four chains *ϵ*, δ, *γ*, and ζ, all have a single globular extracellular domain and a large intracytoplasmatic tail, with ζ having the larger one. The CD3 chains *ϵ*, δ, and *γ* have in their tail one signaling motif, containing two tyrosines, termed immunotyrosine activation motif (ITAM), while CD3 ζ chains contain two ITAMs [1, 2]. There is evidence that TCR-pMHC binding induces a conformational change in the membrane proximal constant domain of the TCR*β* chain that is transmitted to the CD3 extracellular domains, which drives the conversion of CD3 intracellular tails from an inactive to an active form [1, 3–6]. This allows the phosphorylation of CD3 ITAMs, which, in turn, triggers an intracellular molecular signaling cascade that eventually leads to T-cell activation. The degree of that signaling depends to a variable extent on the pMHC concentration and the kinetic rate constants of both the TCR-pMHC interaction and the CD3 phosphorylation events [7–9]. Phosphorylated TCRs that reach the signaling state are denoted activated or triggered TCRs.

Due to the complexity of those processes, the relationship between these quantities and T-cell activation is not well understood yet. Over the years, numerous quantitative experimental data have been obtained (reviewed in [1]), inspiring the formulation of several mathematical models, most of them with a common core based on the kinetic proofreading concept [10]. In each of these models, an output can be defined that corresponds to the dynamics of a selected set of experimentally measurable quantities (phenotypic features of activated T cells) and that define a T-cell activation phenotype [11–13]. However, it becomes challenging to determine which of those models (from now on, denoted phenotypic models) best describes T-cell activation.

Model parameters that cannot be directly measured are usually estimated by fitting the model output, that is, measurable quantities of the model dynamics, to experimental data [14, 15]. However, finding the optimal fit does not guarantee that the corresponding parameter estimates are correct or meaningful. This issue is particularly important for those parameters that have a significant impact on the time-varying output, so that even a small variation in them has a big effect on model predictions. Therefore, before attempting to calibrate a model or use it to predict the behavior of a particular biological system under study, it should be ensured that there are no model structural issues that can lead to unreliable and wrong model predictions [16, 17].

As a step to achieve this ultimate goal, here we perform extensive analyses of most (if not all) published phenotypic models of T-cell activation. In particular, we examine the characteristics of these models with respect to their *parameter identifiability, state observability*, and *output sensitivity* to parameter values. Identifiability is the property that describes whether it is possible to determine the values of unknown parameters from the model output [18]. It is tightly linked with observability, which is a property that determines the feasibility of inferring the unmeasured dynamic variables, or states, from the model output [19], thus informing about a model’s ability to make reliable predictions about the state of a system over time. Lastly, sensitivity analysis evaluates the extent the model output is affected by variations in parameter values [20, 21]. These properties depend on which variables can be experimentally measured, also known as model outputs. Therefore, in our analyses, we consider as outputs different potentially measurable quantities like the amount of total, free, and activated or triggered TCRs per T cell (the latter as a surrogate of T-cell activation level). In addition, we also analyze as outputs two frequently measured quantities, the maximum ligand effect (*E*_*max*_) and the half-maximal effective ligand concentration (*EC*_50_).

In this article we address, thus, the following questions: First, which phenotypic models or model structures, i.e. which sets of dynamic and output equations can be used to predict T-cell activation reliably, considering the need to estimate their parameters from fitting procedures (identifiability)? Second, to what extent is the behavior of those phenotypic models influenced by the particular values of said kinetic parameters (sensitivity)? Finally, how can this combined knowledge about parameter identifiability and sensitivity be leveraged to determine which parameters are the best candidates to be tuned experimentally for driving T-cell activation in a controlled way?

## T-cell Activation Models

Mathematical models of T lymphocyte activation have been developed to elucidate how a signaling network translates Ag dose and kinetic parameters into a T-cell response [12, 22, 23]. These models aim to analyze the biochemistry of signaling in detail. However, given their complexity and the wide range of features they must account for, it has been suggested to reformulate these models into simplified versions known as *phenotypic models* [11]. Phenotypic models of T-cell activation focus on explaining the observable characteristics of the T-cell response in terms of a few key biochemical processes rather than the intricate biochemical details. The objective is to build a relatively simple yet efficient model capable of reproducing the observed phenotypes [13].

Any model designed to describe T-cell activation quantitatively must be capable of considering and quantifying the following key features of the adaptive immune response: sensitivity to detect foreign pMHCs at very low concentrations amid a significantly higher abundance of self-pMHC ligands, (2) discrimination or specificity to distinguish between foreign and self-ligands and not allow self-peptides to trigger an immune response, and (3) a short reaction time to foreign peptides, ensuring these requirements are met promptly [24]. This section briefly describes the T-cell activation phenotypic models we have found in the literature (listed in Table 1). A general diagram of the mechanisms they describe is depicted in Fig. 1, and for each model, its specific conceptual scheme and equations are given in the Supplementary Information.

**Table 1.** Phenotypic models analysed in this work.

| Model | Short name | Reference |
| --- | --- | --- |
| Occupancy | **Occ** | [25] |
| Kinetic proofreading (McKeithan) | **KPR-1** | [10] |
| KPR with limited signaling | **KPR-LS** | [11] |
| KPR with sustained signaling | **KPR-SS** | [11, 26] |
| KPR with induced rebinding | **KPR-IR** | [27] |
| KPR with limited signaling and incoherent feedforward loop | **KPR-LS-IFF** | [12] |
| KPR with negative feedback I | **KPR-NF1** | [24] |
| KPR with negative feedback II | **KPR-NF2** | [2] |
| KPR with stabilizing activation chain | **KPR-SAC** | [22] |
| Zero-order ultrasensitivity | **ZU** | [28] |
| KPR with zero-order ultrasensitivity | **KPZU** | [28] |
| KPR with concentration compensation | **KPC** | [28] |
| Serial triggering | **ST** | [29] |

**Fig 1.**
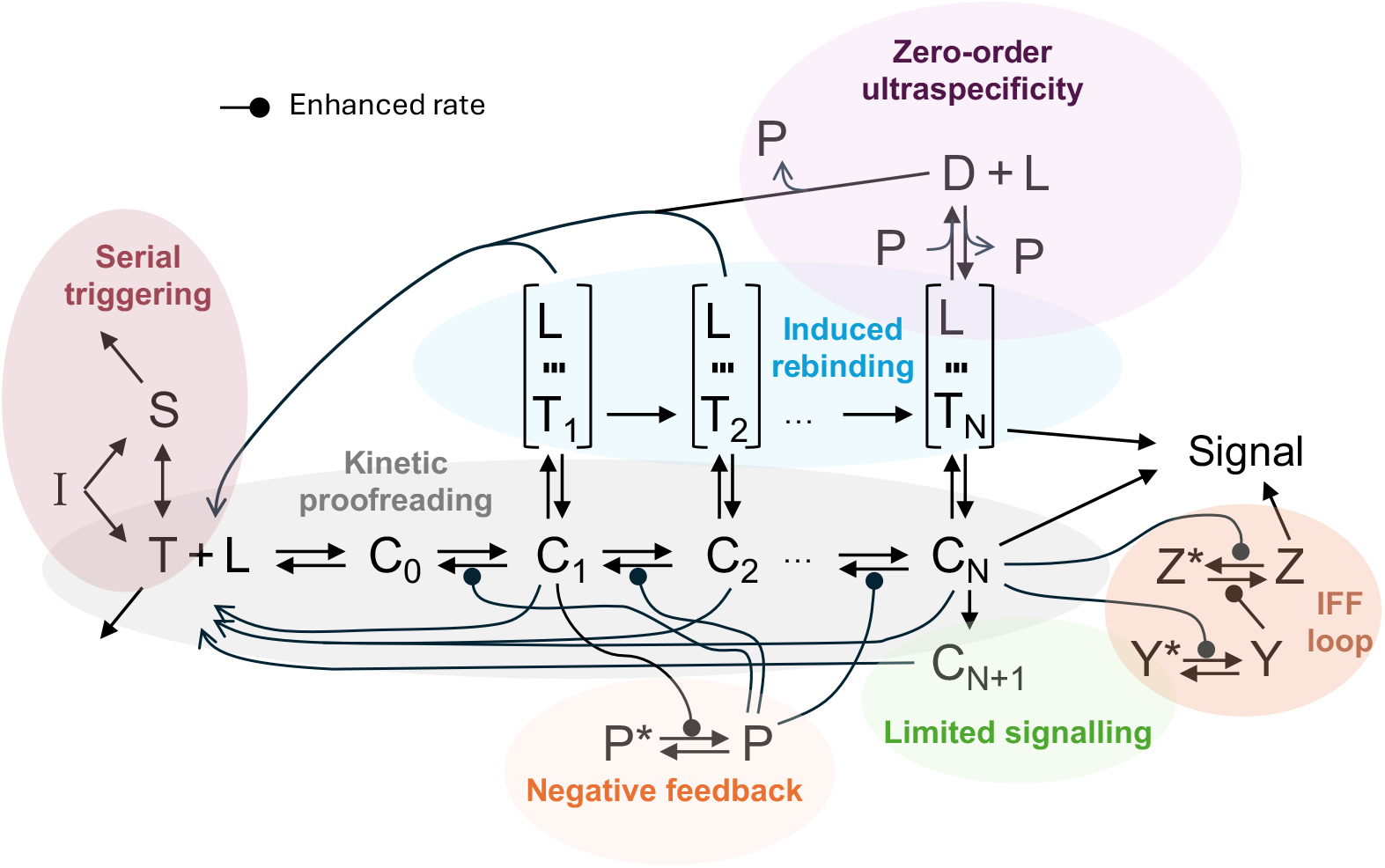
Master diagram of the models analyzed in this work. Each model describes a subset of the interactions shown in the figure. Parts corresponding to specific mechanisms are colored. *Common model variables*: *T* denotes free TCRs, *L* represents free pMHC ligands, *C* denotes TCR-ligand complexes, and *N* is an integer number that refers to the number of proofreading steps required for TCR triggering. *Zero-order Ultraspecificity*: *P* denotes free phosphatase and *D* denotes a complex of a phosphatase with a triggered TCR. *Negative Feedback*: *P* and *P*^∗^ denote, respectively, active and inactive SHP-1 phosphatases. *Induced rebinding*: *T*_*i*_ denotes TCRs at proofreading step *i* that have recently unbound from their pMHC ligand but are still very close to it, in an intermediate state, such that they have the potential to rebind it again; this receptor-ligand state is indicated with three short lines. *Serial triggering*: *S* refers to TCRs in the membrane spare pool of T cells. *Limited signaling*: *C*_*N*+1_ refers to TCR-ligand complexes that underwent one proofreading step beyond the TCR-triggering one. *Incoherent feed-forward (IFF) loop*: *Y* denotes an intermediate metabolite required to enhance the transformation of a signaling molecule (denoted *Z*) from an inactive state to an active state.

### Occupancy Model (*Occ*)

The occupancy model [25] posits that T-cell activation is proportional to the concentration of TCR-pMHC complexes. It assumes, therefore, that TCRs reach a signaling-competent state immediately after binding a pMHC.

### Kinetic Proofreading Models (*KPR*)

The kinetic proofreading (KPR) mechanism was initially proposed by J. Hopfield [30] and J. Ninio [31] to explain how enzymes lead to correct products over incorrect ones with high accuracy. A couple of decades later, this mechanism was adapted by McKeithan to the case of T-cell activation as a possible way to explain how T cells achieve precise discrimination between cognate foreign and self-pMHC ligands, despite the overwhelming presence of the later [10]. This concept posits that the interaction between TCRs and pMHC ligands does not immediately result in signaling upon docking. Instead, the TCR must undergo a series of post-translational modifications and spatial rearrangements before initiating effective signaling into the cell. During this process, if the TCR-pMHC complex dissociates prematurely due to high unbinding rates, the signaling process is halted and restarted, thereby preventing incorrect activation. Only TCR-pMHC complexes with sufficiently small unbinding rates persist long enough to complete all the necessary steps and trigger an effective immune response. Over the years, different versions and extensions of McKeithan’s original KPR model have been proposed; they are described below.

#### KPR McKeithan (*KPR-1*)

This model is perhaps the simplest representation of lymphocyte receptor activation through kinetic proofreading [10]. According to it, TCR binds reversibly to pMHC, and bound TCRs initiate a sequential process of phosphorylation at the CD3 intracellular ITAM motifs. At each TCR phosphorylation stage, if the pMHC unbinds the TCR, this reverts to its basal unmodified state. Only those pMHC-bound TCRs that have gone through a threshold number of steps (*i*.*e*., that have accumulated a threshold number of phosphorylations) become triggered and start signaling T-cell activation.

#### KPR with Limited Signaling (*KPR-LS*)

This model extends the *KPR* model by postulating that pMHC-bound TCRs that reach the triggering state can be further phosphorylated into a state in which they lose the signaling ability before unbinding pMHC. Its development was motivated by experimental observations on serial triggering suggesting that efficient T-cell activation requires that the TCR-pMHC dwell time lies within an optimal range [32]. This model, proposed in [11], incorporates into a *KPR* framework the serial triggering feature that triggered TCRs signal T cells for a fixed period of time, leading to an optimum T-cell activation level with respect to the TCR-pMHC dissociation time.

#### KPR with Sustained Signaling (*KPR-SS*)

This model was developed following experimental observations indicating that at high ligand densities optimal T-cell activation also takes place at extended TCR-pMHC dwell times and not only within an optimal range [33]. Thus, unlike the *KPR-LS* model, in the *KPR-SS* model pMHC-bound TCRs that have reached the triggering state dissociate reversibly from the pMHC, and recently dissociated, triggered TCRs remain signaling for some time before reverting to the basal state [11].

#### KPR with Induced Rebinding (*KPR-IR*)

The *KPR-1* model predicts high specificity but reduced receptor sensitivity. This occurs because, as the number of phosphorylation steps needed to complete the phosphorylation of a bound TCR and trigger it (phosphorylation threshold) increases, so does the time required for TCR triggering, and therefore the proportion of TCRs that remain bound to pMHCs decreases exponentially. Thus, there is a trade-off such that a larger phosphorylation threshold leads to higher specificity but lower sensitivity, rendering T cells less capable of responding to low concentrations of specific antigens. In order to correct for this shortcoming, the *KPR-IR* extension accounts for the possibility that recently unbound TCRs remain for a time in a state and at a distance that facilitate their subsequent binding to the same pMHC [27]. This extension assumes, thus, that the initial binding of a pMHC ligand to a TCR induces in the receptor some changes, such as TCR clustering and/or conformational alterations, that could significantly increase rebinding to that pMHC. This mechanism enhances both sensitivity and specificity. We analysed an approximate version of this model derived in [34]. In this approximation, we assume that rebinding after unbinding occurs fast enough so that bound states and unbound but encounter-oriented states can be merged into one effective state [35].

#### KPR with Limited Signaling and Incoherent Feed-Forward Loop (*KPR-LS-IFF*)

Shortly after proposing the *KPR-LS* model, its authors performed a series of *in vitro* experiments, systematically analyzing T-cell responses to 10^6^-fold variations in both pMHC concentration and TCR-pMHC 3-dimensional effective affinity. They observed four key phenotypic features related to dose-response and peak amplitude of the response [12]. This led them to explore systematically thousands of variants of the *KPR* model of increasing complexity to find those able to explain the observed features of T-cell responses. They concluded that a *KPR-LS* model with a coupled incoherent feed-forward (IFF) loop is the simplest model able to explain the four phenotypic features. The feed-forward loop consists of a signaling network where pMHC-bound, triggered TCRs activate two metabolic reactions: one that indirectly activates the final signaling process and another one that directly inhibits it (detailed in Fig. 1).

#### KPR with negative feedback, variants I and II (*KPR-NF1, KPR-NF2*)

Experimental findings showing that the Src homology 2 domain phosphatase-1 (SHP-1) has an important impact on T-cell activation by regulating the phosphorylation level of pMHC-bound TCRs [36] led to another extension of the basic *KPR* model. According to it, TCRs in complexes *C*_1_ of the proofreading chain activate a phosphatase which, in turn, reverses with similar rate constant each phosphorylation step in the proofreading chain [24]. This model, KPR with negative feedback I (in short, *KPR-NF1*), can explain the above mentioned three key properties in T-cell signaling: speed, specificity, and sensitivity. Recent experimental results prompted the same authors to modify the *KPR-NF1* model by postulating that the TCRs in the last *r* complexes of the proofreading chain can contribute to signaling the T cell, but with increasing contribution from *C*_*N−r*+1_ to *C*_*N*_ [2]. Notice that this new model, denoted here *KPR-NF2*, has the same ODEs as the *KPR-NF1* model.

#### KPR with stabilizing/destabilizing activation chain (*KPR-SAC*)

This extension addresses the characteristic trade-off between specificity and sensitivity by postulating that for TCR-pMHC complexes with agonist peptides, the unbinding rate (*k*_*off*_) decreases as the complexes progress through the proofreading chain, while the propagation rate (*k*_*p*_), with which TCR-pMHC complexes proceed along the proofreading chain, increases. For non-agonist peptides, however, the *KPR-SAC* model postulates that *k*_*off*_ increases and *k*_*p*_ decreases as the complexes proceed along the proofreading chain [22].

### Zero-order ultraspecificiy

More than forty years ago, Goldbeter and Koshland [37] proposed the so-called zero-order ultrasensitivity (*ZU*) mechanism to explain enhanced sensitivity to changes in enzyme concentration in an enzymatic system in which the enzyme operates in saturating conditions with respect to substrate. Recently, the *ZU* mechanism was applied to T-cell activation by TCR-pMHC complexes [28]. Based on that mechanism, the authors presented three models of increasing complexity, from a basic *ZU* model to a model combining the *ZU* and the *KPR* mechanisms, all of which are briefly described below.

#### Zero-order ultrasensitivity (*ZU*)

In this adaptation of the zero-order ultrasensitivity mechanism to TCR-driven T-cell activation, pMHC ligands play the same role as the first enzyme in the original model, and TCRs and phosphorylated TCRs act as substrate and product, respectively. Here, the role played by the second enzyme in the original model is ascribed to a phosphatase. Thus, when nearly all cognate pMHC ligands are bound to TCRs, the sensitivity of the system to small changes in ligand concentration is highly enhanced. Unlike multistep proofreading mechanisms, zero-order ultrasensitivity increases sensitivity and enables rapid responses, while compromising specificity, that is, reducing the ability to discriminate against self-antigens [28].

#### KPR with zero-order ultraspecificity (*KPZU*)

The KPR with zero-order ultraspecificity model, *KPZU* for short (*GKP* in the original paper [28]), combines the kinetic proofreading with the zero-order ultrasensitivity mechanism. This combination enables it to balance specificity, sensitivity, and speed effectively.

#### KPZU with concentration compensation (*KPC*)

The *KPC* model simplifies the *KPZU* model by assuming that the same molecule regulates both TCR phosphorylation and dephosphorylation [28]. This allows for ligand concentration compensation, enabling the system to remain insensitive to high concentrations of non-target ligands while maintaining sensitivity to low concentrations of target ligands.

### Serial triggering (*ST*)

In 1995, Valitutti and Lanzavecchia found that on average a single pMHC molecule can engage a large number of TCRs, triggering their phosphorylation and internalization (downregulation), and eventually activate T cells [38]. Based on this finding, they proposed the serial triggering (*ST*) conceptual model, according to which a small number of pMHC complexes can activate a T cell by binding, each of them, to a TCR long enough to trigger its phosphorylation and downregulation, and then dissociating from this receptor and binding to another TCR, thereby sequentially triggering many different receptors. Subsequently, Ohashi and *col* developed a mathematical model to analyze this finding [39]. They proposed the following dimerization model of TCR-pMHC complexes to explain the observed kinetics of TCR downregulation, and hence of serial triggering:

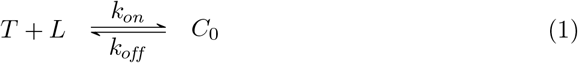

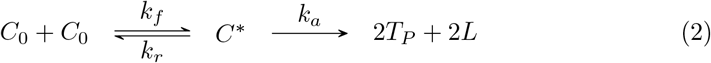

where *T* is the free TCR density on a T cell, *L* is the ligand (pMHC) density on an APC, and *C*_0_, *C*^∗^ and *T*_*P*_ are the densities of TCR-pMHC complexes, *C*_0_ dimers, and triggered TCRs, respectively. Assuming that *C*_0_ formation reaches fast a quasi-steady state with *T* and *L*, and assuming also a quasi-steady state approximation for dimers, the authors arrived at the following equation for triggered TCRs [39]:

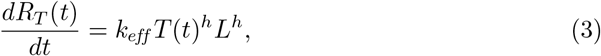

where *k*_*eff*_ is an effective binding rate defined by 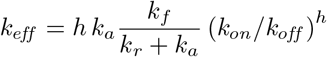, and *h* = 2.

To account for experimental results not explained by this model, other authors extended it a few years later by incorporating additional features of TCR dynamics [29]. In the enhanced *ST* model, TCRs are partitioned into two subsets, one consisting of the TCRs within the membrane interface or contact area between a T cell and an APC (named the interface pool), and another with those TCRs outside the interface area (named the spare pool). Free TCRs are assumed to diffuse freely from one pool to the other, and to turn over between the membrane and the cytoplasm. In this model, only TCRs of the interface pool can interact with pMHC molecules presented by an APC, and they do it irreversibly, with effective kinetic rate *k*_*eff*_, as defined above, and kinetic order *h >* 1, leading to triggered (phosphorylated) TCRs, which are then internalized with rate *k*_*i*_ [29].

The original mathematical formulation of the *ST* model suffers, however, from some inconsistencies, which we discussed in [23]. To remove them, we reformulated the model by considering that the inter-pool diffusion of free TCRs takes place with rate *ϕ* proportional to the ratio of the interface to spare pool areas, and that receptors in each pool turnover between the membrane and the cytoplasm with rate *σ*, also proportional to the respective areas of the two TCR pools.

## Results

The structural identifiability and observability (SIO) of a model depend on the quantities that can be measured (output variables). The five most commonly measured quantities in T-cell activation research are *T*_*T*_ (the cell density of total TCRs); *T* (*t*) (the cell density of free TCRs); *R*(*t*) (the total amount of triggered or activated TCRs per cell, except in the *KPR-LS-IFF* model where it is the cell density of the signaling molecule P); *E*_*max*_ (the maximal effective response); and *EC*_50_ (the ligand concentration leading to a half-maximal response). The latter two quantities can be used as outputs if we know the mathematical expression that relates them to parameters and state variables. Recently, we have derived those expressions [23], and here we make use of them in the identifiability and observability analyses of the models for several output configurations. We also analyzed in each model the sensitivity of the response to changes in key parameters.

### Identifiability and Observability

When output information in complex models is limited, we can expect that a number of unknown parameters will be unidentifiable [40]. Identifiability, however, can be improved by obtaining data for additional quantities, either state variables or parameters. To take this possibility into account, we also conducted identifiability analyses assuming that several measurements are available simultaneously. Specifically, we considered the possibility of measuring *R*(*t*), *T* (*t*), *T*_*T*_, *E*_*max*_, and *EC*_50_ either alone or in some combination. Likewise, we explored the possibility that the ligand-receptor binding rate (*k*_*on*_) or the forward rate of proofreading (*k*_*p*_) are also known.

For each of the models, we determined which parameters and state variables are structurally globally identifiable (SGI), locally identifiable (SLI), and non-identifiable (SNI) for different output configurations. The results are summarized in Table 2. There the number of parameters and state variables that are SGI, SLI, and SNI are indicated. Detailed results are given in Tables S1 and S2 (Supplementary Information). Table S1 shows the identifiability results when the assumed measured outputs are *T* (*t*), *T*_*T*_ and *R*(*t*) (or a combination of *R*(*t*) with *T* (*t*) or *T*_*T*_) either alone or assuming that parameters *k*_*on*_ and *k*_*p*_ are also known. Table S2 shows the results when the outputs are *E*_*max*_ or *EC*_50_, either alone or combined with *T* (*t*), *T*_*T*_, *k*_*on*_ and *k*_*p*_. In what follows we describe the results for the different output configurations, focusing on the most relevant ones. Tables S3 and S4 rank, respectively, the models and output configurations that are more informative in terms of identifiability.

**Table 2.**
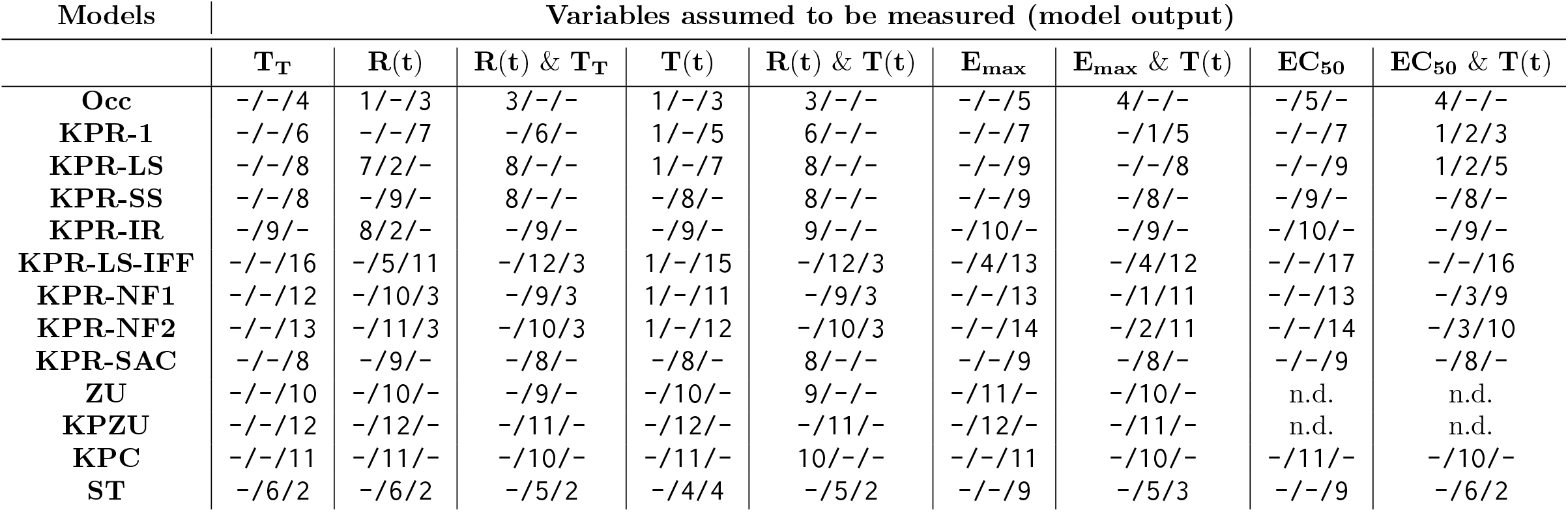
Number of parameters and initial conditions of each model that are structurally globally identifiable (SGI), locally identifiable (SLI), and non-identifiable (SNI) (indicated in each column as: #SGI/#SLI/#SNI) when the indicated specific variables are assumed to be measured; n.d., not done.

| Models | Variables assumed to be measured (model output) |  |  |  |  |  |  |  |  |
| --- | --- | --- | --- | --- | --- | --- | --- | --- | --- |
| | $T_T$ | $R(t)$ | $R(t) \& T_T$ | $T(t)$ | $R(t) \& T(t)$ | $E_{max}$ | $E_{max} \& T(t)$ | $EC_{50}$ | $EC_{50} \& T(t)$ |
| **Occ** | -/-/4 | 1/-/3 | 3/-/- | 1/-/3 | 3/-/- | -/-/5 | 4/-/- | -/5/- | 4/-/- |
| **KPR-1** | -/-/6 | -/-/7 | -/6/- | 1/-/5 | 6/-/- | -/-/7 | -/1/5 | -/-/7 | 1/2/3 |
| **KPR-LS** | -/-/8 | 7/2/- | 8/-/- | 1/-/7 | 8/-/- | -/-/9 | -/-/8 | -/-/9 | 1/2/5 |
| **KPR-SS** | -/-/8 | -/9/- | 8/-/- | -/8/- | 8/-/- | -/-/9 | -/8/- | -/9/- | -/8/- |
| **KPR-IR** | -/9/- | 8/2/- | -/9/- | -/9/- | 9/-/- | -/10/- | -/9/- | -/10/- | -/9/- |
| **KPR-LS-IFF** | -/-/16 | -/5/11 | -/12/3 | 1/-/15 | -/12/3 | -/4/13 | -/4/12 | -/-/17 | -/-/16 |
| **KPR-NF1** | -/-/12 | -/10/3 | -/9/3 | 1/-/11 | -/9/3 | -/-/13 | -/1/11 | -/-/13 | -/3/9 |
| **KPR-NF2** | -/-/13 | -/11/3 | -/10/3 | 1/-/12 | -/10/3 | -/-/14 | -/2/11 | -/-/14 | -/3/10 |
| **KPR-SAC** | -/-/8 | -/9/- | -/8/- | -/8/- | 8/-/- | -/-/9 | -/8/- | -/-/9 | -/8/- |
| **ZU** | -/-/10 | -/10/- | -/9/- | -/10/- | 9/-/- | -/11/- | -/10/- | n.d. | n.d. |
| **KPZU** | -/-/12 | -/12/- | -/11/- | -/12/- | -/11/- | -/12/- | -/11/- | n.d. | n.d. |
| **KPC** | -/-/11 | -/11/- | -/10/- | -/11/- | 10/-/- | -/-/11 | -/10/- | -/11/- | -/10/- |
| **ST** | -/6/2 | -/6/2 | -/5/2 | -/4/4 | -/5/2 | -/-/9 | -/5/3 | -/-/9 | -/6/2 |

#### (a) Identifiability when measuring the response *R*(*t*) (triggered TCRs)

If the response *R*(*t*) is the only output measured, seven of the models are fully identifiable (locally or globally), namely, *KPR-LS, KPR-SS, KPR-IR, KPR-SAC* and the three models of the *zero-ultraspecificity* family. Surprisingly, in contrast with the *KPR-LS* model, the *KPR-LS-IFF* variant has eleven unidentifiable parameters. Likewise, in the *KPR-NF1* and *KPR-NF2* models one parameter and the variables involved in the regulation of phosphatase SHP-1 activity are unidentifiable, and in the *Occ* and *ST* models some parameters and variables are also unidentifiable. Interestingly, in this case the simplest kinetic proofreading model (*KPR-1*) is not identifiable.

However, when either *T*_*T*_, the total number of TCRs, or *T* (*t*), the total number of free TCRs, is measured in addition to *R*(*t*), the *Occ* and *KPR-1* models become globally identifiable, and seven parameters in the *KPR-LS-IFF* model previously unidentifiable become identifiable. Lastly, measuring *k*_*on*_ and/or *k*_*p*_ in addition to *R*(*t*) has no positive impact on the identifiability, except in the *KPR-1* model, where all parameters become identifiable.

#### (b) Identifiability when measuring *T*_*T*_ (total number of receptors) or *T* (*t*) (instantaneous number of free receptors)

Measuring only *T*_*T*_ yields the worst results in terms of identifiability: all parameters in all models, except for the *KPR-IR* and *ST* models, are unidentifiable. Intuitively, it seems logical that *T*_*T*_ be the least informative measurement: being the sum of all TCRs, free and bound, its aggregate nature does not provide detailed information about individual variables, whose effects can be offset, thus making them indistinguishable.

However, while measuring only *T*_*T*_ is not enough for parameter identification, results for *Occ, KPR-1* and *KPR-LS-IFF* models considerably improve by combining it with *R*(*t*): many parameters that in those models were unidentifiable from *T*_*T*_ or *R*(*t*) alone become at least locally identifiable when both outputs are known (Table 2).

When *T* (*t*) is the measured output, all parameters of the *KPR-SS, KPR-IR, KPR-SAC, ZU, KPZU* and *KPC* models are identifiable, albeit only locally. In contrast, in the *Occ, KPR-1, KPR-LS KPR-LS-IFF, KPR-NF1* and *KPR-NF2* models, when only *T* (*t*) is measured *k*_*on*_ is the only identifiable parameter (globally in this case), and identifiability is not improved by knowing additionally *k*_*on*_ or *k*_*p*_; however, measuring *R*(*t*) in addition to *T* (*t*) makes all parameters identifiable in these models, except those parameters linked to the SHP-1 regulator in the *KPR-NF1* and *KPR-NF2* models, and those linked to *Y*, the activator of the signaling process, in the *KPR-LS-IFF* model. The ST model is singular in that measuring only *T* (*t*) yields four identifiable parameters (out of eight), while measuring also *R*(*t*) increases by one the number of identifiable parameters.

In general, knowing the values of *k*_*on*_ and/or *k*_*p*_ in addition to the output variables indicated in Table S1 does not improve or improve a little the identifiability of other parameters, with the important exception of the *KPR-1* model (see above).

#### (c) Identifiability when measuring *EC*_50_ and *E*_*max*_

Thus far, we have presented results for measurements that correspond directly to a single basic variable or a combination of two basic variables common to all models. The quantities *E*_*max*_ and *EC*_50_, however, are derived variables, defined by particular parameter-dependent expressions, specific to each model. For the *Occ, KPR-1* and *KPR-LS* models, these expressions can be found in the literature, as well as *E*_*max*_ for the *KPR-SS* model [11]. For the remaining models, recently, we have derived the corresponding expressions for *E*_*max*_ and *EC*_50_ (except *EC*_50_ in the *ZU* and *KPZU* models) [23] following the procedure outlined in Box 1. They are provided in the Supplementary Information.

Importantly, measuring only *EC*_50_ or *E*_*max*_ gives poor identifiability results that are only a little better than measuring only *T*_*T*_ (Table 2). Thus, when measuring only *EC*_50_ all parameters are unidentifiable in seven out of eleven models (*KPR-1, KPR-LS, KPR-LS-IFF, KPR-NF1, KPR-NF2, KPR-SAC*, and *ST*), in contrast to four models where all parameters are locally identifiable (*Occ, KPR-SS, KPR-IR*, and *KPC*). However, when *T* (*t*) is measured in addition to *EC*_50_, parameters *k*_*on*_ and *k*_*off*_, and *L*_*T*_ become identifiable in the above unidentifiable models, except in the *KPR-LS-IFF* model, where all parameters remain unidentifiable.

The case of *E*_*max*_ is even worse, resulting less informative than *EC*_50_ for all models. Thus, in this case only the *KPR-IR, ZU*, and *KPZU* models are fully locally identifiable, and in the *KPR-LS-IFF* model four out of seventeen parameters are locally identifiable. Adding measurements of *T* (*t*) considerably improves the results, so that not only the *Occ, KPR-SS, KPR-SAC* and *KPC* models become now fully identifiable, but in the *ST* model five parameters out of nine become locally identifiable, one parameter in the *KPR-1* and *KPR-NF1* models, and two parameters in the *KPR-NF2* model. All parameters in the *KPR-LS* model remain unidentifiable.

### Sensitivity analysis

To perform sensitivity analysis, we fixed a set of reference parameter values taken from the literature (Table 3) and focused on a subset of parameters usually considered more relevant for T-cell activation, namely, *k*_*off*_, *k*_*on*_, *k*_*p*_ and *L*_*T*_, common to all models—except the *ST* model that does not include *k*_*off*_ and *k*_*p*_, and the *Occ* model that does not include *k*_*p*_. We varied the values of these parameters within biologically reasonable ranges, and quantified the corresponding sensitivity of *R*(*t*). Also, given that the *KPR-NF1* and *KPR-NF2* models include a dephosphorylation mechanism—contributed by a basal rate constant *b* and a variable rate *γ P* (*t*)—, in these two models the sensitivity of *R*(*t*) was also analyzed with respect to *b* and *γ*. Moreover, since the dynamics of *P* (*t*) (active phosphatase) is key for both models’ behavior, we also analyzed the sensitivity of *P* (*t*) to parameters *k*_*off*_, *b* and *γ*. Naturally, other analyses could be performed as well; however, while it would be possible to compute the sensitivities of the outputs of all models to all parameters, many of the results would be of limited interest.

**Table 3.** Parameter values used in the sensitivity analyses.

| Name | Value | Description and reference |
| --- | --- | --- |
| $^{\dagger}k_{on}$ | $5.0 \times 10^{-5}$ | TCR-pMHC binding rate [22] |
| $^{\ddagger}k_{off}$ | $1.0 \times 10^{-2}$ | TCR-pMHC unbinding rate [22] |
| $T_T$ | $2.0 \times 10^4$ | Total number of TCRs per T cell [22] |
| $L_T$ | $1.0 \times 10^2$ | Total number of pMHC ligands per APC [22] |
| $^{\ddagger}k_p$ | $1.0 \times 10^0$ | Phosphorylation rate in KPR models [22] |
| $P_T$ | $6.0 \times 10^5$ | Total number of SHP-1 molecules per T cell [2, 36] |
| $^{\ddagger}\alpha$ | $2.0 \times 10^{-4}$ | SHP-1 activation rate [22] |
| $^{\ddagger}\beta$ | $1.0 \times 10^0$ | SHP-1 deactivation rate [22] |
| $\beta/\alpha$ | $5.0 \times 10^2$ | SHP-1 deactivation-activation ratio for KPR-NF2 [2] |
| $^{\ddagger}b$ | $4.0 \times 10^{-2}$ | Spontaneous dephosphorylation rate [24] |
| $^{\ddagger}\gamma$ | $4.4 \times 10^{-4}$ | Dephosphorylation rate by SHP-1 [2] |
| $^{\ddagger}\phi$ | $9.0 \times 10^{-2}$ | Phosphorylation rate of $C_N$ to $C_{N+1}$ [11] |
| $^{\ddagger}\lambda$ | $1.0 \times 10^5$ | Decay rate of phosphorylated free TCRs [27] |
| $^{\ddagger}\rho$ | $1.0 \times 10^3$ | TCR-pMHC rebinding rate [27] |
| $r$ | $1.5 \times 10^0$ | Factor in dephosphorylation rate [59] |
| $r_p$ | $1.03 \times 10^0$ | Factor in phosphorylation rate [59] |
| $Y_T$ | $1.0 \times 10^2$ | Total number of Y molecules per T cell [12] |
| $Z_T$ | $2.5 \times 10^0$ | Total number of Z molecules per T cell [12] |
| $^{\dagger}\sigma$ | $1.0 \times 10^2$ | $C_N$-dependent Y activation rate [12] |
| $^{\dagger}\delta$ | $1.0 \times 10^2$ | Y-dependent Z activation rate [12] |
| $^{\dagger}\mu$ | $5.0 \times 10^2$ | $C_N$-dependent Z deactivation rate [12] |
| $^{\ddagger}\gamma_+^y$ | $1.0 \times 10^0$ | Basal Y activation rate [12] |
| $^{\ddagger}\gamma_-^y$ | $5.0 \times 10^2$ | Basal Y deactivation rate [12] |
| $^{\ddagger}\gamma_+^z$ | $1.0 \times 10^0$ | Basal Z activation rate [12] |
| $^{\ddagger}\gamma_-^z$ | $5.0 \times 10^2$ | Basal Z deactivation rate [12] |
| $^{\dagger}k_+$ | $1.0 \times 10^0$ | Association rate constant [28] |
| $^{\ddagger}k_-$ | $1.0 \times 10^1$ | Dissociation rate constant [28] |

$^{\dagger}$ Units are $s^{-1}(\text{molecules/cell})^{-1}$ . $^{\ddagger}$ Units are $s^{-1}$ .

Positive sensitivity values indicate a direct relationship between a state variable and a parameter, while negative values indicate an inverse relationship. However, for the purpose of this work what is relevant is how intense is the sensitivity. Therefore, we calculated in all cases the absolute value of relative sensitivities.

#### (a) Sensitivity of *R*(*t*) to the unbinding rate *k*_*off*_

We calculated the influence of *k*_*off*_ on the intensity of the response, *R*(*t*), along a time interval and for a wide range of *k*_*off*_ values (Figure 2). The time interval was chosen specifically for each model to better capture the sensitivity variations along the range of *k*_*off*_ values. This provides a *contour map* that allows to pinpoint the *k*_*off*_ values and times at which sensitivity is highest. The model where *R*(*t*) is less sensitive to *k*_*off*_ is *ZU*, with a maximum sensitivity in the order of 10^*−*3^. Models *KPR-IR* and *KPR-LS-IFF* are next, with a maximum sensitivity ≈ 0.5. On the other hand, the *KPR-LS, KPR-SS, KPR-SAC*, and *KPZU* models exhibit the highest sensitivity, with a maximum that is 4-5 times higher than that in the previously mentioned models. Nevertheless, those sensitivities are still moderate. Also, we notice that in the negative feedback models (*KPR-NF1* and *KPR-NF2*) the highest values of the sensitivity of *R*(*t*) to *k*_*off*_ are restricted to a very small range of values of that parameter.

**Fig 2.**
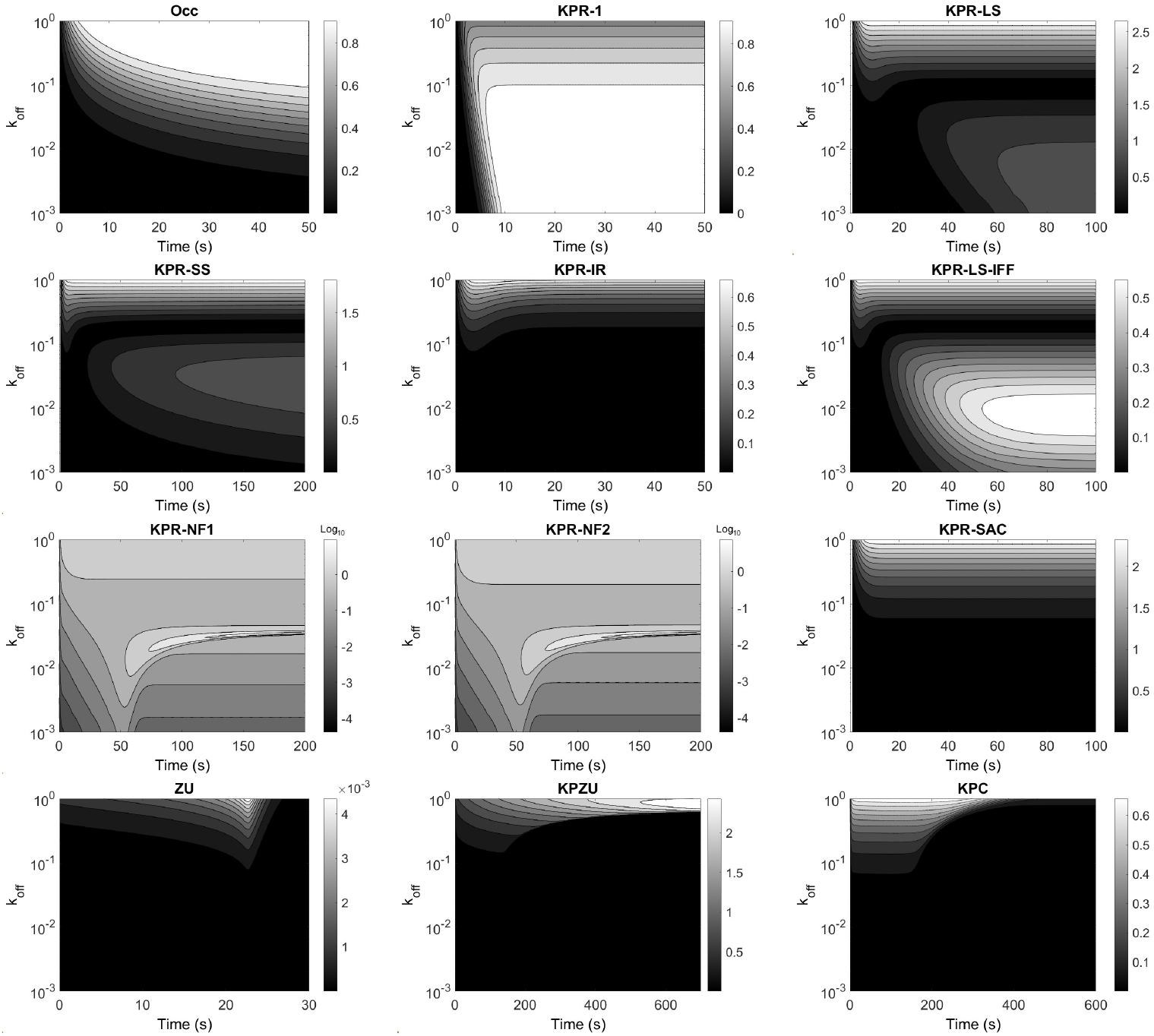
Contour map of the sensitivity of the response *R*(*t*) to variations in the unbinding rate *k*_*off*_ in each model. Note that the sensitivity gradient bars at the right of each panel are in arithmetic scale, except in the panels corresponding to the *KPR-NF1* and *KPR-NF2* models, where they are in base-10 logarithmic scale. The *ST* model is not included because *k*_*off*_ is not a parameter in this model.

In general, the sensitivity of *R*(*t*) to *k*_*off*_ strongly depends on the specific values of *k*_*off*_ and less on the specific time, except in the *KPR-LS-IFF* and *KPZU* models, where it also depends strongly on the specific time.

#### (b) Sensitivity of *R*(*t*) to the binding rate *k*_*on*_

Next, we analyzed the sensitivity of the response to the TCR-pMHC binding rate. Interestingly, all models follow a similar trend: the sensitivity is low (*KPZU, KPC* and *ST* models) to very low (rest of the models). The results are summarized in Figure S4 (Supplementary Information). In the models with very low sensitivity, the maximum values ranged from 0.07 to 0.53 and were attained only during the first few seconds of simulation and for very low *k*_*on*_ values.

The sensitivity results for the *KPZU* and *KPC* models differ significantly from the other models, with the following three features standing out: (i) within the lower values of the analyzed *k*_*on*_ range the response is moderately sensitive to changes in *k*_*on*_ for most or all of the simulation time; (ii) the sensitivity slightly increases over time; and (iii) at high parameter values, changes in *k*_*on*_ has minimal impact on the response at all simulated times. Finally, in the *ST* model, the sensitivity of *R*(*t*) to *k*_*eff*_ is qualitatively similar to most models, but for low values of *k*_*eff*_ the sensitivity is highest at all plotted times, and for the first ten seconds the sensitivity is highest for the whole plotted range of *k*_*eff*_.

#### (c) Sensitivity of *R*(*t*) to the phosphorylation rate *k*_*p*_

The sensitivity of the response to the phosphorylation forward rate, *k*_*p*_, follows a pattern qualitatively similar to that of the sensitivity to *k*_*on*_, but with the highest sensitivity attained at all times and, as a general rule, for higher values of *k*_*p*_ compared to *k*_*on*_. The results are summarized in Fig. S5. Notably, for the *KPR-NF1* and *NF2* models a very short range is observed, within which the response becomes highly sensitive to variations in *k*_*p*_; in other words, there is a critical value of *k*_*p*_ ≈ (*k*_*p*_ 1.2 *s*^*−*1^) that not only maximizes its impact on the response, but that impact is 50- to 60-fold higher than the highest sensitivity to any of the other parameters. Below this critical value, the sensitivity to *k*_*p*_ becomes moderate at all times. Another interesting case is the *KPR-LS-IFF* model, whose response is maximally sensitive to *k*_*p*_ within two distinct ranges of this parameter, separated by a wide and deep basin. Finally, the case of the *KPZU* model is qualitatively like that of the *KPR-NF1* and *NF2* models, but with a wider central range, where the sensitivity is highest at *k*_*p*_ ≈ 1.25, and with such sensitivity being moderate.

#### (d) Sensitivity of *R*(*t*) to total ligand *L*_*T*_

With respect to the sensitivity of the response to the total ligand or antigen dose, *L*_*T*_, the models *Occ, KPR-1, KPR-LS, KPR-SS, KPR-IR, KPR-SAC*, and *ST* follow a quite similar pattern and have a maximum sensitivity ≈ 1, except in the *ST* model, where it is ≈ 4. In all these models, except *KPR-SS*, at any given ligand density higher than 10^3^ molecules/cell, the sensitivity decreases with time, initially fast and then gradually slower. At low ligand densities, the sensitivity is not only highest but independent of time during all the computed time interval (Fig. S6). In the *KPR-SS* model, the sensitivity is nearly constant in time at any given ligand density. In these seven models, at any time after the first two seconds, the sensitivity increases steadily from near zero to a maximum value for decreasing ligand density (Fig. S6). The *KPR-LS-IFF* model differs substantially from the above models in that the sensitivity of *R*(*t*) to *L*_*T*_ is biphasic at any given time after the first few seconds, exhibiting a maximum at intermediate ligand densities (200-500 molecules/cell) that decreases with time. The *KPC* model displays a similar biphasic behavior, but here the sensitivities are largely independent of time, and the maximum sensitivity is attained at *L*_*T*_ ≈ 100 molecules/cell. The *KPR-NF1* and *NF2* models have also a biphasic behavior of the sensitivity of *R*(*t*) to *L*_*T*_, but only within a time window from about 20 *s* to 70 *s* and a very small range of ligand density, located near the highest values of *L*_*T*_. Finally, the *ZU* and *KPZU* models display a very low sensitivity for *L*_*T*_ *<* 10 molecules/cell at all computed time. On the other hand, in the *ZU* model, the sensitivity increases for decreasing values of *L*_*T*_ and for increasing time. In contrast, the *KPZU* model displays a biphasic behavior with a maximum sensitivity at *L*_*T*_ ≈ 5 molecules/cell that increases with time.

The sensitivities of *R*(*t*) to parameters *k*_*off*_, *k*_*on*_, *k*_*p*_ and *L*_*T*_ are summarized as histograms in Fig. 3. For each model, histograms were obtained from data at a time when maximum sensitivity is attained for some value of the concerned parameter. At other times the sensitivities were either similar but lower, or much lower.

**Fig 3.**
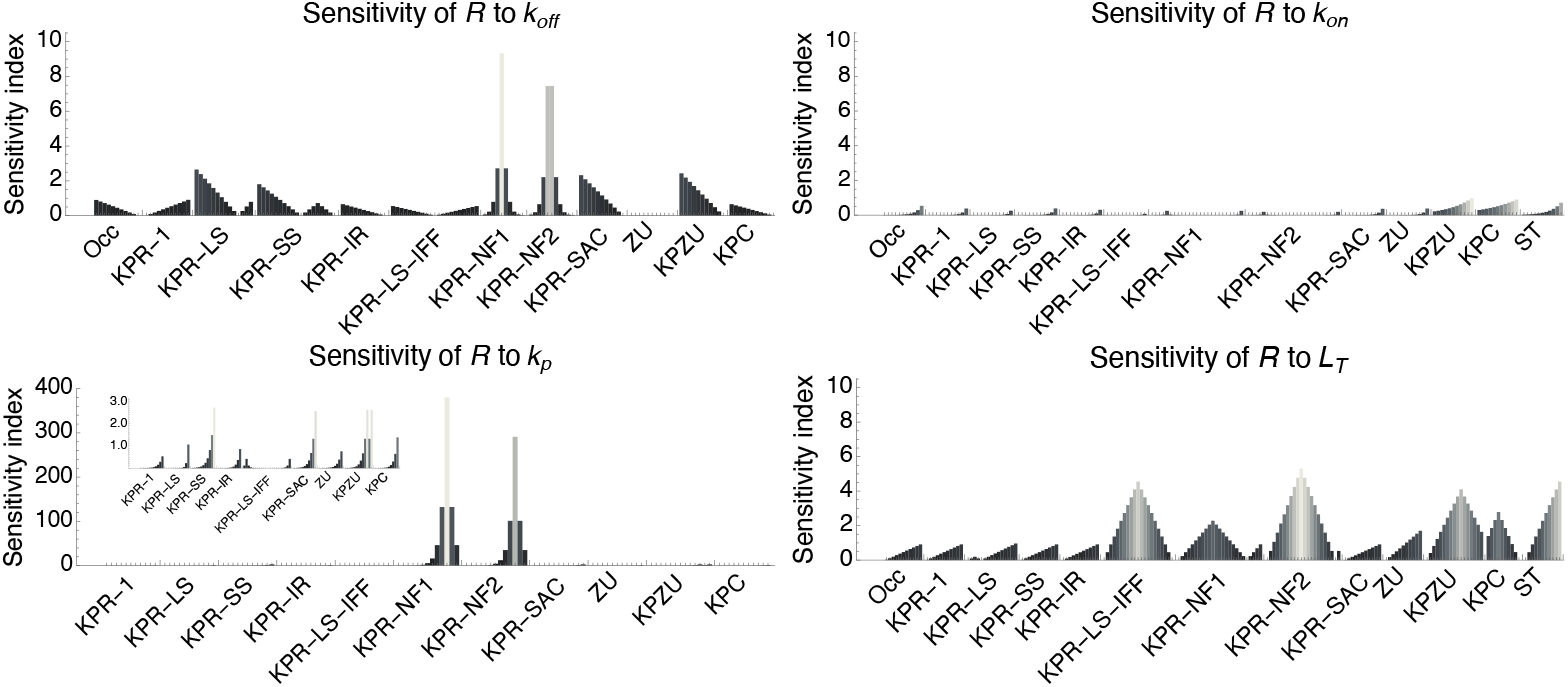
Summary of the sensitivity of the response *R*(*t*) to the three main kinetic parameters, *k*_*off*_, *k*_*on*_ and *k*_*p*_, and to the ligand density per cell, *L*_*T*_, in the different models. Data was taken from the graphs in Fig. 2, and Figs. S4 to S6. For each graph, a corresponding histogram was obtained using the sensitivity values at a particular time at which maximum sensitivity is attained. In each histogram the bars from left to right correspond to the sensitivity values from the maximum parameter value (top of the graph) to the minimum parameter value (bottom of the graph) at that particular time. Note the very high sensitivity of *R*(*t*) to *k*_*p*_ in models KPR-NF1 and KPR-NF2 compared to the other parameters. In that graph an inset is included with a scale appropriate for visualizing the sensitivity in the other models.

#### (e) Sensitivity of *R*(*t*) to *b* (spontaneous dephosphorylation rate constant for *C*_*i*_ complexes), *γ* (rate constant for SHP-1-dependent dephosphorylation of *C*_*i*_ complexes) and *P*_*T*_ (total SHP-1 molecules)

The *KPR-NF1* and *KPR-NF2* models introduce a passive or spontaneous dephosphorylation of *C*_*i*_ complexes to a *C*_*i−*1_ state, and an active dephosphorylation mediated by active *SHP-1* phosphatase. Disclosing the impact that the parameters involved in such dephosphorylation process have on the response could be valuable for explaining how SHP-1 cell level impacts the efficiency of antigen discrimination, which is an open question [24]. To shed light on this aspect, we performed additional sensitivity analyses for these two models. Two parameters, *b* and *γ*, and one state variable, *P* (*t*) (active *SHP-1*), determine the *C*_*i*_ dephosphorylation rate; in addition, for fixed rates of the dynamics of *SHP-1* in those models, the level of *P* (*t*) is determined by *P*_*T*_, the total cellular amount of *SHP-1*. Hence, we analyzed the sensitivity of *R*(*t*) to parameters *b, γ* and *P*_*T*_. Results are shown in Fig. S7(a), left and middle panels (Supplementary Information).

In spite of the different definitions of *R*(*t*) in *KPR-NF1 KPR-NF2*, the sensitivities are very similar in both models. Specifically, *R*(*t*) has a quite modest sensitivity to *γ*, with maximum values of 2.72 for *KPR-NF1* and 2.14 for *KPR-NF2* (Fig. S7(b)). In contrast, the maximum sensitivities to *b* and *P*_*T*_ are, respectively, approximately 10- and 300-fold higher than that to *γ* (Fig. S7(b)). However, such maximum sensitivities to *b* and *P*_*T*_ are attained only within a very narrow range of parameters’ values (Fig. S7(a)), close to their reference values (Table 3).

#### (f) Sensitivity of *P* (*t*) (active SHP-1 level) to *k*_*off*_, *b* and *γ*

Given the high impact of *k*_*p*_ on the response (Fig. 3), the direct effect of *C*_1_ levels on the generation of active *SHP-1*, and the direct effect of *b, γ* and *P* (*t*) on the generation of *C*_1_ from *C*_2_ (Fig. S1(g)), we hypothesized that *b* and *γ* could also have an important impact on the levels of *P* (*t*). Since *k*_*off*_ has also a direct effect on the levels of *C*_1_, we considered that it could also have an important impact on the levels of *P* (*t*). Hence, we investigated the sensitivity of *P* (*t*) to *k*_*off*_, *b* and *γ*. Because the definition of *P* (*t*) is the same in both models, the results apply to both of them. For these three parameters the sensitivity has a triphasic behavior after the first 50 seconds (Fig. S7(a) right panels). Thus, for a fixed time, and going from highest to lowest parameter values, the sensitivity first decreases, then increases until a maximum, and finally decreases again. The maximum sensitivity to *k*_*off*_ and *γ* are moderately high, respectively, 6.48 and 5.05 (Fig. S7(b) right panel). However, the maximum sensitivity of *P* (*t*) to *b* is quite high, about 6-fold higher than that to *k*_*off*_ and *γ*.

## Discussion

Identifiability analysis is being increasingly used in theoretical biology, particularly in biochemical networks and systems biology, to assess whether a mathematical model is well formulated to provide reliable predictions [15–19, 41]. However, in the immunological field modeling works have rarely analyzed parameter identifiability (*e*.*g*., [42–44]) and models of basic immune processes, such as lymphocyte activation or the germinal center reaction, to the best of our knowledge have not yet been analyzed in this respect. In contrast, sensitivity analysis is being increasingly performed also in the study of models of basic immune processes (*e*.*g*., [20, 45]). In the present study we have analyzed thirteen *phenotypic* models of T-cell activation from the point of view of parameter identifiability and state variables’ sensitivity to parameters.

### Identifiability of T-cell Activation Models

Identifiability analysis assumes a given known or measured output. We considered here the following *known output* configurations: (1) five individual output variables (*T* (*t*), *T*_*T*_, *R*(*t*), *E*_*max*_ and *EC*_50_), (2) a combination of two individual output variables (*e*.*g*., *R*(*t*) + *T* (*t*)), and (3) a combination of one or two output variables with parameters *k*_*on*_ and/or *k*_*p*_ (*e*.*g*., *R*(*t*) + *T* (*t*) + *k*_*on*_ + *k*_*p*_). Surprisingly, with the only important exception of the *KPR-1* model, where knowing *k*_*on*_ or *k*_*p*_ in addition to *R*(*t*) makes all parameters change from non-identifiable to identifiable, in general knowing *k*_*on*_ and *k*_*p*_ in addition to those output variables does not improve, or improves only marginally, the identifiability of the analyzed models. Consequently, the identifiability analysis was focused on the nine different output configurations indicated in Table 2.

We found that three models (*KPR-LS-IFF, KPR-NF1* and *KPR-NF2*) are unidentifiable under any of the considered output configurations. Of the remaining models, nine are identifiable under output configurations *R*(*t*) + *T* (*t*) and *R*(*t*) + *T*_*T*_, seven models under *R*(*t*) and *E*_*max*_ + *T* (*t*), and six under *T* (*t*). Other output configurations considerably reduce the number of identifiable models (summarized in Table S4). An interesting realization is that knowledge of only the total number of receptors (*T*_*T*_) provides no information for parameter identification, except in the *KPR-IR* and *ST* models. Likewise, the quantities *E*_*max*_ and *EC*_50_ are the next least informative measurements for parameter identification, with, respectively, three and four models being identifiable. In contrast, measuring only the response (*R*(*t*)) or the number of free TCRs per cell over time (*T* (*t*)) is enough to identify, respectively, up to seven or six models.

Lever *et al*. systematically derived and analyzed many models of T-cell activation. Their results suggested that the *KPR-LS-IFF* model is the one that best captures all phenotypic traits analyzed by them [12]. However, our identifiability analysis indicates that that model is non-identifiable under any feasible output configuration. More specifically, parameter δ and the values *Y* (0) and *Y*_*T*_ of the trigger of T-cell activation remain non-identifiable under all output configurations. This, together with the fact that *Y* is a hypothetical metabolite, hampers the reliability of this model for predictive purposes. A similar case is that of the *KPR-NF1* and *KPR-NF2* models. The *KPR-NF1* model was proposed by Altan-Bonnet *et al*. to account for the observed important impact of *SHP-1* on T-cell activation [24, 36]. Several years later a further modification was proposed, named here the *KPR-NF2* model, to account for recent observations indicating that the number of available ITAMs in TCRs has an important impact on the kinetics of the T-cell response [2]. However, similarly to the *KPR-LS-IFF* model, our present analysis indicates that the parameter *γ* and the values of *P* (0) and *P*_*T*_ that define the negative feedback modulus are non-identifiable under any output configuration examined (Table S1), thus affecting the reliability of these models in practice. Nevertheless, in this case this could be remedied by measuring additionally the parameter *γ* as well as *P*_*T*_ and *P* (0) (the amount of active *SHP-1* in resting conditions).

Besides the merits of the different models in coping with the three key features of sensitivity, specificity and reaction speed, the question of which of the proposed models is better to reliably predict the response of T cells to antigen must be answered considering identifiability and observability issues. For instance, if both the number of individual outputs and the total number of output configurations that make a model identifiable are taken as a measure of how robustly a model is defined, our results indicate that the *KPR-IR* model is better than others, because it is identifiable not only under all combined output configurations but also under all individual outputs (Tables 2 and S3). The *KPR-IR* model was originally proposed [27] to enhance the poor sensitivity of the *KPR-1* model while retaining specificity; to this end it requires a high number of proofreading steps (*N* ≥ 25), otherwise the kinetics of the response would be very similar to that of the *KPR-1* model [27, 46]. Here we analyzed a simpler *KPR-IR* version [34] for which, unlike the *KPR-1* model, all parameters are at least SLI under all output configurations, and SGI when *R*(*t*) (alone or together with *T* (*t*) or *T*_*T*_) or *T* (*t*) together with *k*_*p*_ are known, supporting the relevance of this model. The next best models are *KPR-SS* and *KPC* with seven output configurations that make them identifiable, followed by the *KPR-SAC, ZU* and *KPZU* models (summarized in Table S3).

### Sensitivity in T-cell Activation Models

The combination of sensitivity and identifiability analyses is a powerful tool to determine model parameters that could be used to control or fine tune a T cell response. There are four potential scenarios that stand out: (1) the response *R*(*t*) has moderate sensitivity to a parameter that is identifiable; then, that parameter is a good candidate for controlling the response; (2) the sensitivity of *R*(*t*) to a parameter is low or very low within a range of biologically relevant values; then, even if the parameter is identifiable, its specific value will affect very little the predicted output and hence that parameter cannot be used to control it; (3) the sensitivity of *R*(*t*) to a parameter is high or very high; in this case even small changes in its value will greatly impact the response, making that parameter useless to control it; and (4) a model has some non-identifiable parameters under all output configurations, but the sensitivity of the response to those parameters is very low; then, that model can still be used to make reliable predictions.

The sensitivity analyses allowed us to identify in each model which of the four main parameters, that is, the unbinding rate, *k*_*off*_, the binding rate, *k*_*on*_, the phosphorylation rate, *k*_*p*_ and total ligand, *L*_*T*_, have a moderate influence on the response over time (*R*(*t*)) such that variations in their values can lead to comparable changes in the response. If we focus on the three main kinetic parameters that are common to most T-cell activation models (i.e., *k*_*off*_, *k*_*on*_ and *k*_*p*_), our results clearly show that *k*_*off*_ has quite moderate impact on the response in most models. With respect to the phosphorylation rate *k*_*p*_, with the exception of the *KPR-NF1* and *KPR-NF2* models (where the maximum sensitivity of *R*(*t*) to *k*_*p*_ is very high, but only within a very narrow range of *k*_*p*_ values), this parameter also has a moderate effect on the response (*R*(*t*)) in the other eleven models and at almost any time step. We investigated further what could be special in the *KPR-NF1* and *KPR-NF2* models such that in them *R*(*t*) has so high sensitivity to *k*_*p*_. We found that of the parameters involved in the negative feedback modulus only the level of total *SHP-1* (*P*_*T*_) has also a very high impact on *R*(*t*). Thus, all the three main parameters affecting the level of the complex *C*_1_ (the activator of *SHP-1*) have the greatest impact on *R*(*t*). As for the sensitivity to *k*_*on*_, there is a remarkable difference between the moderate maximum sensitivity in the *KPZU, KPC* and *ST* models and the low maximum sensitivities in the other models. In other words, *k*_*on*_ has in general a very limited impact on *R*(*t*). Interestingly, in the *KPZU* and *KPC* models the sensitivity of the response to *k*_*off*_ and *k*_*on*_ is somehow opposite to each other, thus, while it is very low to *k*_*off*_ within a large parameter range, it is moderate to *k*_*on*_ also within a large parameter range. Finally, we have shown here that the sensitivity of *R*(*t*) to the total ligand per APC, *L*_*T*_, is moderate in all models (Fig. 3).

In summary, in most models the response may be best regulated by tuning *k*_*off*_, *k*_*p*_ and *L*_*T*_ provided they remain within a range not too far from the maximum sensitivity of *R*(*t*) to them (Fig. S5). Interestingly, that range is large in nearly all models. In contrast, in general, except the *KPZU, KPC* and *KPR-ST* ones, *k*_*on*_ is not a good candidate to control the response because the sensitivity of *R*(*t*) to it is very low. Finally, in the case of the *KPR-IR* model —the one favored by identifiability analysis— the maximum sensitivities of *R*(*t*) to *k*_*off*_, *k*_*on*_, *k*_*p*_ and *L*_*T*_ are, respectively, 0.66, 0.31, 0.86 and 0.91. This suggests that in this model the parameters *k*_*p*_ and *L*_*T*_ are particularly well suited to tune the response.

### Conclusion

The older phenotypic models (*Occ* and *KPR-1*) could only account for a limited subset of observed T-cell phenotypes. Variants of those models were then developed to incorporate an increasing number of those phenotypes. As shown here, some of them are identifiable when *R*(*t*) or *T* (*t*) are the only known output. Further modifications of these identifiable models, devised to more comprehensibly capture the diverse T-cell phenotypes, would likely exhibit identifiability issues. However, such issues should not lead to their automatic rejection: as we have found, some of the non-identifiable models can become identifiable by measuring a second output; moreover, as indicated above for the *KPR-NF1* and *KPR-NF2* models, structural identifiability issues of T-cell activation models can be remedied in some cases by measuring still more outputs. When this solution is not experimentally feasible or convenient, an alternative may be to simplify or reparametrize the equations of the model [17]. Lastly, we note that the findings reported here have been obtained with a structural identifiability approach that implicitly assumes the availability of noiseless data, collected at as many time points as necessary. This is of course an ideal setting; real experimental setups introduce additional limitations, which can be accounted for with practical identifiability techniques [47].

Any immune system deficiency in discriminating between tumor- or pathogen-derived Ags and normal self-protein Ags can lead to pathologies such as cancer or persistent viral infections (tolerance or low response level) or autoimmunity (excess reactivity). Having a model of T lymphocyte activation with reliable predictive capacity in different experimental situations would be instrumental in the development of theoretically based methods to control the immune response and make it a more efficient process in the face of pathological threats. Incorporating identifiability and sensitivity analysis in the study of models of fundamental immune processes should greatly enhance the values of those models and the immunological insights that can be gained from them.

## Materials and Methods

### Modeling framework and notation

We consider here deterministic models described by ordinary differential equations (ODE) of the form:

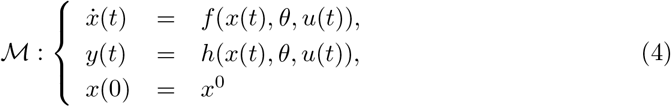

where *x*(*t*) = (*x*_1_(*t*),…, *x*_*n*_(*t*)) is the state variables vector, *u*(*t*) = (*u*_1_(*t*),…, *u*_*q*_(*t*)) is the input variables vector (usually, in biological models there are no input variables), *θ* = (*θ*_1_,…, *θ*_*p*_) is the parameters vector, *x*(0) = (*x*_1_(0),…, *x*_*n*_(0)) is the initial conditions vector, and *y*(*t*) = (*y*_1_(*t*),…, *y*_*m*_(*t*)) is the output vector. Since this last vector corresponds to experimentally measurable functions, it is usually, but not always, a subset of the state variables. All the components of the four vectors take real values. The functions *f* (*x*(*t*), *θ, u*(*t*)) and *h*(*x*(*t*), *θ, u*(*t*)) considered in this work are generally non-linear, and contain rational terms. The output is known, while the parameters (or a subset of them) are typically unknown. To simplify the notation we sometimes omit the dependence on *t*.

### Structural identifiability and observability analysis

In order to ascertain whether model parameters can be uniquely determined from the outputs, i.e., from experimentally measured quantities, we perform a *Structural identifiability* analysis (SIA) [40]. A parameter with an infinite number of values compatible with the experimental data is structurally unidentifiable, as its true numerical value cannot be determined. Conversely, a parameter is globally identifiable if there is only one possible solution in the parameter space, and locally identifiable if there are a finite number of solutions.

*Observability* is a property that informs about the possibility of inferring the internal state of a system at time *t* by observing its output at times *τ* such that *t* ≤ *τ*. Notice that, by considering the unmeasured initial conditions *x*^0^ as parameters, their observability can be studied as structural identifiability, *i*.*e*., identifiability of an initial condition *x*^0^ is equivalent to observability of *x*_*i*_(*t*) [19]. Thus, we assessed the observability of state variables *x*(*t*) by analysing the identifiability of their initial conditions *x*^0^.

Structural identifiability was assessed using a number of methods. Whenever possible, we followed a differential algebra approach that can be applied to systems defined by rational ODE functions, and involves deriving algebraic equations that connect the model parameters with the inputs and outputs [48]. Briefly, model equations (4) are transformed into a set of polynomial differential equations that depend solely on the output variables *y* [49], preserving the dynamics of the model output while removing the state variables. Extracting from these equations the coefficients that multiply the same variables in terms of *y* and grouping them into the same equation, we obtain the so-called exhaustive summary. Then, the identifiability of model parameters can be determined by evaluating whether the mapping defined by the exhaustive summary is unique [50]. The differential algebra approach can distinguish between global and local identifiability. To apply it we used SIAN (Structural Identifiability ANalyser), an open-source tool that integrates differential algebra with a Taylor series-based method [51, 52]. This choice was motivated by SIAN’s computational efficiency, reliability, and its ability to include the initial conditions of state variables, *x*^0^, as parameters in the analysis [34]. In those cases where the analysis could not be performed with SIAN due to non-rationalities, we used alternative methods that are more generally applicable, but can only determine local identifiability: STRIKE-GOLDD [53] and StrucID [54].

#### Box 1

Outline of the procedure for calculating *E*_*max*_ and *EC*_50_ in T-cell activation ODE models.

*Step 1: Conservation equations for ligands and receptors:* We assume that the total amount of pMHC and TCR does not change during the T-cell activation process. Therefore, denoting *L*_*T*_ = total cognate pMHC; *L* = total unbound cognate pMHC; *T*_*T*_ = total TCR; *T* = total unbound TCR; and *C*_*T*_ = total TCR-pMHC complex in the system, the following conservation equations hold:

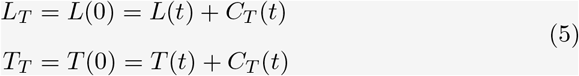

*Step 2: Calculate* 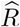, *the response level R*(*t*) *at steady state, as a function of L*_*T*_, *T*_*T*_ *and θ*.

*Step 3: Calculate E*_*max*_ *as* 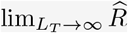. In mathematical terms, ligand saturation is expressed as *L*_*T*_ → ∞. Under these conditions, most model systems reach their maximal efficacy, allowing the response to be evaluated at its upper limit.

*Step 4: Calculate EC*_50_. The ligand potency or *EC*_50_ is defined as the value of the total amount of ligands, *L*_*T*_, at which 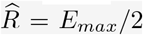. Replacing 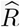 and *E*_*max*_ with their expressions (obtained in steps 2 and 3) yields, in most cases, an expression for *L*_*T*_ in terms of *T*_*T*_ and *θ*.

### Output variables

Structural identifiability and observability depend on the output definition. Hence, in our analyses we considered different possible output configurations for each model, including, as mentioned above, the total amount of TCRs per cell (*T*_*T*_), the total amount of free TCRs per cell (the state variable *T*), the total amount of triggered or activated TCRs per cell (which is *R*(*t*) except in one model, see below), the maximum achievable T-cell activation level or maximum *R*(*t*) (defined in most of the considered models as *E*_*max*_ = *R*(*t*) *at steady state and saturating conditions of pMHC*), and the amount of pMHC per APC that results in 50% of *E*_*max*_ (*EC*_50_).

In each model, the first two equations of the ODE system govern the dynamics of free receptors and ligands. Therefore, the outputs *T* and *T*_*T*_ are common to all models. However, the definition of *R*(*t*) is not the same in all models. Thus, while in most phenotypic models *R*(*t*) = *amount of TCRs in ligand-receptor complexes that reach the signaling state* per T cell, in some models (*KPR-SS* and *KPR-IR*) *R*(*t*) also includes the amount of recently unbound TCRs that remain in the signaling state. This complicates considerably the calculation of *E*_*max*_ and *EC*_50_ in several of the models. In Box 1 we outline the main steps we followed to derive the equations for *E*_*max*_ and *EC*_50_ in most of the models. A detailed description of the derivation of *E*_*max*_ and *EC*_50_ for each phenotypic model is provided in [23].

### Sensitivity analysis

We apply sensitivity analysis to quantify the extent to which a model prediction of T-cell activation depends on each parameter. More specifically, we perform a local sensitivity analysis to calculate quantitative changes in model outputs relative to parameter variations around a local point in the parameter space [55]. For that, we first compute at each time point *t*_*k*_ (*k* = 1, 2,…, *N*) the absolute sensitivity coefficient of output variable *y*_*i*_ to parameter *θ*_*j*_, defined as [56, 57]:

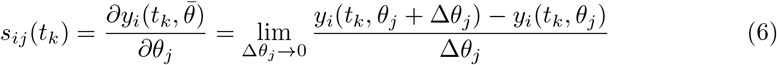

where 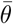 is the nominal parameter vector. Then, in order to compare the sensitivities to the different parameters of all output values, we normalize them, that is, we obtain the relative sensitivities [21]:

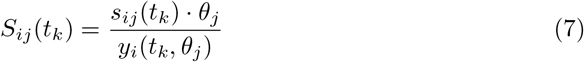

Absolute sensitivities were calculated following the method presented in [58], which approximates the gradients from (6) by extending the output function to the complex space (see Supplementary Information, subsection 1.2). We used our own Matlab implementation of the code described in [58], adjusting the presentation of the results to produce plots that better capture how the impact on the response evolves with time. Since the relevant time interval may differ for each model, we explored a sufficiently broad range to capture the model dynamics before reaching the steady state. The code for these calculations is available on GitHub (https://github.com/Xabo-RB/Inmunology-analysis.git).

Unlike structural identifiability analysis, which can be performed symbolically, sensitivity analysis entails integrating ordinary differential equations, which requires assigning numerical values to the parameters. The values used in our analyses are shown in Table 3. In this table we indicate, for each parameter, the value that we used and the reference from which that value was taken.

## Supporting information

**S1 File. Supplementary Information**. Text file that provides theoretical background of the methods applied in this paper, the equations and diagrams of all models, and supplementary results.

