## Supplementary Text 1 for "Leveraging mathematical models to predict and control T-cell activation"

### Supplementary information of the paper "Leveraging mathematical models to predict and control T-cell activation"

**Contents:**

1 Methods:

1.1 Structural identifiability analysis with differential algebra. Basic theoretical background

1.2 Sensitivity analysis with complex perturbation. Basic theoretical background

2 Models: equations and diagrams

3 Supplementary results:

3.1 Identifiability analysis

3.2 Sensitivity analysis

4 Bibliography

### 1 Methods

#### 1.1 Structural identifiability analysis with differential algebra. Basic theoretical background.

Differential algebra focuses on identifying algebraic equations that relate model parameters to inputs and outputs. This approach can handle systems of rational equations; however, it generally cannot be used to analyze the global structural identifiability of non-rational models. Let us consider, as in Eqn. [4] of the main text of the paper, a model of the form:

$$\mathcal{M}: \begin{cases} \dot{x}(t) &= f(x(t), \theta, u(t)), \\ y(t) &= h(x(t), \theta, u(t)), \\ x(0) &= x^0 \end{cases} \tag{1}$$

For this system, we denote the input-output variable map, given an initial state $x^0$, as:

$$y = \Phi(x^0, \theta, u) \tag{2}$$

The number of model solutions is defined by this mapping. A model is locally identifiable if the input-output map leads to a finite number of solutions for the model parameters. Formally, a model is locally identifiable if there is a finite number of parameter sets compatible with a given set of output observations.

***Definition* 1.1.** *Structurally locally identifiable model. Consider a model defined by Eqn. 1, with parametric space $\Theta$ and a parameter vector $\tilde{\theta}$. The model is locally identifiable in $\tilde{\theta}$ if there is an open neighborhood $\Theta_0$ of $\tilde{\theta}$ in $\Theta$ such that for all initial conditions $x^0 \in \mathbb{R}^n$, there is a unique parameter vector $\hat{\theta}$ in $\Theta_0$ that satisfies the following equation:*

$$\Phi(x^0, \hat{\theta}, u) = \Phi(x^0, \tilde{\theta}, u) \tag{3}$$

This definition implies that it is possible to uniquely determine the parameter vector within an open neighborhood of a point in the parameter space. It can also be proven that, for models given by rational functions, there is a finite number of solutions, each of which is isolated in different open sets of the space.

Similarly, a globally identifiable model is one in which the parameter vector can be unambiguously computed across the entire parameter space. In contrast, an unidentifiable model contains infinitely many points in parameter space where the input-output equation does not provide a unique solution for the parameter. For a more detailed account see [1].

##### 1.2 Sensitivity analysis with complex perturbation. Basic theoretical background.

Equation [6] in the main text computes the absolute sensitivity coefficients with respect to parameter $\theta_j$ for each output function $y_i$ at time point $t_k$. If we consider the coefficients for all components in vector $\theta \in \mathbb{R}^p$, for all output functions in vector $y \in \mathbb{R}^m$, and for all time points $(t_1, \ldots, t_n)$, and we denote with $\bar{\theta}$ the vector of nominal parameter values, we get the so-called *sensitivity matrix*:

$$\mathbf{S}_{(m \times n) \times p} = \begin{bmatrix} s_{11}(t_1) & \cdots & s_{1p}(t_1) \\ \vdots & \ddots & \vdots \\ s_{m1}(t_1) & \cdots & s_{mp}(t_1) \\ \vdots & \ddots & \vdots \\ s_{11}(t_n) & \cdots & s_{1p}(t_n) \\ \vdots & \ddots & \vdots \\ s_{m1}(t_n) & \cdots & s_{mp}(t_n) \end{bmatrix}. \tag{4}$$

where:

$$s_{ij}(t_k) = \frac{\partial y_i(t_k, \bar{\theta})}{\partial \theta_j} \tag{5}$$

In order to apply the complex perturbation method, the ODE system must be defined by analytic functions. A system of ordinary differential equations expressed in the form of Eqn. 1 is analytic at a point if it can be expressed as a convergent power series around that point. The system's time trajectory, $x(t, \theta)$, is also analytic in $\theta$ if $f(x(t), \theta)$ is analytic since the solution inherits the analytic properties of the function defining the ODE. The derivative of any real analytic function with respect to the parameter $\theta_j$ can be computed as an approximation based on complex values, in order to avoid rounding problems and increase accuracy [2]. The method for computing this derivative can be briefly described as follows:

**Sensitivity analysis with complex perturbation**

*Step 1: Directional derivative.* To calculate the directional derivative of the solution $z(t, \bar{\theta})$ of an ODE with respect to the parameter $\theta_j$, we can use the derivative structure defined below, evaluating the function in the complex plane:

$$g'(\theta) = \frac{\operatorname{Imag} g(\theta_j + \Delta i)}{\Delta} + O(\Delta^2) \tag{6}$$

where we have written $g(\theta_j + \Delta i)$ instead of $g(\theta_1, \ldots, \theta_{j-1}, \theta_j + \Delta i, \theta_{j+1}, \ldots, \theta_p)$ to simplify notation.

*Step 2: Solve the ODE with a Complex Parameter.* Replace the parameter $\theta_j$ with a complex parameter $\theta_j + \Delta i$, where $0 < \Delta \ll 1$. This gives us a new ODE: $\frac{dz(t, \bar{\theta})}{dt} = g[z(t), \theta_j + \Delta i]$. Solve this ODE to obtain $z(t, \theta_j + \Delta i)$.

*Step 3: Extract the imaginary part.* Once we have the solution, take the imaginary part of this solution: $\operatorname{Imag}[\, z(t, \theta_j + \Delta i)]$.

*Step 4: Divide by* $\Delta$. To approximate the directional derivative, divide the imaginary part by $\Delta$: $\frac{\partial z(t, \bar{\theta})}{\partial \theta_j} \approx \frac{\operatorname{Imag} z(t, \theta_j + \Delta i)}{\Delta}$

By doing so, we compute the differential sensitivity of a model defined as the gradient with respect to the parameters $\theta$ in equation [4] in the main text at their estimated values.

### 2 Models: equations and diagrams

A detailed scheme of each of the models analyzed in this work is shown in Fig. S1:

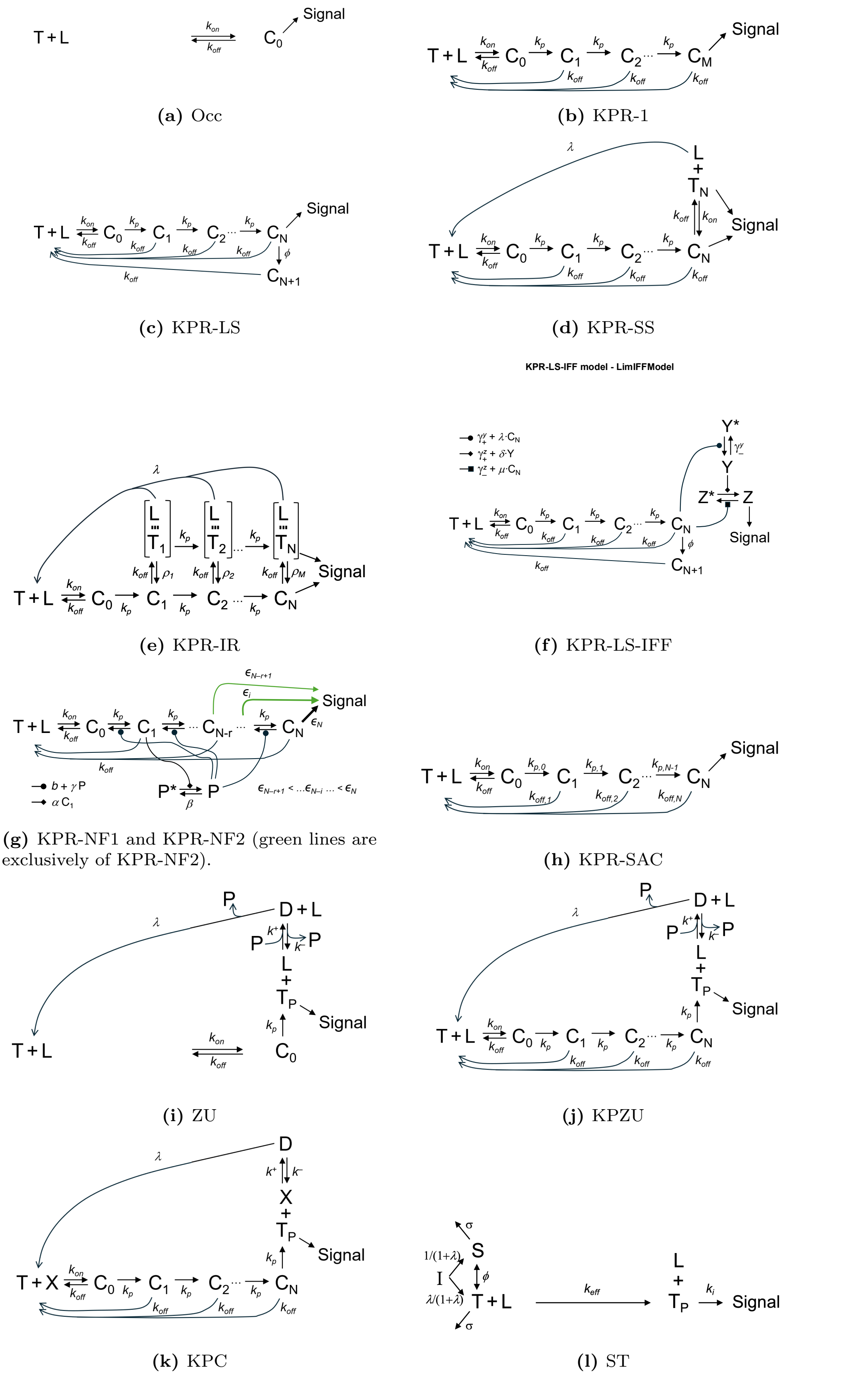


**Figure S1:** Diagrams describing the different models analysed in this paper. (a) Occupancy model; (b) Kinetic proofreading (KPR); (c) KPR with limited signaling; (d) KPR with sustained signaling; (e) KPR with induced rebinding (ligand-receptor encounter-oriented states are indicated by enclosing brackets); (f) KPR with limited signaling and incoherent feed-forward loop; (g) KPR with negative feedback, models 1 and 2; (h) KPR with stabilization activation chain; (i) Zero-order ultrasensibility; (j) KPR with zero-order ultrasensibility; (k) KPR with concentration compensation; (l) Serial triggering.

Models KPR-NF1 and KPR-NF2 (Fig. S1(g)) share the same set of reactions, but differ in the definition of $R(t)$ (the response level). To better clarify this difference, the schemes corresponding to each of these two models are depicted, again, together with their equations (Figs. S2 and S3).

In the following, the ODE system describing each model is provided as well as the specific definition of the response level, $R(t)$, and the corresponding expressions for $E_{max}$ and $EC_{50}$ obtained in [3]. These formulae are used as outputs in the calculations of identifiability, observability and sensitivity.

#### 2.1 Occupancy model (Occ)

The equations of the model are:

$$\begin{cases} \dfrac{dL}{dt} = -k_{on}LT + k_{off}C_0 \\ \dfrac{dT}{dt} = -k_{on}LT + k_{off}C_0 \\ \dfrac{dC_0}{dt} = k_{on}LT - k_{off}C_0 \end{cases} \tag{7}$$

The response in this model is defined as: $R(t) = C_0(t)$.
The maximal response, $E_{max}$, and the half-maximal response ligand concentration, $EC_{50}$, are:

$$E_{max} = T_T,$$

$$EC_{50} = \frac{k_{off}}{k_{on}} + T_T/2.$$

#### 2.2 KPR McKeithan (KPR-1)

The equations of the model are:

$$\begin{cases} \dfrac{dL}{dt} = -k_{on}LT + \displaystyle\sum_{i=0}^{N} k_{off}C_i \\ \dfrac{dT}{dt} = -k_{on}LT + \displaystyle\sum_{i=0}^{N} k_{off}C_i \\ \dfrac{dC_0}{dt} = k_{on}LT - (k_{off} + k_p)C_0 \\ \dfrac{dC_i}{dt} = k_pC_{i-1} - (k_{off} + k_p)C_i; \quad 1 \le i \le N-1 \\ \dfrac{dC_N}{dt} = k_pC_{N-1} - k_{off}C_N \end{cases} \tag{8}$$

The response in this model is defined as: $R(t) = C_N(t)$. The corresponding $E_{max}$ and $EC_{50}$ are:

$$E_{\max} = \left(\frac{k_p}{k_p + k_{off}}\right)^N T_T,$$

$$EC_{50} = \frac{T_T}{2} + \frac{k_{off}}{k_{on}}.$$

##### 2.3 KPR with limited signaling (KPR-LS)

The equations of the model are:

$$\begin{cases} \dfrac{dL}{dt} = -k_{on} LT + \displaystyle\sum_{i=0}^{N+1} k_{off} C_i \\ \dfrac{dT}{dt} = -k_{on} LT + \displaystyle\sum_{i=0}^{N+1} k_{off} C_i \\ \dfrac{dC_0}{dt} = k_{on} LT - (k_{off} + k_p) C_0 \\ \dfrac{dC_i}{dt} = k_p C_{i-1} - (k_{off} + k_p) C_i \qquad \text{for } 1 \le i \le N-1 \\ \dfrac{dC_N}{dt} = k_p C_{N-1} - (k_{off} + \phi) C_N \\ \dfrac{dC_{N+1}}{dt} = \phi C_N - k_{off} C_{N+1} \end{cases} \tag{9}$$

The response in this model is defined as: $R(t) = C_N(t)$, and the corresponding $E_{max}$ and $EC_{50}$ are:

$$E_{\max} = \left( \frac{k_{off}}{k_{off} + \phi} \right) \left( \frac{k_p}{k_p + k_{off}} \right)^N T_T,$$

$$EC_{50} = \frac{T_T}{2} + \frac{k_{off}}{k_{on}}.$$

##### 2.4 KPR with sustained signaling (KPR-SS)

The equations of the model are:

$$\begin{cases} \dfrac{dL}{dt} = -k_{on} LT + \displaystyle\sum_{i=0}^{N} k_{off} C_i - k_{on} LT_N \\ \dfrac{dT}{dt} = -k_{on} LT + \displaystyle\sum_{i=0}^{N-1} k_{off} C_i + \lambda T_N \\ \dfrac{dC_0}{dt} = k_{on} LT - (k_{off} + k_p) C_0 \\ \dfrac{dC_i}{dt} = k_p C_{i-1} - (k_{off} + k_p) C_i \qquad \text{for } 1 \le i \le N-1 \\ \dfrac{dC_N}{dt} = k_p C_{N-1} - k_{off} C_N + k_{on} LT_N \\ \dfrac{dT_N}{dt} = k_{off} C_N - k_{on} LT_N - \lambda T_N \end{cases} \tag{10}$$

The response in this model is defined as: $R(t) = C_N(t) + T_N(t)$, and the corresponding $E_{max}$ and $EC_{50}$ are:

$$E_{\max} = T_T,$$

$$EC_{50} = \frac{\lambda\,(k_{on}\,T_T + 2\,\lambda) + (k_{on}\,T_T - 2\,\lambda)\big(a_1 + \psi^N (2\,\lambda + k_{off})\big)}{4\,k_{on}\,\lambda},$$

with

$$a_1 = \sqrt{(1 - 2\,\psi^N)^2\,\lambda^2 + k_{off}\,\psi^N \big(k_{off}\,\psi^N + 2\,\lambda\,(1 + 2\,\psi^N)\big)}, \text{ and } \psi = \frac{k_p}{k_p + k_{off}}$$

##### 2.5 KPR with induced rebinding (KPR-IR)

The equations of the model are:

$$\begin{cases} \dfrac{dL}{dt} = -k_{on}LT + k_{off}C_0 + \sum_{i=1}^{N} \lambda_r \overline{C}_i \\ \dfrac{dT}{dt} = -k_{on}LT + k_{off}C_0 + \sum_{i=1}^{N} \lambda_r \overline{C}_i \\ \dfrac{dC_0}{dt} = k_{on}LT - (k_{off} + k_p)C_0 \\ \dfrac{dC_i}{dt} = k_pC_{i-1} - (k_{off} + k_p)C_i + \rho_i C_i^*;\ 1 \le i \le N-1 \\ \dfrac{dC_N}{dt} = k_pC_{N-1} - k_{off}C_N + \rho_N C_N^* \\ \dfrac{d\overline{C}_1}{dt} = k_{off}C_1 - (\rho_1 + \lambda_r + k_p)\overline{C}_1 \\ \dfrac{d\overline{C}_i}{dt} = k_p\overline{C}_{i-1} + k_{off}C_i - (\rho_i + \lambda_r + k_p)\overline{C}_i;\ 2 \le i \le N-1 \\ \dfrac{d\overline{C}_N}{dt} = k_p\overline{C}_{N-1} + k_{off}C_N - (\rho_N + \lambda_r)\overline{C}_N \end{cases} \tag{11}$$

where $\alpha_i = \dfrac{k_{off}}{k_{off} + \rho_i}$ and $\rho_i = \rho$ for $i \le 21$. The response in this model is defined as: $R(t) = \overline{C}_N(t) + C_N(t)$.

In order to derive a workable approximate version of this ODE system, we assume that rebinding after unbinding occurs fast enough so that bound states and unbound but encounter-oriented states can be merged into one effective state [4]. In mathematical terms, this means that, in a coarse-grained observation timescale, we can safely assume that $k_{off}C_i = \rho_i\overline{C}_i$ (in statistical thermodynamics this hypothesis is often referred to as the local steady state hypothesis). Then, defining $C_i^* = C_i + \overline{C}_i$, one has $\overline{C}_i = \dfrac{k_{off}}{k_{off} + \rho_i}C_i^*$ and $C_i = \dfrac{\rho_i}{k_{off} + \rho_i}C_i^*$. Consequently, the above ODE system reduces to the following one [3]:

$$\begin{cases} \dfrac{dL}{dt} = -k_{on}LT + k_{off}C_0 + \lambda \sum_{i=1}^{N} \alpha_i C_i^* \\ \dfrac{dT}{dt} = -k_{on}LT + k_{off}C_0 + \lambda \sum_{i=1}^{N} \alpha_i C_i^* \\ \dfrac{dC_0}{dt} = k_{on}LT - (k_{off} + k_p)C_0 \\ \dfrac{dC_i^*}{dt} = k_p(C_{i-1}^* - C_i^*) - \alpha_i C_i^*;\ 1 \le i \le N-1 \\ \dfrac{dC_N^*}{dt} = k_pC_{N-1}^* - \alpha_i C_N^* \end{cases} \tag{12}$$

Note that in this approximate system the response level is $R(t) = C_N^*(t)$ and, hence, it still verifies $R(t) = \overline{C}_N(t) + C_N(t)$. The corresponding $E_{max}$ and $EC_{50}$ are:

$$E_{\max} = \frac{k_p}{\alpha_N}\left(\prod_{i=1}^{N-1} \frac{k_p}{\alpha_i + k_p}\right)\mu T_T,$$

$$EC_{50} = \frac{k_{off} + k_p}{k_{on}}\mu + \frac{T_T}{2},$$

with

$$\mu = 1 + \left(1 + \frac{k_p}{\alpha_N}\right)\prod_{i=1}^{N-1} \frac{k_p}{\alpha_i + k_p}, \text{ and } \alpha_i = \frac{k_{off}}{k_{off} + \rho_i}$$

The identifiability analyses were performed for $N = 2$, *i.e.*, two kinetic proofreading-steps, to limit the complexity of calculations. In this case:

$$E_{max} = \frac{k_p^2\left(k_{\mathit{off}}+\rho_1\right)\left(k_{\mathit{off}}+\rho_2\right)}{k_{\mathit{off}}\,k_p\left(k_{\mathit{off}}+\rho_1\right)+k_{\mathit{off}}^2}\left[1+\frac{k_p(k_{\mathit{off}}+\rho_1)}{k_{\mathit{off}}+k_p(k_{\mathit{off}}+\rho_1)}\left(1+\frac{k_p\left(k_{\mathit{off}}+\rho_2\right)}{k_{\mathit{off}}}\right)\right]T_T,$$

$$EC_{50} = \frac{k_{\mathit{off}}+k_p}{k_{on}}\left[1+\frac{k_p(k_{\mathit{off}}+\rho_1)}{k_{\mathit{off}}+k_p(k_{\mathit{off}}+\rho_1)}\left(1+\frac{k_p\left(k_{\mathit{off}}+\rho_2\right)}{k_{\mathit{off}}}\right)\right]+\frac{T_T}{2}.$$

#### 2.6 KPR with limited signaling and incoherent feed-forward loop (KPR-LS-IFF)

The equations of the model are:

$$\begin{cases} \dfrac{dL}{dt} = -k_{on}LT + \displaystyle\sum_{i=0}^{N+1} k_{\mathit{off}}C_i \\ \dfrac{dT}{dt} = -k_{on}LT + \displaystyle\sum_{i=0}^{N+1} k_{\mathit{off}}C_i \\ \dfrac{dC_0}{dt} = k_{on}LT - (k_{\mathit{off}}+k_p)C_0 \\ \dfrac{dC_i}{dt} = k_pC_{i-1} - (k_{\mathit{off}}+k_p)C_i & \text{for } 1 \le i \le N-1 \\ \dfrac{dC_N}{dt} = k_pC_{N-1} - (k_{\mathit{off}}+\phi)C_N \\ \dfrac{dC_{N+1}}{dt} = \phi C_N - k_{\mathit{off}}C_{N+1} \\ \dfrac{dY}{dt} = \gamma_+^y(Y_T-Y) - \gamma_-^yY + \sigma C_N(Y_T-Y) \\ \dfrac{dZ}{dt} = \gamma_+^z(Z_T-Z) - \gamma_-^zZ + \delta Y(Z_T-Z) - \mu C_N Z \end{cases} \tag{13}$$

The response in this model is defined as $R(t) = Z(t)$.

Due to its length, the corresponding $E_{max}$ and $EC_{50}$ for $N = 5$ can be found in in our online repository (`https://github.com/Xabo-RB/Inmunology-analysis.git`).

#### 2.7 KPR with negative feedback I (KPR-NF1)

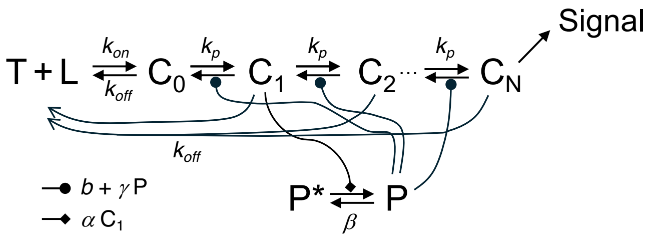


**Figure S2:** Conceptual scheme of the KPR with Negative Feedback model I.

The equations of the model are:

$$\begin{cases} \frac{dL}{dt} = -k_{on}LT + k_{off}\sum_{i=0}^{N} C_i \\ \frac{dT}{dt} = -k_{on}LT + k_{off}\sum_{i=0}^{N} C_i \\ \frac{dC_0}{dt} = k_{on}LT + (b+\gamma P)C_1 - (k_{off}+k_p)C_0 \\ \frac{dC_i}{dt} = k_pC_{i-1} - (k_{off}+k_p+b+\gamma P)C_i + (b+\gamma P)C_{i+1} \quad \text{for } 1 \le i \le N-1 \\ \frac{dC_N}{dt} = k_pC_{N-1} - (k_{off}+b+\gamma P)C_N \\ \frac{dP}{dt} = \alpha C_1(P_T - P) - \beta P \end{cases} \tag{14}$$

The response in this model is defined as: $R(t) = C_N(t)$.

Since the symbolic expressions for $E_{max}$ and $EC_{50}$ are extensive and cannot be explicitly shown, we express the response in terms of the phosphatase. These resulting equations are valid only under the assumption of steady state; this remark also applies to the identifiability results. The corresponding $E_{max}$ and $EC_{50}$, based on the derivation from [5], are:

$$E_{\max} = \left( \frac{1 - \frac{q_-}{q_+}}{1 - (\frac{q_-}{q_+})^{N+1}} \right) q_-^N T_T$$

where $\quad q_{\pm} = \dfrac{k_p + b + \gamma P + k_{off} \;\pm\; \sqrt{(k_p + b + \gamma P + k_{off})^2 \;-\; 4k_p\,(b+\gamma P)}}{2\,(b+\gamma P)},$

$$EC_{50} = \frac{T_T}{2} + \frac{k_{off}}{k_{on}}.$$

#### 2.8 KPR with negative feedback II (KPR-NF2)

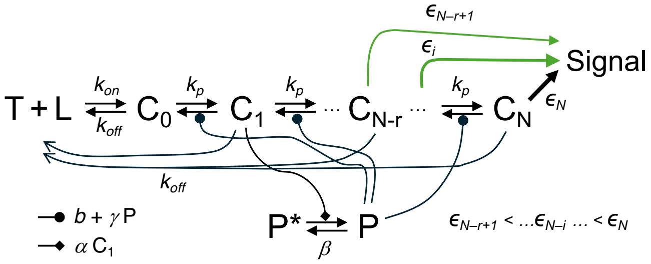


**Figure S3:** Conceptual scheme of the KPR with Negative Feedback model II.

The equations of this model are the same as for KPR-NF1. The response, however, is:

$R(t) = \sum_{i=N-r+1}^{N} \epsilon_i C_i$, where $\epsilon_i < \epsilon_{i+1}$ (for instance, $\epsilon_i = e^{-k\,(N-i)}$ for $i = N-r+1,\ldots,N$).

The corresponding $E_{max}$ and $EC_{50}$ (for $N = 2$ and $r = 3$) are:

$$E_{\max} = T_T \left( \frac{\epsilon_0 \left(b\left(\left(1-3q_-^2\right)q_+^2 + q_-^2 + q_-q_+\right) + k_{off}\left(-2q_-^2q_+^2 + q_-^2 + q_-q_+ + q_+^2\right)\right)}{\left(q_-^2 + q_-q_+ + q_+^2\right)\left(b + k_{off} + 2k_p + \gamma P\right)} \right.$$

$$\left. + \frac{(k_p+\gamma P)\left(-3q_-^2q_+^2 + q_-^2 + q_-q_+ + q_+^2\right) + \epsilon_1\left(k_p\left(\left(1-3q_-^2\right)q_+^2 + q_-^2 + q_-q_+\right) - k_{off}q_-^2q_+^2\right)}{\left(q_-^2 + q_-q_+ + q_+^2\right)\left(b + k_{off} + 2k_p + \gamma P\right)} + \frac{q_-^2q_+^2}{q_-^2 + q_-q_+ + q_+^2} \right)$$

where $q_{\pm} = \dfrac{k_p + b + \gamma P + k_{off} \;\pm\; \sqrt{(k_p + b + \gamma P + k_{off})^2 \;-\; 4k_p\,(b+\gamma P)}}{2\,(b+\gamma P)},$

$$EC_{50} = \frac{T_T}{2} + \frac{k_{off}}{k_{on}}.$$

The derivation of these formulae can be found in our online repository (`https://github.com/Xabo-RB/Inmunology-analysis.git`)

#### 2.9 KPR with stabilizing activation chain (KPR-SAC)

The equations of the model are:

$$\begin{cases} \frac{dL}{dt} = -k_{on}LT + \sum_{i=0}^{N} k_{off}(i)C_i \\ \frac{dT}{dt} = -k_{on}LT + \sum_{i=0}^{N} k_{off}(i)C_i \\ \frac{dC_0}{dt} = k_{on}LT - (k_{off}(0) + k_p(0))C_0 \\ \frac{dC_i}{dt} = k_p(i-1)C_{i-1} - (k_{off}(i) + k_p(i))C_i \quad \text{for } 1 \leq i \leq N-1 \\ \frac{dC_N}{dt} = k_p(N-1)C_{N-1} - k_{off}(N)C_N \\ k_{off}(i) = k_{off} \cdot \frac{1+i}{1+r \cdot i} \\ k_p(i) = k_p \cdot r_p^i \end{cases} \tag{15}$$

The response in this model is defined as: $R(t) = C_N(t)$. The corresponding $E_{max}$ and $EC_{50}$ are:

$$E_{\max} = T_T \left[ 1 + F_N \left( 1 + \prod_{i=1}^{N-1} F_i + \sum_{i=1}^{N-2} \left( \prod_{j=1}^{i} F_{N-j} \right) \right) \right]^{-1}$$

with

$$F_i = \left[ \frac{k_{off} \frac{1+i}{i \cdot r + 1} + k_p \cdot r_p^i}{k_p \cdot r_p^{i-1}} \right], \quad \text{for} \quad i = 1 \text{ to } N-1; \quad F_N = \left[ \frac{k_{off} \cdot \frac{1+N}{N \cdot r + 1}}{k_p \cdot r_p^{N-1}} \right].$$

and

$$EC_{50} = \frac{T_T}{2} + \frac{k_{off} + k_p}{G\, k_{on}}.$$

with

$$G = \frac{1}{F_N} \cdot \prod_{i=1}^{N-1} \frac{1}{F_i} \left[ 1 + F_N \left( 1 + \prod_{i=1}^{N-1} F_i + \sum_{i=1}^{N-2} \left( \prod_{j=1}^{i} F_{N-j} \right) \right) \right].$$

Thus, for $N = 2$:

$$E_{\max} = T_T \left[ 1 + \frac{3 \cdot k_{off}}{k_p r_p (2 \cdot r + 1)} \left( 1 + \frac{2 \cdot k_{off}}{k_p (r+1)} + r_p \right) \right].$$

$$EC50 = \frac{T_T}{2} + \frac{k_{off} + k_p}{k_{on}} \cdot \left[ \frac{3 \cdot k_{off}}{k_p r_p (2 \cdot r + 1)} \cdot \left( \frac{2 \cdot k_{off}}{r+1} + r_p \right) \cdot \left( 1 + \frac{3 \cdot k_{off}}{k_p r_p (2 \cdot r + 1)} \left( 1 + \frac{2 \cdot k_{off}}{k_p (r+1)} + r_p \right) \right) \right].$$

##### 2.10 Zero-order ultrasensitivity (ZU)

The equations of the model are:

$$\begin{cases} \frac{dT}{dt} = -k_{on}TL + \lambda D + k_{off}C_0 \\ \frac{dL}{dt} = -k_{on}TL + (k_p + k_{off})C_0 \\ \frac{dC_0}{dt} = k_{on}TL - (k_{off} + k_p)C_0 \\ \frac{dD}{dt} = k_+T_pP - (k_- + \lambda)D \\ \frac{dT_p}{dt} = -k_+T_pP + k_-D + k_pC_0 \\ \frac{dP}{dt} = -k_+T_pP + k_-D + \lambda D \end{cases} \tag{16}$$

The response in this model is defined as: $R(t) = T_p(t)$, and the corresponding $E_{max}$ and $EC_{50}$ are:

$$E_{\max} = \frac{\alpha + \sqrt{\alpha^2 + 4\,k_p^2\,\lambda^2\,k_+\,(k_- + \lambda)\,T_T}}{2\,k_p\,k_+\,\lambda}$$

with

$$\alpha = \big(k_p\,k_+\,\lambda\,T_T - k_+\,\lambda\,(\lambda + k_p)\,P_T - k_p\,\lambda\,(k_- + \lambda)\big).$$

##### 2.11 KPR with zero-order ultraspecificity (KPZU)

The equations of the model are:

$$\begin{cases} \frac{dT}{dt} = -k_{on}TL + \lambda D + k_{off}\sum_{i=0}^{n} C_i \\ \frac{dL}{dt} = -k_{on}TL + k_pC_N + k_{off}\sum_{i=0}^{n} C_i \\ \frac{dT_p}{dt} = k_pC_N - k_+T_pP + k_-D \\ \frac{dC_0}{dt} = -k_{off}C_0 - k_pC_0 + k_{on}TL \\ \frac{dC_i}{dt} = -k_{off}C_i - k_pC_i + k_pC_{i-1} \quad \text{for } 1 \le i \le N \\ \frac{dP}{dt} = -k_+T_pP + k_-D + \lambda D \\ \frac{dD}{dt} = k_+T_pP - k_-D - \lambda D \end{cases} \tag{17}$$

The response in this model is defined as: $R(t) = T_p(t)$. The corresponding $E_{max}$ and $EC_{50}$ are:

$$E_{\max} = \frac{\alpha + \sqrt{\alpha^2 + 4\,\Phi^2\,\lambda^2\,k_+\,(k_- + \lambda)\,T_T}}{2\,\Phi\,k_+\,\lambda}$$

with

$$\alpha = \big(\Phi\,k_+\,\lambda\,T_T - k_+\,\lambda\,(\lambda + \Phi)\,P_T - \Phi\,\lambda\,(k_- + \lambda)\big),\ \Phi = k_p\,\psi^N\left(\frac{1-\psi}{1-\psi^{N+1}}\right),\ \text{and}\ \psi = \frac{k_p}{k_{off} + k_p}.$$

$P_T$ is defined by the conservation equation: $P_T = P + D$.

##### 2.12 KPR with concentration compensation (KPC)

The equations of the model are:

$$\begin{cases} \dfrac{dT}{dt} = -k_{on}TX + \lambda D + k_{\mathit{off}}\displaystyle\sum_{i=0}^{n} C_i \\ \dfrac{dX}{dt} = -(k_{on}TX + k_{+}T_pX) + k_pC_N + k_{\mathit{off}}\displaystyle\sum_{i=0}^{n} C_i + k_{-}D + \lambda D \\ \dfrac{dC_i}{dt} = -(k_{\mathit{off}}C_i + k_pC_i) + k_pC_{i-1} \quad \text{for } 1 \leq i \leq N \\ \dfrac{dC_0}{dt} = -(k_{\mathit{off}} + k_p)C_0 + k_{on}TX \\ \dfrac{dT_p}{dt} = k_pC_N - k_{+}T_pX + k_{-}D \\ \dfrac{dD}{dt} = k_{+}T_pX - (k_{-} + \lambda)D \end{cases} \tag{18}$$

The response in this model is defined as: $R(t) = T_p(t)$. The corresponding $E_{max}$ and $EC_{50}$ are:

$$E_{\max} = \frac{k_{on}\,(k_{-} + \lambda)\,k_p^{N+1}\,T_T}{k_{+}\,\lambda\,(k_{\mathit{off}} + k_p)^{N+1} + k_{on}\,(k_{-} + \lambda)\,k_p^{N+1}},$$

$$EC_{50} = \frac{T_T}{2} + \frac{k_{\mathit{off}}\left(k_{-}\,k_{on}\,k_p^{N+1} + \left(k_{on}\,k_p^{N+1} + k_{+}\,\lambda\,(k_{\mathit{off}} + k_p)^{N+1}\right)\right)}{k_{+}\,k_{on}\,\left(k_{\mathit{off}}\left(k_p^{N+1} + \lambda\,(k_{\mathit{off}} + k_p)^{N}\right) - \lambda\,k_p\,\left(k_p^{N} - (k_{\mathit{off}} + k_p)^{N}\right)\right)}.$$

##### 2.13 Serial triggering (ST)

The equations of the model are:

$$\begin{cases} \dfrac{dS}{dt} = -\phi\left(S - \dfrac{1}{\lambda}T\right) + \sigma\left(\dfrac{1}{1+\lambda} - S\right) \\ \dfrac{dT}{dt} = \phi\left(S - \dfrac{1}{\lambda}T\right) + \sigma\left(\dfrac{\lambda}{1+\lambda} - T\right) - k_{\text{eff}}T^hL^h \\ \dfrac{dT_p}{dt} = k_{\text{eff}}T^hL^h - k_iT_p \end{cases} \tag{19}$$

Here $S$, $T$ and $T_p$ are fractional quantities with respect total TCRs in the resting state (absence of ligand).
The response in this model is defined as: $R(t) = T_p(t)$. The corresponding $E_{max}$ and $EC_{50}$ for $h = 2$ are:

$$E_{\max} = \frac{\sigma\left(\sigma\,\lambda + \phi\,(1+\lambda)\right)}{k_i\,(1+\lambda)\,(\sigma+\phi)},$$

$$EC_{50} = \frac{\sqrt{2\,\sigma\,(1+\lambda)\,\left(\phi + \lambda\,(\sigma+\phi)\right)}}{\lambda\,\sqrt{k_{\mathit{eff}}\,(\sigma+\phi)}}.$$

### 3 Supplementary results

##### 3.1 Identifiability results

**Table S1:** Identifiability results with measurements of lymphocyte concentration and T-cell response $R(t)$. Variables with an asterisk (*) denote initial conditions of state variables (i.e., observability). "Id": identifiable parameters and variables. "Non-id": non identifiable parameters and variables. In the "Id" columns, variables in boldface indicate structural global identifiability (SGI). The index $i$ goes from 1 to $N$, except in the cases labeled with **†** in the **Known output** column, where it goes from 1 to $N-1$.

| | | **Additional known output (parameters)** | | | | | | | |
|---|---|---|---|---|---|---|---|---|---|
| | | — | | $k_{on}$ | | $k_p$ | | $k_{on}$, $k_p$ | |
| **Models** | **Known output** | Id | Non-id | Id | Non-id | Id | Non-id | Id | Non-id |
| **Occ** | $T(t)$ | $\boldsymbol{k_{on}}$ | $k_{off}$, $L^*$, $C_0^*$ | *None* | $k_{off}$, $L^*$, $C_0^*$ | — | — | — | — |
| | $T(t)$, $R(t)$ | $\boldsymbol{k_{on}, k_{off}, L^*}$ | *None* | $\boldsymbol{k_{off}, L^*}$ | *None* | — | — | — | — |
| | $R(t)$ | $\boldsymbol{k_{on}}$ | $k_{off}$, $L^*$, $T^*$ | *None* | $k_{off}$, $L^*$, $T^*$ | — | — | — | — |
| | $R(t)$, $T_T$ | $\boldsymbol{k_{on}, k_{off}, L^*}$ | *None* | $\boldsymbol{k_{off}, L^*}$ | *None* | — | — | — | — |
| | $T_T$ | *None* | $k_{on}$, $k_{off}$, $L^*$, $C_0^*$ | *None* | $k_{off}$, $C_0^*$, $L^*$ | — | — | — | — |
| **KPR-1** | $T(t)$ | $\boldsymbol{k_{on}}$ | $k_{off}$, $k_p$, $L^*$, $C_0^*$, $C_i^*$ | *None* | $k_{off}$, $k_p$, $L^*$, $C_0^*$, $C_i^*$ | $\boldsymbol{k_{on}}$ | $k_{off}$, $L^*$, $C_0^*$, $C_i^*$ | *None* | $k_{off}$, $L^*$, $C_0^*$, $C_i^*$ |
| | $T(t)$, $R(t)$**†** | $\boldsymbol{k_{on}, k_{off}, k_p, L^*}$, $\mathbf{C}_0^*$, $\mathbf{C}_\mathbf{i}^*$ | *None* | $\boldsymbol{k_{off}, k_p, L^*}$, $\mathbf{C}_0^*$, $\mathbf{C}_\mathbf{i}^*$ | *None* | $\boldsymbol{k_{on}, k_{off}, L^*}$, $\mathbf{C}_0^*$, $\mathbf{C}_\mathbf{i}^*$ | *None* | $\boldsymbol{k_{off}, L^*, C_0^*, C_i^*}$ | *None* |
| | $R(t)$**†** | *None* | $k_{on}$, $k_{off}$, $k_p$, $L^*$, $T^*$, $C_0^*$, $C_i^*$ | $k_{off}$, $k_p$, $L^*$, $T^*$, $C_0^*$, $C_i^*$ | *None* | $\boldsymbol{k_{on}, k_{off}}$, $L^*$, $T^*$, $\boldsymbol{C_0^*, C_i^*}$ | *None* | $\boldsymbol{k_{off}}$, $L^*$, $T^*$, $\boldsymbol{C_0^*, C_i^*}$ | *None* |
| | $R(t)$, $T_T$**†** | $k_{on}$, $k_{off}$, $k_p$, $L^*$, $C_0^*$, $C_i^*$ | *None* | $\boldsymbol{k_{off}, k_p, L^*}$, $\mathbf{C}_0^*$, $\mathbf{C}_\mathbf{i}^*$ | *None* | $\boldsymbol{k_{off}, k_{on}, L^*}$, $\mathbf{C}_0^*$, $\mathbf{C}_\mathbf{i}^*$ | *None* | $\boldsymbol{k_{off}, L^*, C_0^*, C_i^*}$ | *None* |
| | $T_T$ | *None* | $k_{on}$, $k_{off}$, $k_p$, $L^*$, $C_0^*$, $C_i^*$ | *None* | $k_{off}$, $k_p$, $L^*$, $C_0^*$, $C_i^*$ | *None* | $k_{on}$, $k_{off}$, $L^*$, $C_0^*$, $C_i^*$ | *None* | $k_{off}$, $L^*$, $C_0^*$, $C_i^*$ |
| **KPR-LS** | $T(t)$ | $\boldsymbol{k_{on}}$ | $k_{off}$, $k_p$, $\phi$, $L^*$, $C_0^*$, $C_i^*$, $C_{N+1}^*$ | *None* | $k_{off}$, $k_p$, $\phi$, $L^*$, $C_0^*$, $C_i^*$, $C_{N+1}^*$ | $\boldsymbol{k_{on}}$ | $k_{off}$, $\phi$, $L^*$, $C_0^*$, $C_i^*$, $C_{N+1}^*$ | *None* | $k_{off}$, $\phi$, $L^*$, $C_0^*$, $C_i^*$, $C_{N+1}^*$ |
| | $T(t)$, $R(t)$**†** | $\boldsymbol{k_{on}, k_{off}, k_p, \phi, L^*, C_0^*, C_i^*, C_{N+1}^*}$ | *None* | $\boldsymbol{k_{off}, k_p, \phi, L^*, C_0^*, C_i^*, C_{N+1}^*}$ | *None* | $\boldsymbol{k_{on}, k_{off}, \phi, L^*, C_0^*, C_i^*, C_{N+1}^*}$ | *None* | $\boldsymbol{k_{off}, \phi, L^*, C_0^*, C_i^*, C_{N+1}^*}$ | *None* |
| | $R(t)$**†** | $\boldsymbol{k_{on}, k_{off}, k_p, \phi}$, $L^*$, $T^*$, $\boldsymbol{C_0^*, C_i^*, C_{N+1}^*}$ | *None* | $\boldsymbol{k_{off}, k_p, \phi}$, $L^*$, $T^*$, $\boldsymbol{C_0^*, C_i^*, C_{N+1}^*}$ | *None* | $\boldsymbol{k_{on}, k_{off}, \phi}$, $L^*$, $T^*$, $\boldsymbol{C_0^*, C_i^*, C_{N+1}^*}$ | *None* | $\boldsymbol{k_{off}, \phi}$, $L^*$, $T^*$, $\boldsymbol{C_0^*, C_i^*, C_{N+1}^*}$ | *None* |
| | $R(t)$, $T_T$**†** | $\boldsymbol{k_{on}, k_{off}, k_p, \phi, L^*, C_0^*, C_i^*, C_{N+1}^*}$ | *None* | $\boldsymbol{k_{off}, k_p, \phi, L^*, C_0^*, C_i^*, C_{N+1}^*}$ | *None* | $\boldsymbol{k_{on}, k_{off}, \phi, L^*, C_0^*, C_i^*, C_{N+1}^*}$ | *None* | $\boldsymbol{k_{off}, \phi, L^*, C_0^*, C_i^*, C_{N+1}^*}$ | *None* |
| | $T_T$ | *None* | $k_{on}$, $k_{off}$, $k_p$, $\phi$, $L^*$, $C_0^*$, $C_i^*$, $C_{N+1}^*$ | *None* | $k_{off}$, $k_p$, $\phi$, $L^*$, $C_0^*$, $C_i^*$, $C_{N+1}^*$ | *None* | $k_{on}$, $k_{off}$, $\phi$, $L^*$, $C_0^*$, $C_i^*$, $C_{N+1}^*$ | *None* | $k_{off}$, $\phi$, $L^*$, $C_0^*$, $C_i^*$, $C_{N+1}^*$ |

**Table S1 (cont.):** Identifiability results with measurements of lymphocyte concentration and T-cell response $R(t)$. Variables with an asterisk (*) denote initial conditions of state variables (i.e., observability). "Id": identifiable parameters and variables. "Non-id": non identifiable parameters and variables. In the "Id" columns, variables in boldface indicate structural global identifiability (SGI). The index $i$ goes from 1 to $N$, except in the cases labeled with † in the **Known output** column, where it goes from 1 to $N-1$.

| | | **Additional known output (parameters)** | | | | | | | |
|---|---|---|---|---|---|---|---|---|---|
| | | — | | $k_{on}$ | | $k_p$ | | $k_{on}$, $k_p$ | |
| **Models** | **Known output** | Id | Non-id | Id | Non-id | Id | Non-id | Id | Non-id |
| **KPR-SS** | $T(t)$ | $k_{on}, k_{off}, k_p, \lambda, T_N^*, L^*, C_0^*, C_i^*$ | *None* | $k_{off}, k_p, \lambda, T_N^*, L^*, C_0^*, C_i^*$ | *None* | $\boldsymbol{k_{on}, k_{off}, \lambda, T_N^*, L^*, C_0^*, C_i^*}$ | *None* | $\boldsymbol{k_{off}, \lambda, T_N^*, L^*, C_0^*, C_i^*}$ | *None* |
| | $T(t)$, $R(t)$ | $\boldsymbol{k_{on}, k_{off}, k_p, \lambda, T_N^*, L^*, C_0^*, C_i^*}$ | *None* | $\boldsymbol{k_{off}, k_p, \lambda, T_N^*, L^*, C_0^*, C_i^*}$ | *None* | $\boldsymbol{k_{on}, k_{off}, \lambda, T_N^*, L^*, C_0^*, C_i^*}$ | *None* | $\boldsymbol{k_{off}, \lambda, T_N^*, L^*, C_0^*, C_i^*}$ | *None* |
| | $R(t)$ | $k_{on}, k_{off}, k_p, \lambda, T^*, L^*, T_N^*, C_0^*, C_i^*$ | *None* | $k_{off}, k_p, \lambda, T^*, L^*, T_N^*, C_0^*, C_i^*$ | *None* | $k_{on}, k_{off}, \lambda, T^*, L^*, T_N^*, C_0^*, C_i^*$ | *None* | $\boldsymbol{k_{off}, \lambda, T^*, T_N^*, L^*, C_0^*, C_i^*}$ | *None* |
| | $R(t)$, $T_T$ | $\boldsymbol{k_{on}, k_{off}, k_p, \lambda, L^*, T_N^*, C_0^*, C_i^*}$ | *None* | $k_{off}, k_p, \lambda, L^*, T_N^*, C_0^*, C_i^*$ | *None* | $\boldsymbol{k_{on}, k_{off}, \lambda, L^*, T_N^*, C_0^*, C_i^*}$ | *None* | $k_{off}, \lambda, L^*, T_N^*, C_0^*, C_i^*$ | *None* |
| | $T_T$ | *None* | $k_{on}, k_{off}, k_p, \lambda, L^*, T_N^*, C_0^*, C_i^*,$ | *None* | $k_{off}, k_p, \lambda, L^*, T_N^*, C_0^*, C_i^*, C_N^*$ | *None* | $k_{on}, k_{off}, \lambda, L^*, T_N^*, C_0^*, C_i^*, C_N^*$ | *None* | $k_{off}, \lambda, L^*, T_N^*, C_0^*, C_i^*, C_N^*$ |
| **KPR-IR** | $T(t)$ | $k_{on}, k_{off}, k_p, \lambda, \rho_1, \rho_i, L^*, \bar{C}_0^*, \bar{C}_i^*$ | *None* | $k_{off}, k_p, \lambda, \rho_1, \rho_i, L^*, \bar{C}_0^*, \bar{C}_i^*$ | *None* | $\boldsymbol{k_{on}, k_{off}, \lambda, \rho_1, \rho_i, L^*, \bar{C}_0^*, \bar{C}_i^*}$ | *None* | $\boldsymbol{k_{off}, \lambda, \rho_1, \rho_i, L^*, \bar{C}_0^*, \bar{C}_i^*}$ | *None* |
| | $T(t)$, $R(t)$† | $\boldsymbol{k_{on}, k_{off}, k_p, \lambda, \rho_1, \rho_i, L^*, \bar{C}_0^*, \bar{C}_i^*}$ | *None* | $\boldsymbol{k_{off}, k_p, \lambda, \rho_1, \rho_i, L^*, \bar{C}_0^*, \bar{C}_i^*}$ | *None* | $\boldsymbol{k_{on}, k_{off}, \lambda, \rho_1, \rho_i, L^*, \bar{C}_0^*, \bar{C}_i^*}$ | *None* | $\boldsymbol{k_{off}, \lambda, \rho_1, \rho_i, L^*, \bar{C}_0^*, \bar{C}_i^*}$ | *None* |
| | $R(t)$† | $\boldsymbol{k_{on}, k_{off}, k_p, \lambda, \rho_1}$, $\boldsymbol{\rho_i}$, $T^*, L^*$, $\boldsymbol{\bar{C}_0^*, \bar{C}_i^*}$ | *None* | $\boldsymbol{k_{off}, k_p, \lambda, \rho_1}$, $\boldsymbol{\rho_i}$, $T^*, L^*$, $\boldsymbol{\bar{C}_0^*, \bar{C}_i^*}$ | *None* | $\boldsymbol{k_{on}, k_{off}, \lambda, \rho_1}$, $\boldsymbol{\rho_i}$, $T^*, L^*$, $\boldsymbol{\bar{C}_0^*, \bar{C}_i^*}$ | *None* | $\boldsymbol{k_{off}, \lambda, \rho_1}$, $\boldsymbol{\rho_i}$, $T^*, L^*$, $\boldsymbol{\bar{C}_0^*, \bar{C}_i^*}$ | *None* |
| | $R(t)$, $T_T$† | $\boldsymbol{k_{on}, k_{off}, k_p, \lambda, \rho_1, \rho_i, L^*, \bar{C}_0^*, \bar{C}_i^*}$ | *None* | $\boldsymbol{k_{off}, k_p, \lambda, \rho_1, \rho_i, L^*, \bar{C}_0^*, \bar{C}_i^*}$ | *None* | $\boldsymbol{k_{on}, k_{off}, \lambda, \rho_1, \rho_i, L^*, \bar{C}_0^*, \bar{C}_i^*}$ | *None* | $\boldsymbol{k_{off}, \lambda, \rho_1, \rho_i, L^*, \bar{C}_0^*, \bar{C}_i^*}$ | *None* |
| | $T_T$ | $k_{on}, k_{off}, k_p, \lambda, \rho_1, \rho_i, L^*, \bar{C}_0^*, \bar{C}_i^*$ | *None* | $k_{off}, k_p, \lambda, \rho_1, \rho_i, L^*, \bar{C}_0^*, \bar{C}_i^*$ | *None* | $\boldsymbol{k_{on}, k_{off}, \lambda, \rho_1, \rho_i, L^*, \bar{C}_0^*, \bar{C}_i^*}$ | *None* | $\boldsymbol{k_{off}, \lambda, \rho_1, \rho_i, L^*, \bar{C}_0^*, \bar{C}_i^*}$ | *None* |
| **KPR-LS-IFF** | $T(t)$ | $\boldsymbol{k_{on}}$ | $k_{off}, k_p, \lambda, \gamma^{y/z}_{+/-}, \phi, \mu, \delta, L^*, Z^*, Z_T, C_0^*, C_i^*, C_{N+1}^*, Y^*, Y_T$ | *None* | $k_{off}, k_p, \lambda, \gamma^{y/z}_{+/-}, \phi, \mu, \delta, L^*, Z^*, Z_T, C_0^*, C_i^*, C_{N+1}^*, Y^*, Y_T$ | $\boldsymbol{k_{on}}$ | $k_{off}, \lambda, \gamma^{y/z}_{+/-}, \phi, \mu, \delta, L^*, Z^*, Z_T, C_0^*, C_i^*, C_{N+1}^*, Y^*, Y_T$ | *None* | $k_{off}, \lambda, \gamma^{y/z}_{+/-}, \phi, \mu, \delta, L^*, Z^*, Z_T, C_0^*, C_i^*, C_{N+1}^*, Y^*, Y_T$ |
| | $T(t)$, $R(t)$ | $k_{on}, k_{off}, k_p, \lambda, \gamma^{y/z}_{+/-}, \phi, \mu, L^*, Z_T, C_0^*, C_i^*, C_{N+1}^*$ | $\delta, Y^*, Y_T$ | $k_{off}, k_p, \lambda, \gamma^{y/z}_{+/-}, \phi, \mu, L^*, Z_T, C_0^*, C_i^*, C_{N+1}^*$ | $\delta, Y^*, Y_T$ | $k_{on}, k_{off}, \lambda, \gamma^{y/z}_{+/-}, \phi, \mu, L^*, Z_T, C_0^*, C_i^*, C_{N+1}^*$ | $\delta, Y^*, Y_T$ | $k_{off}, \lambda, \gamma^{y/z}_{+/-}, \phi, \mu, L^*, Z_T, C_0^*, C_i^*, C_{N+1}^*$ | $\delta, Y^*, Y_T$ |
| | $R(t)$ | $k_{off}, k_p, \gamma^{y/z}_{+/-}, \phi, Z_T$ | $k_{on}, \lambda, \delta, \mu, L^*, T^*, Y^*, Y_T, C_0^*, C_i^*, C_{N+1}^*$ | $k_{off}, k_p, \lambda, \gamma^{y/z}_{+/-}, \phi, \mu, L^*, T^*, Z_T, C_0^*, C_i^*, C_{N+1}^*$ | $\delta, Y^*, Y_T$ | $k_{off}, \gamma^{y/z}_{+/-}, \phi, Z_T$ | $k_{on}, \lambda, \delta, \mu, L^*, T^*, Y^*, Y_T, C_0^*, C_i^*, C_{N+1}^*$ | $k_{off}, \lambda, \gamma^{y/z}_{+/-}, \phi, \mu, L^*, T^*, Z_T, C_0^*, C_i^*, C_{N+1}^*$ | $\delta, Y^*, Y_T$ |
| | $R(t)$, $T_T$ | $k_{on}, k_{off}, k_p, \lambda, \gamma^{y/z}_{+/-}, \phi, \mu, L^*, Z_T, C_0^*, C_i^*, C_{N+1}^*$ | $\delta, Y^*, Y_T$ | $k_{off}, k_p, \lambda, \gamma^{y/z}_{+/-}, \phi, \mu, L^*, Z_T, C_0^*, C_i^*, C_{N+1}^*$ | $\delta, Y^*, Y_T$ | $k_{on}, k_{off}, \lambda, \gamma^{y/z}_{+/-}, \phi, \mu, L^*, Z_T, C_0^*, C_i^*, C_{N+1}^*$ | $\delta, Y^*, Y_T$ | $k_{off}, \lambda, \gamma^{y/z}_{+/-}, \phi, \mu, L^*, Z_T$ $C_0^*, C_i^*, C_{N+1}^*$ | $\delta, Y^*, Y_T$ |
| | $T_T$ | *None* | $k_{on}, k_{off}, k_p, \lambda, \gamma^{y/z}_{+/-}, \phi, \mu, \delta, L^*, Z^*, Z_T, Y^*, Y_T, C_0^*, C_i^*, C_{N+1}^*$ | *None* | $k_{off}, k_p, \lambda, \gamma^{y/z}_{+/-}, \phi, \mu, \delta, L^*, Z^*, Z_T, Y^*, Y_T, C_0^*, C_i^*, C_{N+1}^*$ | *None* | $k_{on}, k_{off}, \lambda, \gamma^{y/z}_{+/-}, \phi, \mu, \delta, L^*, Z^*, Z_T, Y^*, Y_T, C_0^*, C_i^*, C_{N+1}^*$ | *None* | $k_{off}, \lambda, \gamma^{y/z}_{+/-}, \phi, \mu, \delta, L^*, Z^*, Z_T, Y^*, Y_T, C_0^*, C_i^*, C_{N+1}^*$ |

**Table S1 (cont.):** Identifiability results with measurements of lymphocyte concentration and T-cell response $R(t)$. Variables with an asterisk (*) denote initial conditions. "Id", identifiable parameters and variables. "Non-id", non identifiable parameters and variables. In the "Id" columns, variables in boldface indicate structurally global identifiability (SGI). The index $i$ goes from 1 to $N$, except in the cases labeled with † in the **Known output** column, where it goes from 1 to $N-1$.

| | | **Additional known output (parameters)** | | | | | | | |
|---|---|---|---|---|---|---|---|---|---|
| | | — | | $k_{on}$ | | $k_p$ | | $k_{on}$, $k_p$ | |
| **Models** | **Known output** | Id | Non-id | Id | Non-id | Id | Non-id | Id | Non-id |
| **KPR-NF1** $N=2$ | $T(t)$ | $\boldsymbol{k_{on}}$ | $k_{off}$, $k_p$, $\alpha$, $\beta$, $b$, $\gamma$, $L^*$, $P^*$, $P_T$, $C_0^*$, $C_i^*$ | *None* | $k_{off}$, $k_p$, $\alpha$, $\beta$, $b$, $\gamma$, $L^*$, $P^*$, $P_T$, $C_0^*$, $C_i^*$ | $k_{on}$ | $k_{off}$, $\alpha$, $\beta$, $b$, $\gamma$, $L^*$, $P^*$, $P_T$, $C_0^*$, $C_i^*$ | *None* | $k_{off}$, $\alpha$, $\beta$, $b$, $\gamma$, $L^*$, $P^*$, $P_T$, $C_0^*$, $C_i^*$ |
| | $T(t)$, $R(t)$† | $\boldsymbol{k_{on}}$, $\boldsymbol{k_{off}}$, $\boldsymbol{k_p}$, $\boldsymbol{\alpha}$, $\boldsymbol{\beta}$, $\boldsymbol{b}$, $\boldsymbol{L^*}$, $\boldsymbol{C_0^*}$, $\boldsymbol{C_i^*}$ | $\gamma$, $P^*$, $P_T$ | $\boldsymbol{k_{off}}$, $\boldsymbol{k_p}$, $\boldsymbol{\alpha}$, $\boldsymbol{\beta}$, $\boldsymbol{b}$, $\boldsymbol{L^*}$, $\boldsymbol{C_0^*}$, $\boldsymbol{C_i^*}$ | $\gamma$, $P^*$, $P_T$ | $\boldsymbol{k_{on}}$, $\boldsymbol{k_{off}}$ $\boldsymbol{\alpha}$, $\boldsymbol{\beta}$, $\boldsymbol{b}$, $\boldsymbol{L^*}$, $\boldsymbol{C_0^*}$, $\boldsymbol{C_i^*}$ | $\gamma$, $P^*$, $P_T$ | $\boldsymbol{k_{off}}$, $\boldsymbol{\alpha}$, $\boldsymbol{\beta}$, $\boldsymbol{b}$, $\boldsymbol{L^*}$, $\boldsymbol{C_0^*}$, $\boldsymbol{C_i^*}$ | $\gamma$, $P^*$, $P_T$ |
| | $R(t)$† | $k_{on}$, $k_{off}$, $k_p$, $\alpha$, $\beta$, $b$, $L^*$ $T^*$, $C_0^*$, $C_i^*$ | $\gamma$, $P^*$, $P_T$ | $k_{off}$, $k_p$, $\alpha$, $\beta$, $b$, $L^*$ $T^*$, $C_0^*$, $C_i^*$ | $\gamma$, $P^*$, $P_T$ | $k_{on}$, $k_{off}$, $\alpha$, $\beta$, $b$, $L^*$ $T^*$, $C_0^*$, $C_i^*$ | $\gamma$, $P^*$, $P_T$ | $k_{off}$, $\alpha$, $\beta$, $b$, $L^*$ $T^*$, $C_0^*$, $C_i^*$ | $\gamma$, $P^*$, $P_T$ |
| | $R(t)$, $T_T$† | $k_{on}$, $k_{off}$, $k_p$, $\alpha$, $\beta$, $b$, $L^*$, $C_0^*$, $C_i^*$ | $\gamma$, $P^*$, $P_T$ | $k_{off}$, $k_p$, $\alpha$, $\beta$, $b$, $L^*$, $C_0^*$, $C_i^*$ | $\gamma$, $P^*$, $P_T$ | $k_{on}$, $k_{off}$, $\alpha$, $\beta$, $b$, $L^*$, $C_0^*$, $C_i^*$ | $\gamma$, $P^*$, $P_T$ | $k_{off}$, $\alpha$, $\beta$, $b$, $L^*$, $C_0^*$, $C_i^*$ | $\gamma$, $P^*$, $P_T$ |
| | $T_T$ | *None* | $k_{on}$, $k_{off}$, $k_p$, $\alpha$, $\beta$, $b$, $\gamma$, $L^*$, $P^*$, $P_T$, $C_0^*$, $C_i^*$ | *None* | $k_{off}$, $k_p$, $\alpha$, $\beta$, $b$, $\gamma$, $L^*$, $P^*$, $P_T$, $C_0^*$, $C_i^*$ | *None* | $k_{on}$, $k_{off}$, $\alpha$, $\beta$, $b$, $\gamma$, $L^*$, $P^*$, $P_T$, $C_0^*$, $C_i^*$ | $\boldsymbol{k_{off}}$, $\boldsymbol{L^*}$ | $\alpha$, $\beta$, $b$, $\gamma$, $L^*$, $P^*$, $P_T$, $C_0^*$, $C_i^*$ |
| **KPR-NF2** $N=2$; $R(t)=\epsilon_1 C_1+\epsilon_2 C_2$ | $T(t)$ | $\boldsymbol{k_{on}}$ | $k_{off}$, $k_p$, $\alpha$, $\beta$, $\epsilon_i$ $b$, $\gamma$, $L^*$, $P^*$, $P_T$, $C_0^*$, $C_i^*$ | *None* | $k_{off}$, $k_p$, $\alpha$, $\beta$, $\epsilon_i$ $b$, $\gamma$, $L^*$, $P^*$, $P_T$, $C_0^*$, $C_i^*$ | $k_{on}$ | $k_{off}$, $\alpha$, $\beta$, $\epsilon_i$ $b$, $\gamma$, $L^*$, $P^*$, $P_T$, $C_0^*$, $C_i^*$ | *None* | $k_{off}$, $\alpha$, $\beta$, $\epsilon_i$ $b$, $\gamma$, $L^*$, $P^*$, $P_T$, $C_0^*$, $C_i^*$ |
| | $T(t)$, $R(t)$ | $k_{on}$, $k_{off}$, $k_p$, $\alpha$, $\beta$, $b$, $\epsilon_i$, $L^*$, $C_0^*$, $C_i^*$ | $\gamma$, $P^*$, $P_T$ | $k_{off}$, $k_p$, $\alpha$, $\beta$, $b$, $\epsilon_i$, $L^*$, $C_0^*$, $C_i^*$ | $\gamma$, $P^*$, $P_T$ | $k_{on}$, $k_{off}$, $\alpha$, $\beta$, $b$, $\epsilon_i$, $L^*$, $C_0^*$, $C_i^*$ | $\gamma$, $P^*$, $P_T$ | $k_{off}$, $\alpha$, $\beta$, $b$, $\epsilon_i$, $L^*$, $C_0^*$, $C_i^*$ | $\gamma$, $P^*$, $P_T$ |
| | $R(t)$ | $k_{on}$, $k_{off}$, $k_p$, $\alpha$, $\beta$, $b$, $\epsilon_i$, $L^*$ $T^*$, $C_0^*$, $C_i^*$ | $\gamma$, $P^*$, $P_T$ | $k_{off}$, $k_p$, $\alpha$, $\beta$, $b$, $\epsilon_i$, $L^*$ $T^*$, $C_0^*$, $C_i^*$ | $\gamma$, $P^*$, $P_T$ | $k_{on}$, $k_{off}$, $\alpha$, $\beta$, $b$, $\epsilon_i$, $L^*$ $T^*$, $C_0^*$, $C_i^*$ | $\gamma$, $P^*$, $P_T$ | $k_{off}$, $\alpha$, $\beta$, $b$, $\epsilon_i$, $L^*$ $T^*$, $C_0^*$, $C_i^*$ | $\gamma$, $P^*$, $P_T$ |
| | $R(t)$, $T_T$ | $k_{on}$, $k_{off}$, $k_p$, $\alpha$, $\beta$, $b$, $\epsilon_i$ $L^*$, $C_0^*$, $C_i^*$ | $\gamma$, $P^*$, $P_T$ | $k_{off}$, $k_p$, $\alpha$, $\beta$, $b$, $\epsilon_i$ $L^*$, $C_0^*$, $C_i^*$ | $\gamma$, $P^*$, $P_T$ | $k_{on}$, $k_{off}$, $\alpha$, $\beta$, $b$, $\epsilon_i$ $L^*$, $C_0^*$, $C_i^*$ | $\gamma$, $P^*$, $P_T$ | $k_{off}$, $\alpha$, $\beta$, $b$, $\epsilon_i$, $L^*$, $C_0^*$, $C_i^*$ | $\gamma$, $P^*$, $P_T$ |
| | $T_T$ | *None* | $k_{on}$, $k_{off}$, $k_p$, $\alpha$, $\beta$, $\epsilon_i$, $b$, $\gamma$, $L^*$, $P^*$, $P_T$, $C_0^*$, $C_i^*$ | *None* | $k_{off}$, $k_p$, $\alpha$, $\beta$, $\epsilon_i$ $b$, $\gamma$, $L^*$, $P^*$, $P_T$, $C_0^*$, $C_i^*$ | *None* | $k_{on}$, $k_{off}$, $\alpha$, $\beta$, $\epsilon_i$ $b$, $\gamma$, $L^*$, $P^*$, $P_T$, $C_0^*$, $C_i^*$ | $\boldsymbol{k_{off}}$, $\boldsymbol{L^*}$ | $\alpha$, $\beta$, $b$, $\epsilon_i$ $\gamma$, $L^*$, $P^*$, $P_T$, $C_0^*$, $C_i^*$ |

**Table S1 (cont.):** Identifiability results with measurements of lymphocyte concentration and T-cell response $R(t)$. Variables with an asterisk (*) denote initial conditions. "Id", identifiable parameters and variables. "Non-id", non identifiable parameters and variables. In the "Id" columns, variables in boldface indicate structurally global identifiability (SGI). The index $i$ goes from 1 to $N$, except in the cases labeled with † in the **Known output** column, where it goes from 1 to $N-1$.

| | | Additional known output (parameters) | | | | | | | |
|---|---|---|---|---|---|---|---|---|---|
| | | — | | $k_{on}$ | | $k_p$ | | $k_{on}$, $k_p$ | |
| **Models** | **Known output** | Id | Non-id | Id | Non-id | Id | Non-id | Id | Non-id |
| **KPR-SAC** | $T(t)$ | $k_{on}, k_{off}, k_p, r, r_p, L^*, C_0^*, C_i^*$ | *None* | $\boldsymbol{k_{off}, k_p, r, r_p, L^*, C_0^*, C_i^*}$ | *None* | $\boldsymbol{k_{on}, k_{off}, r, r_p, L^*, C_0^*, C_i^*}$ | *None* | $\boldsymbol{k_{off}, r, r_p, L^*, C_0^*, C_i^*}$ | *None* |
| | $T(t)$, $R(t)$† | $\boldsymbol{k_{on}, k_{off}, k_p, r, r_p, L^*, C_0^*, C_i^*}$ | *None* | $\boldsymbol{k_{off}, k_p, r, r_p, L^*, C_0^*, C_i^*}$ | *None* | $\boldsymbol{k_{on}, k_{off}, r, r_p, L^*, C_0^*, C_i^*}$ | *None* | $\boldsymbol{k_{off}, r, r_p, L^*, C_0^*, C_i^*}$ | *None* |
| | $R(t)$† | $k_{on}, k_{off}, k_p, r, r_p, L^*, T^*, C_0^*, C_i^*$ | *None* | $\boldsymbol{k_{off}, k_p, r, r_p}$, $L^*, T^*$, $\boldsymbol{C_0^*, C_i^*}$ | *None* | $\boldsymbol{k_{on}, k_{off}, r, r_p}$, $L^*, T^*$, $\boldsymbol{C_0^*, C_i^*}$ | *None* | $\boldsymbol{k_{on}, k_{off}, r, r_p}$, $L^*, T^*$, $\boldsymbol{C_0^*, C_i^*}$ | *None* |
| | $R(t)$, $T_T$† | $\boldsymbol{k_{on}, k_{off}, k_p, r, r_p, L^*, C_0^*, C_i^*}$ | *None* | $\boldsymbol{k_{off}, k_p, r, r_p, L^*, C_0^*, C_i^*}$ | *None* | $\boldsymbol{k_{on}, k_{off}, r, r_p, L^*, C_0^*, C_i^*}$ | *None* | $\boldsymbol{k_{off}, r, r_p, L^*, C_0^*, C_i^*}$ | *None* |
| | $T_T$ | *None* | $k_{on}, k_{off}, k_p, r, r_p, L^*, C_0^*, C_i^*$ | *None* | $k_{off}, k_p, r, r_p, L^*, C_0^*, C_i^*$ | *None* | $k_{on}, k_{off}, r, r_p, L^*, C_0^*, C_i^*$ | *None* | $k_{off}, r, r_p, L^*, C_0^*, C_i^*$ |
| **ZU** | $T(t)$ | $k_{on}, k_{off}, k_+, k_-, \lambda, k_p, L^*, D^*, P^*, C_0^*, T_p^*$ | *None* | $\boldsymbol{k_{off}, k_+, k_-, \lambda, k_p, L^*, D^*, P^*, C_0^*, T_p^*}$ | *None* | $k_{on}, k_{off}, k_+, k_-, \lambda, L^*, D^*, P^*, C_0^*, T_p^*$ | *None* | $\boldsymbol{k_{off}, k_+, k_-, \lambda, L^*, D^*, P^*, C_0^*, T_p^*}$ | *None* |
| | $T(t)$, $R(t)$ | $\boldsymbol{k_{on}, k_{off}, k_+, k_-, \lambda, k_p, L^*, D^*, P^*, C_0^*}$ | *None* | $\boldsymbol{k_{off}, k_+, k_-, \lambda, k_p, L^*, D^*, P^*, C_0^*}$ | *None* | $\boldsymbol{k_{on}, k_{off}, k_+, k_-, \lambda, L^*, D^*, P^*, C_0^*}$ | *None* | $\boldsymbol{k_{off}, k_+, k_-, \lambda, L^*, D^*, P^*, C_0^*}$ | *None* |
| | $R(t)$ | $k_{on}, k_{off}, k_+, k_-, \lambda, k_p, L^*, T^*, D^*, P^*, C_0^*$ | *None* | $k_{off}, k_+, k_-, \lambda, k_p, L^*, T^*, D^*, P^*, C_0^*$ | *None* | $k_{on}, k_{off}, k_+, k_-, \lambda$ $L^*, T^*, D^*, P^*, C_0^*$ | *None* | $k_{off}, k_+, k_-, \lambda, L^*, T^*, D^*, P^*, C_0^*$ | *None* |
| | $R(t)$, $T_T$ | $k_{on}, k_{off}, k_+, k_-, k_p, \lambda, L^*, D^*, P^*\ C_0^*$ | *None* | $k_{off}, k_+, k_-, k_p, \lambda, L^*, D^*, P^*\ C_0^*$ | *None* | $k_{on}, k_{off}, k_+, k_-, \lambda, L^*, D^*, P^*\ C_0^*$ | *None* | $k_{off}, k_+, k_-, \lambda, L^*, D^*, P^*\ C_0^*$ | *None* |
| | $T_T$ | *None* | $k_{on}, k_{off}, k_+, k_-, \lambda, k_p, L^*, D^*, P^*, C_0^*, T_p^*$ | *None* | $k_{off}, k_+, k_-, \lambda, k_p, L^*, D^*, P^*, C_0^*, T_p^*$ | *None* | $k_{on}, k_{off}, k_+, k_-, \lambda, L^*, D^*, P^*, C_0^*, T_p^*$ | *None* | $k_{off}, k_+, k_-, \lambda, L^*, D^*, P^*, C_0^*, T_p^*$ |
| **KPZU** | $T(t)$ | $k_{on}, k_{off}, k_p, k_+, k_-, \lambda, L^*, D^*, P^*, C_0^*, C_i^*, T_p^*$ | *None* | $k_{off}, k_p, k_+, k_-, \lambda, L^*, D^*, P^*, C_0^*, C_i^*, T_p^*$ | *None* | $k_{on}, k_{off}, k_+, k_-, \lambda, L^*, D^*, P^*, C_0^*, C_i^*, T_p^*$ | *None* | $k_{off}, k_+, k_-, \lambda, L^*, D^*, P^*, C_0^*, C_i^*, T_p^*$ | *None* |
| | $T(t)$, $R(t)$ | $k_{on}, k_{off}, k_p, k_+, k_-, \lambda, L^*, D^*, P^*, C_0^*, C_i^*$ | *None* | $k_{off}, k_p, k_+, k_-, \lambda, L^*, D^*, P^*, C_0^*, C_i^*$ | *None* | $k_{on}, k_{off}, k_+, k_-, \lambda, L^*, D^*, P^*, C_0^*, C_i^*$ | *None* | $k_{off}, k_+, k_-, \lambda, L^*, D^*, P^*, C_0^*, C_i^*$ | *None* |
| | $R(t)$ | $k_{on}, k_{off}, k_p, k_+, k_-, \lambda, L^*, T^*, D^*, P^*, C_0^*, C_i^*$ | *None* | $k_{off}, k_p, k_+, k_-, \lambda, L^*, T^*, D^*, P^*, C_0^*, C_i^*$ | *None* | $k_{on}, k_{off}, k_+, k_-, \lambda, L^*, T^*, D^*, P^*, C_0^*, C_i^*$ | *None* | $k_{on}, k_{off}, k_p, k_+, k_-, \lambda, L^*, T^*, D^*, P^*, C_0^*, C_i^*$ | *None* |
| | $R(t)$, $T_T$ | $k_{on}, k_{off}, k_p, k_+, k_-, \lambda, L^*, D^*, P^*, C_0^*, C_i^*$ | *None* | $k_{on}, k_{off}, k_p, k_+, k_-, \lambda, L^*, D^*, P^*, C_0^*, C_i^*$ | *None* | $k_{on}, k_{off}, k_p, k_+, k_-, \lambda, L^*, D^*, P^*, C_0^*, C_i^*$ | *None* | $k_{off}, k_+, k_-, \lambda, L^*, D^*, P^*, C_0^*, C_i^*$ | *None* |
| | $T_T$ | *None* | $k_{on}, k_{off}, k_p, k_+, k_-, \lambda, L^*, D^*, P^*, C_0^*, C_i^*, T_p^*$ | *None* | $k_{off}, k_p, k_+, k_-, \lambda, L^*, D^*, P^*, C_0^*, C_i^*, T_p^*$ | *None* | $k_{on}, k_{off}, k_+, k_-, \lambda, L^*, D^*, P^*, C_0^*, C_i^*, T_p^*$ | *None* | $k_{off}, k_+, k_-, \lambda, L^*, D^*, P^*, C_0^*, C_i^*, T_p^*$ |

**Table S1 (cont.):** Identifiability results with measurements of lymphocyte concentration and T-cell response $R(t)$. Variables with an asterisk (*) denote initial conditions. "Id", identifiable parameters and variables. "Non-id", non identifiable parameters and variables. In the "Id" columns, variables in boldface indicate structurally global identifiability (SGI). The index $i$ goes from 1 to $N$.

| | | **Additional known output (parameters)** | | | | | | | |
|---|---|---|---|---|---|---|---|---|---|
| | | — | | $k_{on}$ | | $k_p$ | | $k_{on}$, $k_p$ | |
| **Models** | **Known output** | Id | Non-id | Id | Non-id | Id | Non-id | Id | Non-id |
| **KPC** | $T(t)$ | $k_{on}$, $k_{off}$, $k_p$, $k_+$, $k_-$, $\lambda$, $D^*$, $X^*$, $C_0^*$, $C_i^*$, $T_p^*$ | *None* | $k_{off}$, $k_p$, $k_+$, $k_-$, $\lambda$, $D^*$, $X^*$, $C_0^*$, $C_i^*$, $T_p^*$ | *None* | $k_{on}$, $k_{off}$, $k_+$, $k_-$, $\lambda$, $D^*$, $X^*$, $C_0^*$, $C_i^*$, $T_p^*$ | *None* | $k_{off}$, $k_+$, $k_-$, $\lambda$, $D^*$, $X^*$, $C_0^*$, $C_i^*$, $T_p^*$ | *None* |
| | $T(t)$, $R(t)$ | $\boldsymbol{k_{on}, k_{off}, k_p, k_+, k_-, \lambda, D^*, X^*, C_0^*, C_i^*}$ | *None* | $\boldsymbol{k_{off}, k_p, k_+, k_-, \lambda, D^*, X^*, C_0^*, C_i^*}$ | *None* | $\boldsymbol{k_{on}, k_{off}, k_+, k_-, \lambda, D^*, X^*, C_0^*, C_i^*}$ | *None* | $\boldsymbol{k_{off}, k_+, k_-, \lambda, D^*, X^*, C_0^*, C_i^*}$ | *None* |
| | $R(t)$ | $k_{on}$, $k_{off}$, $k_p$, $k_+$, $k_-$, $\lambda$, $T^*$, $D^*$, $X^*$, $C_0^*$, $C_i^*$ | *None* | $k_{off}$, $k_p$, $k_+$, $k_-$, $\lambda$, $T^*$, $D^*$, $X^*$, $C_0^*$, $C_i^*$ | *None* | $k_{on}$, $k_{off}$, $k_+$, $k_-$, $\lambda$, $T^*$, $D^*$, $X^*$, $C_0^*$, $C_i^*$ | *None* | $k_{off}$, $k_+$, $k_-$, $\lambda$, $T^*$, $D^*$, $X^*$, $C_0^*$, $C_i^*$ | *None* |
| | $R(t)$, $T_T$ | $k_{on}$, $k_{off}$, $k_p$, $k_+$, $k_-$, $\lambda$, $D^*$, $X^*$, $C_0^*$, $C_i^*$ | *None* | $k_{off}$, $k_p$, $k_+$, $k_-$, $\lambda$, $D^*$, $X^*$, $C_0^*$, $C_i^*$ | *None* | $k_{on}$, $k_{off}$, $k_+$, $k_-$, $\lambda$, $D^*$, $X^*$, $C_0^*$, $C_i^*$ | *None* | $k_{off}$, $k_+$, $k_-$, $\lambda$, $D^*$, $X^*$, $C_0^*$, $C_i^*$ | *None* |
| | $T_T$ | *None* | $k_{on}$, $k_{off}$, $k_p$, $k_+$, $k_-$, $\lambda$, $D^*$, $X^*$, $C_0^*$, $C_i^*$, $T_p^*$ | *None* | $k_{off}$, $k_p$, $k_+$, $k_-$, $\lambda$, $D^*$, $X^*$, $C_0^*$, $C_i^*$, $T_p^*$ | *None* | $k_{on}$, $k_{off}$, $k_+$, $k_-$, $\lambda$, $D^*$, $X^*$, $C_0^*$, $C_i^*$ | *None* | $k_{off}$, $k_+$, $k_-$, $\lambda$, $D^*$, $X^*$, $C_0^*$, $C_i^*$, $T_p^*$ |

| | | — | | $k_{eff}$ | |
|---|---|---|---|---|---|
| | | Id | Non-id | Id | Non-id |
| **ST** $N = 2$ | $T(t)$ | $\lambda$, $\phi$, $\sigma$, $S^*$ | $k_{eff}$, $k_i$, $T_p^*$, $L$ | $\lambda$, $\phi$, $\sigma$, $S^*$, $L$ | $k_i$, $T_p^*$ |
| | $T(t)$, $R(t)$ | $\lambda$, $\phi$, $\sigma$, $k_i$, $S^*$ | $k_{eff}$, $L$ | $\lambda$, $\phi$, $\sigma$, $k_i$, $S^*$, $L$ | *None* |
| | $R(t)$ | $\lambda$, $\phi$, $\sigma$, $k_i$, $S^*$, $T^*$ | $k_{eff}$, $L$ | $\lambda$, $\phi$, $\sigma$, $k_i$, $S^*$, $T^*$, $L$ | *None* |
| | $R(t)$, $T_T$ | $\lambda$, $\phi$, $\sigma$, $k_i$, $S^*$ | $k_{eff}$, $L$ | $\lambda$, $\phi$, $\sigma$, $k_i$, $S^*$, $L$ | *None* |
| | $T_T$ | $\lambda$, $\phi$, $\sigma$, $k_i$, $S^*$, $T_p^*$ | $k_{eff}$, $L$ | $\lambda$, $\phi$, $\sigma$, $k_i$, $S^*$, $L$, $T_p^*$ | *None* |

**Table S2:** Identifiability results with measurements of $EC_{50}$ and $E_{max}$. Variables with an asterisk (*) denote initial conditions. In the "Id" columns, variables in boldface indicate structurally global identifiability (SGI). The index $i$ goes from 1 to $N$.

| Models | Additional known output | Known output: $EC_{50}$ Id | $EC_{50}$ Non-id | $E_{max}$ Id | $E_{max}$ Non-id |
|---|---|---|---|---|---|
| **Occ** | — | $k_{on}$, $k_{off}$, $T^*$, $L^*$, $C_0^*$ | *None* | *None* | $k_{on}$, $k_{off}$, $T^*$, $L^*$, $C_0^*$ |
| | $T(t)$ | $\boldsymbol{k_{on}, k_{off}, L^*, C_0^*}$ | *None* | $\boldsymbol{k_{on}, k_{off}, L^*, C_0^*}$ | *None* |
| | $k_{on}$ | $k_{off}$, $T^*$, $L^*$, $C_0^*$ | *None* | $\boldsymbol{k_{off}}$ | $T^*$, $L^*$, $C_0^*$ |
| | $T(t)$, $k_{on}$ | $\boldsymbol{k_{off}, L^*, C_0^*}$ | *None* | $\boldsymbol{k_{off}, L^*, C_0^*}$ | *None* |
| | $T_T$ | $\boldsymbol{k_{on}, k_{off}, L^*, C_0^*}$ | *None* | *None* | $k_{on}$, $k_{off}$, $L^*$, $C_0^*$ |
| | $T_T$, $k_{on}$ | $\boldsymbol{k_{off}, L^*, C_0^*}$ | *None* | $\boldsymbol{k_{off}, L^*, C_0^*}$ | *None* |
| **KPR-1** | — | *None* | $k_{on}$, $k_{off}$, $k_p$, $L^*$, $T^*$, $C_0^*$, $C_i^*$ | *None* | $k_{on}$, $k_{off}$, $k_p$, $L^*$, $T^*$, $C_0^*$, $C_i^*$ |
| | $T(t)$ | $k_{off}$, $\boldsymbol{k_{on}}$, $L^*$ | $k_p$, $C_0^*$, $C_i^*$ | $k_{on}$ | $k_{off}$, $k_p$, $L^*$, $C_0^*$, $C_i^*$ |
| | $k_{on}$ | *None* | $k_{off}$, $k_p$, $L^*$, $T^*$, $C_0^*$, $C_i^*$ | *None* | $k_{off}$, $k_p$, $L^*$, $T^*$, $C_0^*$, $C_i^*$ |
| | $k_p$ | *None* | $k_{on}$, $k_{off}$, $L^*$, $T^*$, $C_0^*$, $C_i^*$ | *None* | $k_{on}$, $k_{off}$, $L^*$, $T^*$, $C_0^*$, $C_i^*$ |
| | $T(t)$, $k_{on}$, $k_p$ | $k_{off}$, $L^*$ | $C_0^*$, $C_i^*$ | $k_{off}$, $L^*$ | $C_0^*$, $C_i^*$ |
| | $T_T$ | *None* | $k_{on}$, $k_{off}$, $k_p$, $L^*$, $T^*$, $C_0^*$, $C_i^*$ | *None* | $k_{on}$, $k_{off}$, $k_p$, $L^*$, $C_0^*$, $C_i^*$ |
| | $T_T$, $k_{on}$, $k_p$ | $\boldsymbol{k_{off}}$ | $L^*$, $C_0^*$, $C_i^*$ | $\boldsymbol{k_{off}}$ | $L^*$, $T^*$, $C_0^*$, $C_i^*$ |
| **KPR-LS** | — | *None* | $k_{on}$, $k_{off}$, $k_p$, $\phi$, $L^*$, $T^*$, $C_0^*$, $C_i^*$, $C_{N+1}^*$ | *None* | $k_{on}$, $k_{off}$, $k_p$, $L^*$, $T^*$, $\phi$, $C_0^*$, $C_i^*$, $C_{N+1}^*$ |
| | $T(t)$ | $k_{off}$, $\boldsymbol{k_{on}}$, $L^*$ | $k_p$, $\phi$, $C_0^*$, $C_i^*$, $C_{N+1}^*$ | *None* | $k_{on}$, $k_{off}$, $k_p$, $\phi$, $L^*$, $C_0^*$, $C_i^*$, $C_{N+1}^*$ |
| | $k_{on}$ | *None* | $k_{off}$, $k_p$, $\phi$, $L^*$, $T^*$, $C_0^*$, $C_i^*$, $C_{N+1}^*$ | *None* | $k_{off}$, $k_p$, $\phi$, $L^*$, $T^*$, $C_0^*$, $C_i^*$, $C_{N+1}^*$ |
| | $k_p$ | *None* | $k_{on}$, $k_{off}$, $\phi$, $L^*$, $T^*$, $C_0^*$, $C_i^*$, $C_{N+1}^*$ | *None* | $k_{on}$, $k_{off}$, $\phi$, $L^*$, $T^*$, $C_0^*$, $C_i^*$, $C_{N+1}^*$ |
| | $T(t)$, $k_{on}$, $k_p$ | $k_{off}$, $L^*$ | $\phi$, $C_0^*$, $C_i^*$, $C_{N+1}^*$ | *None* | $k_{off}$, $\phi$, $L^*$, $C_0^*$, $C_i^*$, $C_{N+1}^*$ |
| | $T_T$ | *None* | $k_{on}$, $k_{off}$, $k_p$, $\phi$, $L^*$, $C_0^*$, $C_i^*$, $C_{N+1}^*$ | *None* | $k_{on}$, $k_{off}$, $k_p$, $\phi$, $L^*$, $C_0^*$, $C_i^*$, $C_{N+1}^*$ |
| | $T_T$, $k_{on}$, $k_p$ | $\boldsymbol{k_{off}}$ | $\phi$, $L^*$, $C_0^*$, $C_i^*$, $C_{N+1}^*$ | *None* | $k_{off}$, $\phi$, $L^*$, $C_0^*$, $C_i^*$, $C_{N+1}^*$ |
| **KPR-SS** | — | $k_{on}$, $k_{off}$, $k_p$, $\lambda$, $L^*$, $T^*$, $C_0^*$, $C_i^*$, $T_N^*$ | *None* | *None* | $k_{on}$, $k_{off}$, $k_p$, $\lambda$, $L^*$, $T^*$, $C_0^*$, $C_i^*$, $T_N^*$ |
| | $T(t)$ | $k_{on}$, $k_{off}$, $k_p$, $\lambda$, $L^*$, $C_0^*$, $C_i^*$, $T_N^*$ | *None* | $k_{off}$, $k_{on}$, $k_p$, $\lambda$, $L^*$, $C_0^*$, $C_i^*$, $T_N^*$ | *None* |
| | $k_{on}$ | $k_{off}$, $k_p$, $\lambda$, $L^*$, $T^*$, $C_0^*$, $C_i^*$, $T_N^*$ | *None* | *None* | $k_{off}$, $k_p$, $\lambda$, $L^*$, $T^*$, $C_0^*$, $C_i^*$, $T_N^*$ |
| | $k_p$ | $k_{on}$, $k_{off}$, $\lambda$, $L^*$, $T^*$, $C_0^*$, $C_i^*$, $T_N^*$ | *None* | *None* | $k_{on}$, $k_{off}$, $\lambda$, $L^*$, $T^*$, $C_0^*$, $C_i^*$, $T_N^*$ |
| | $T(t)$, $k_{on}$, $k_p$ | $k_{off}$, $\lambda$, $L^*$, $C_0^*$, $C_i^*$, $T_N^*$ | *None* | $\boldsymbol{k_{off}, \lambda, L^*, C_0^*, C_i^*, T_N^*}$ | *None* |
| | $T_T$ | $k_{on}$, $k_{off}$, $k_p$, $\lambda$, $L^*$, $C_0^*$, $C_i^*$, $T_N^*$ | *None* | *None* | $k_{on}$, $k_{off}$, $k_p$, $\lambda$, $L^*$, $T^*$, $C_0^*$, $C_i^*$, $T_N^*$ |
| | $T_T$, $k_{on}$, $k_p$ | $k_{off}$, $\lambda$, $L^*$, $C_0^*$, $C_i^*$, $T_N^*$ | *None* | — | $k_{off}$, $\lambda$, $L^*$, $T^*$, $C_0^*$, $C_i^*$, $T_N^*$ |

**Table S2 (cont.):** Identifiability results with measurements of $EC_{50}$ and $E_{max}$. Variables with an asterisk (*) denote initial conditions. In the "Id" columns, variables in boldface indicate structural global identifiability (SGI). The index $i$ goes from 1 to $N$.

| Models | Additional known output | Known output: $EC_{50}$ Id | $EC_{50}$ Non-id | $E_{max}$ Id | $E_{max}$ Non-id |
|---|---|---|---|---|---|
| **KPR-IR** | — | $k_{on}$, $k_{off}$, $k_p$, $\rho_1$, $\rho_i$, $\lambda$, $T^*$, $L^*$, $\overline{C}_0^*$, $\overline{C}_i^*$ | *None* | $k_{on}$, $k_{off}$, $k_p$, $\rho_1$, $\rho_i$, $\lambda$, $T^*$, $L^*$, $\overline{C}_0^*$, $\overline{C}_i^*$ | *None* |
| | $T(t)$ | $k_{on}$, $k_{off}$, $k_p$, $\rho_1$, $\rho_i$, $\lambda$, $L^*$, $\overline{C}_0^*$, $\overline{C}_i^*$ | *None* | $k_{on}$, $k_{off}$, $k_p$, $\rho_1$, $\rho_i$, $\lambda$, $L^*$, $\overline{C}_0^*$, $\overline{C}_i^*$ | *None* |
| | $k_{on}$ | $k_{off}$, $k_p$, $\rho_1$, $\rho_i$, $\lambda$, $T^*$, $L^*$, $\overline{C}_0^*$, $\overline{C}_i^*$ | *None* | $k_{off}$, $k_p$, $\rho_1$, $\rho_i$, $\lambda$, $T^*$, $L^*$, $\overline{C}_0^*$, $\overline{C}_i^*$ | *None* |
| | $k_p$ | $k_{on}$, $k_{off}$, $\rho_1$, $\rho_i$, $\lambda$, $T^*$, $L^*$, $\overline{C}_0^*$, $\overline{C}_i^*$ | *None* | $\boldsymbol{k_{on}}$, $\boldsymbol{k_{off}}$, $\boldsymbol{\rho_1}$, $\boldsymbol{\rho_i}$, $\boldsymbol{\lambda}$, $\boldsymbol{T^*}$, $\boldsymbol{L^*}$, $\boldsymbol{\overline{C}_0^*}$, $\boldsymbol{\overline{C}_i^*}$ | *None* |
| | $T(t)$, $k_{on}$, $k_p$ | $\boldsymbol{k_{off}}$, $\boldsymbol{\rho_1}$, $\boldsymbol{\rho_i}$, $\boldsymbol{\lambda}$, $\boldsymbol{L^*}$, $\boldsymbol{\overline{C}_0^*}$, $\boldsymbol{\overline{C}_i^*}$ | *None* | $k_{off}$, $\rho_1$, $\rho_i$, $\lambda$, $L^*$, $\overline{C}_0^*$, $\overline{C}_i^*$ | *None* |
| | $T_T$ | $k_{on}$, $k_{off}$, $k_p$, $\rho_1$, $\rho_i$, $\lambda$, $L^*$, $\overline{C}_0^*$, $\overline{C}_i^*$ | *None* | $\boldsymbol{k_{on}}$, $\boldsymbol{k_{off}}$, $\boldsymbol{k_p}$, $\boldsymbol{\rho_1}$, $\boldsymbol{\rho_i}$, $\boldsymbol{\lambda}$, $\boldsymbol{L^*}$, $\boldsymbol{\overline{C}_0^*}$, $\boldsymbol{\overline{C}_i^*}$ | *None* |
| | $T_T$, $k_{on}$, $k_p$ | $\boldsymbol{k_{off}}$, $\boldsymbol{\rho_1}$, $\boldsymbol{\rho_i}$, $\boldsymbol{\lambda}$, $\boldsymbol{L^*}$, $\boldsymbol{\overline{C}_0^*}$, $\boldsymbol{\overline{C}_i^*}$ | *None* | $\boldsymbol{k_{off}}$, $\boldsymbol{\rho_1}$, $\boldsymbol{\rho_i}$, $\boldsymbol{\lambda}$, $\boldsymbol{L^*}$, $\boldsymbol{\overline{C}_0^*}$, $\boldsymbol{\overline{C}_i^*}$ | *None* |
| **KPR-LS-IFF** | — | *None* | $k_{on}$, $k_{off}$, $k_p$, $\lambda$, $\gamma^{y/z}_{+/-}$, $\phi$, $\mu$, $\delta$, $T^*$, $L^*$, $Y^*$, $Y_T$, $C_0^*$, $C_i^*$, $C_{N+1}^*$, $Z^*$, $Z_T$ | $k_{on}$, $Z^*$, $Z_T$, $\delta$ | $k_{off}$, $k_p$, $\lambda$, $\gamma^{y/z}_{+/-}$, $\phi$, $\mu$, $T^*$, $L^*$, $Y^*$, $Y_T$, $C_0^*$, $C_i^*$, $C_{N+1}^*$ |
| | $T(t)$ | *None* | $k_{on}$, $k_{off}$, $k_p$, $\lambda$, $\gamma^{y/z}_{+/-}$, $\phi$, $\mu$, $\delta$, $L^*$, $Y^*$, $Y_T$, $C_0^*$, $C_i^*$, $C_{N+1}^*$, $Z^*$, $Z_T$ | $k_{on}$, $Z^*$, $Z_T$, $\delta$ | $k_{off}$, $k_p$, $\lambda$, $\gamma^{y/z}_{+/-}$, $\phi$, $\mu$, $L^*$, $Y^*$, $Y_T$, $C_0^*$, $C_i^*$, $C_{N+1}^*$ |
| | $k_{on}$ | *None* | $k_{off}$, $k_p$, $\lambda$, $\gamma^{y/z}_{+/-}$, $\phi$, $\mu$, $\delta$, , $T^*$, $L^*$, $Y^*$, $Y_T$, $C_0^*$, $C_i^*$, $C_{N+1}^*$, $Z^*$, $Z_T$ | $Z^*$, $Z_T$, $\delta$ | $k_{off}$, $k_p$, $\lambda$, $\gamma^{y/z}_{+/-}$, $\phi$, $\mu$, $T^*$, $L^*$, $Y^*$, $Y_T$, $C_0^*$, $C_i^*$, $C_{N+1}^*$ |
| | $k_p$ | *None* | $k_{on}$, $k_{off}$, $\lambda$, $\gamma^{y/z}_{+/-}$, $\phi$, $\mu$, $\delta$, $T^*$, $L^*$, $Y^*$, $Y_T$, $C_0^*$, $C_i^*$, $C_{N+1}^*$, $Z^*$, $Z_T$ | $k_{on}$, $k_{off}$, $\lambda$, $\phi$, $\mu$, $\delta$, $Z^*$, $Z_T$, $C_{N+1}^*$ | $\gamma^{y/z}_{+/-}$, $Y^*$, $Y_T$, $L^*$, $T^*$ , $C_0^*$, $C_i^*$ |
| | $T(t)$, $k_{on}$, $k_p$ | *None* | $k_{off}$, $\lambda$, $\gamma^{y/z}_{+/-}$, $\phi$, $\mu$, $\delta$, $L^*$, $Y^*$, $Y_T$, $C_0^*$, $C_i^*$, $C_{N+1}^*$, $Z^*$, $Z_T$ | $k_{off}$, $\lambda$, $\phi$, $\mu$, $\delta$, $Z^*$, $Z_T$, $C_i^*$, $C_{N+1}^*$ | $\gamma^{y/z}_{+/-}$, $L^*$, $Y^*$, $Y_T$, $C_0^*$ |
| | $T_T$ | *None* | $k_{on}$, $k_{off}$, $k_p$, $\lambda$, $\gamma^{y/z}_{+/-}$, $\phi$, $\mu$, $\delta$ $L^*$, $Y^*$, $Y_T$, $C_0^*$, $C_i^*$, $C_{N+1}^*$, $Z^*$, $Z_T$ | $k_{on}$, $Z^*$, $Z_T$, $\delta$ | $k_{off}$, $k_p$, $\lambda$, $\gamma^{y/z}_{+/-}$, $\phi$, $\mu$, $L^*$, $Y^*$, $Y_T$, $C_0^*$, $C_i^*$, $C_{N+1}^*$ |
| | $T_T$, $k_{on}$, $k_p$ | *None* | $k_{off}$, $\lambda$, $\gamma^{y/z}_{+/-}$, $\phi$, $\mu$, $\delta$, $L^*$, $Y^*$, $Y_T$, $C_0^*$, $C_i^*$, $C_{N+1}^*$, $Z^*$, $Z_T$ | $k_{off}$, $\lambda$, $\phi$, $\mu$, $\delta$, $Z^*$, $Z_T$, $C_i^*$, $C_{N+1}^*$ | $\gamma^{y/z}_{+/-}$, $L^*$, $Y^*$, $Y_T$, $C_0^*$ |

**Table S2 (cont.):** Identifiability results with measurements of $EC_{50}$ and $E_{max}$. Variables with an asterisk (*) denote initial conditions. In the "Id" columns, variables in boldface indicate structural global identifiability (SGI). The index $i$ goes from 1 to $N$.

| | | **Known output** | | | |
|---|---|---|---|---|---|
| | **Additional known output** | $EC_{50}$ | | $E_{max}$ | |
| **Models** | | Id | Non-id | Id | Non-id |
| **KPR-NF1** $N = 2$ | — | *None* | $k_{on}$, $k_{off}$, $k_p$, $\alpha$, $\beta$, $b$, $\gamma$, $L^*$, $T^*$, $C_0^*$, $C_i^*$, $P^*$, $P_T$ | *None* | $k_{on}$, $k_{off}$, $k_p$, $\alpha$, $\beta$, $b$, $\gamma$, $L^*$, $T^*$, $C_0^*$, $C_i^*$, $P^*$, $P_T$ |
| | $T(t)$ | $k_{on}$, $k_{off}$, $L^*$ | $k_p$, $\alpha$, $\beta$, $b$, $\gamma$, $C_0^*$, $C_i^*$, $P^*$, $P_T$ | $k_{on}$ | $k_{off}$, $k_p$, $\alpha$, $\beta$, $b$, $\gamma$, $L^*$, $C_0^*$, $C_i^*$, $P^*$, $P_T$ |
| | $k_{on}$ | *None* | $k_p$, $\alpha$, $\beta$, $b$, $\gamma$, $L^*$, $T^*$, $C_0^*$, $C_i^*$, $P^*$, $P_T$ | *None* | $k_p$, $\alpha$, $\beta$, $b$, $\gamma$, $L^*$, $T^*$, $C_0^*$, $C_i^*$, $P^*$, $P_T$ |
| | $k_p$ | *None* | $k_{on}$, $k_{off}$, $\alpha$, $\beta$, $b$, $\gamma$, $L^*$, $T^*$, $C_0^*$, $C_i^*$, $P^*$, $P_T$ | *None* | $k_{on}$, $k_{off}$, $\alpha$, $\beta$, $b$, $\gamma$, $L^*$, $T^*$, $C_0^*$, $C_i^*$, $P^*$, $P_T$ |
| | $T(t)$, $k_{on}$, $k_p$ | $k_{off}$, $L^*$ | $\alpha$, $\beta$, $b$, $\gamma$, $C_0^*$, $C_i^*$, $P^*$, $P_T$ | $k_{off}$, $b$, $\gamma$, $L^*$ | $\alpha$, $\beta$, $C_0^*$, $C_i^*$, $P^*$, $P_T$ |
| | $T_T$ | *None* | $k_{on}$, $k_p$, $\alpha$, $\beta$, $b$, $\gamma$, $L^*$, $C_0^*$, $C_i^*$, $P^*$, $P_T$ | *None* | $k_{on}$, $k_p$, $\alpha$, $\beta$, $b$, $\gamma$, $L^*$, $C_0^*$, $C_i^*$, $P^*$, $P_T$ |
| | $T_T$, $k_{on}$, $k_p$ | $\boldsymbol{k_{off}}$ | $\alpha$, $\beta$, $b$, $\gamma$, $L^*$, $C_0^*$, $C_i^*$, $P^*$, $P_T$ | $k_{off}$, $b$, $\gamma$, $L^*$ | $\alpha$, $\beta$, $C_0^*$, $C_i^*$, $P^*$, $P_T$ |
| **KPR-NF2** $N = 2$; $R(t) = \epsilon_0 C_0 + \epsilon_1 C_1 + \epsilon_2 C_2$ | — | *None* | $k_{on}$, $k_{off}$, $k_p$, $\alpha$, $\beta$, $b$, $\gamma$, $\epsilon_i$, $L^*$, $T^*$, $C_0^*$, $C_i^*$, $P^*$, $P_T$ | *None* | $k_{on}$, $k_{off}$, $k_p$, $\alpha$, $\beta$, $b$, $\gamma$, $\epsilon_i$, $L^*$, $T^*$, $C_0^*$, $C_i^*$, $P^*$, $P_T$ |
| | $T(t)$ | $k_{on}$, $k_{off}$, $L^*$ | $k_p$, $\alpha$, $\beta$, $b$, $\gamma$, $\epsilon_i$, $C_0^*$, $C_i^*$, $P^*$, $P_T$ | $k_{on}$, $\epsilon_i$ | $k_{off}$, $k_p$, $\alpha$, $\beta$, $b$, $\gamma$, $L^*$, $C_0^*$, $C_i^*$, $P^*$, $P_T$ |
| | $k_{on}$ | *None* | $k_{off}$, $k_p$, $\alpha$, $\beta$, $b$, $\gamma$, $\epsilon_i$, $L^*$, $T^*$, $C_0^*$, $C_i^*$, $P^*$, $P_T$ | *None* | $k_{off}$, $k_p$, $\alpha$, $\beta$, $b$, $\gamma$, $\epsilon_i$, $L^*$, $T^*$, $C_0^*$, $C_i^*$, $P^*$, $P_T$ |
| | $k_p$ | *None* | $k_{on}$, $k_{off}$, $\alpha$, $\beta$, $b$, $\gamma$, $\epsilon_i$, $L^*$, $T^*$, $C_0^*$, $C_i^*$, $P^*$, $P_T$ | $k_{off}$ | $k_{on}$, $\alpha$, $\beta$, $b$, $\gamma$, $\epsilon_i$, $L^*$, $T^*$, $C_0^*$, $C_i^*$, $P^*$, $P_T$ |
| | $T(t)$, $k_{on}$, $k_p$ | $k_{off}$, $L^*$ | $\alpha$, $\beta$, $b$, $\gamma$, $\epsilon_i$, $C_0^*$, $C_i^*$, $P^*$, $P_T$ | $k_{off}$, $b$, $\gamma$, $\epsilon_i$, $L^*$ | $\alpha$, $\beta$, $C_0^*$, $C_i^*$, $P^*$, $P_T$ |
| | $T_T$ | *None* | $k_{on}$, $k_{off}$, $k_p$, $\alpha$, $\beta$, $b$, $\gamma$, $\epsilon_i$, $L^*$, $C_0^*$, $C_i^*$, $P^*$, $P_T$ | *None* | $k_{on}$, $k_{off}$, $k_p$, $\alpha$, $\beta$, $b$, $\gamma$, $\epsilon_i$, $L^*$, $C_0^*$, $C_i^*$, $P^*$, $P_T$ |
| | $T_T$, $k_{on}$, $k_p$ | $\boldsymbol{k_{off}}$ | $\alpha$, $\beta$, $b$, $\gamma$, $\epsilon_i$, $L^*$, $C_0^*$, $C_i^*$, $P^*$, $P_T$ | $k_{off}$, $b$, $\gamma$, $\epsilon_i$, $L^*$ | $\alpha$, $\beta$, $C_0^*$, $C_i^*$, $P^*$, $P_T$ |

**Table S2 (cont.):** Identifiability results with measurements of $EC_{50}$ and $E_{max}$. Variables with an asterisk (*) denote initial conditions. In the "Id" columns, variables in boldface indicate structural global identifiability (SGI). The index $i$ goes from 1 to $N$.

| Models | Additional known output | Known output: $EC_{50}$ Id | $EC_{50}$ Non-id | $E_{max}$ Id | $E_{max}$ Non-id |
|---|---|---|---|---|---|
| **KPR-SAC** | — | *None* | $k_{on}$, $k_{off}$, $k_p$, $r$, $r_p$, $T^*$, $L^*$, $C_0^*$, $C_i^*$ | *None* | $k_{on}$, $k_{off}$, $k_p$, $r$, $r_p$, $T^*$, $L^*$, $C_0^*$, $C_i^*$ |
| | $T(t)$ | $k_{on}$, $k_{off}$, $k_p$, $r$, $r_p$, $L^*$, $C_0^*$, $C_i^*$ | *None* | $k_{on}$, $k_{off}$, $k_p$, $r$, $r_p$, $L^*$, $C_0^*$, $C_i^*$ | *None* |
| | $k_{on}$ | *None* | $k_{off}$, $k_p$, $r$, $r_p$, $T^*$, $L^*$, $C_0^*$, $C_i^*$ | *None* | $k_{off}$, $k_p$, $r$, $r_p$, $T^*$, $L^*$, $C_0^*$, $C_i^*$ |
| | $k_p$ | *None* | $k_{on}$, $k_{off}$, $r$, $r_p$, $T^*$, $L^*$, $C_0^*$, $C_i^*$ | *None* | $k_{on}$, $k_{off}$, $r$, $r_p$, $T^*$, $L^*$, $C_0^*$, $C_i^*$ |
| | $T(t)$, $k_{on}$, $k_p$ | $k_{off}$, $r$, $r_p$, $L^*$, $C_0^*$, $C_i^*$ | *None* | $\boldsymbol{k_{off}}$, $\boldsymbol{r}$, $\boldsymbol{r_p}$, $\boldsymbol{L^*}$, $\boldsymbol{C_0^*}$, $\boldsymbol{C_i^*}$ | *None* |
| | $T_T$ | *None* | $k_{on}$, $k_{off}$, $k_p$, $r$, $r_p$, $L^*$, $C_0^*$, $C_i^*$ | *None* | $k_{on}$, $k_{off}$, $k_p$, $r$, $r_p$, $L^*$, $C_0^*$, $C_i^*$ |
| | $T_T$, $k_{on}$, $k_p$ | *None* | $k_{off}$, $r$, $r_p$, $L^*$, $C_0^*$, $C_i^*$ | *None* | $k_{off}$, $r$, $r_p$, $L^*$, $C_0^*$, $C_i^*$ |
| **ZU** | — | n.d. | n.d. | $k_{on}$, $k_{off}$, $k_+$, $k_-$, $\lambda$, $T_p^*$ $L^*$, $T^*$, $D^*$, $P^*$, $C_0^*$ | *None* |
| | $T(t)$ | n.d. | n.d. | $k_{on}$, $k_{off}$, $k_+$, $k_-$, $\lambda$, $T_p^*$ $L^*$, $D^*$, $P^*$, $C_0^*$ | *None* |
| | $k_{on}$ | n.d. | n.d. | $k_{off}$, $k_+$, $k_-$, $\lambda$, $T_p^*$ $L^*$, $T^*$, $D^*$, $P^*$, $C_0^*$ | *None* |
| | $T(t)$, $k_{on}$ | n.d. | n.d. | $k_{off}$, $k_+$, $k_-$, $\lambda$, $T_p^*$ $L^*$, $D^*$, $P^*$, $C_0^*$ | *None* |
| | $T_T$ | n.d. | n.d. | $k_{on}$, $k_{off}$, $k_+$, $k_-$, $\lambda$, $T_p^*$ $L^*$, $D^*$, $P^*$, $C_0^*$ | *None* |
| | $T_T$, $k_{on}$ | n.d. | n.d. | $k_{off}$, $k_+$, $k_-$, $\lambda$, $T_p^*$ $L^*$, $D^*$, $P^*$, $C_0^*$ | *None* |

**Table S2 (cont.):** Identifiability results with measurements of $EC_{50}$ and $E_{max}$. Variables with an asterisk (*) denote initial conditions. In the "Id" columns, variables in boldface indicate structural global identifiability (SGI). The index $i$ goes from 1 to $N$.

| | | **Known output** | | | |
|---|---|---|---|---|---|
| | **Additional known output** | $EC_{50}$ | | $E_{max}$ | |
| **Models** | | Id | Non-id | Id | Non-id |
| **KPZU** | — | n.d. | n.d. | $k_{on}$, $k_{off}$, $k_+$, $k_-$, $k_p$, $\lambda$, $L^*$, $T^*$, $D^*$, $P^*$, $C_0^*$, $C_i^*$ | *None* |
| | $T(t)$ | n.d. | n.d. | $k_{on}$, $k_{off}$, $k_+$, $k_-$, $k_p$, $\lambda$, $L^*$, $D^*$, $P^*$, $C_0^*$, $C_i^*$ | *None* |
| | $k_{on}$ | n.d. | n.d. | $k_{off}$, $k_+$, $k_-$, $k_p$, $\lambda$, $L^*$, $T^*$, $D^*$, $P^*$, $C_0^*$, $C_i^*$ | *None* |
| | $k_p$ | n.d. | n.d. | $k_{on}$, $k_{off}$, $k_+$, $k_-$, $\lambda$, $L^*$, $T^*$, $D^*$, $P^*$, $C_0^*$, $C_i^*$ | *None* |
| | $T(t)$, $k_{on}$, $k_p$ | n.d. | n.d. | $k_{off}$, $k_+$, $k_-$, $\lambda$, $L^*$, $D^*$, $P^*$, $C_0^*$, $C_i^*$ | *None* |
| | $T_T$ | n.d. | n.d. | $k_{on}$, $k_{off}$, $k_+$, $k_-$, $k_p$, $\lambda$, $L^*$, $D^*$, $P^*$, $C_0^*$, $C_i^*$ | *None* |
| | $T_T$, $k_{on}$, $k_p$ | n.d. | n.d. | $k_{off}$, $k_+$, $k_-$, $\lambda$, $L^*$, $D^*$, $P^*$, $C_0^*$, $C_i^*$ | *None* |
| **KPC** | — | $k_{on}$, $k_{off}$, $k_+$, $k_-$, $k_p$, $\lambda$, $X^*$, $T^*$, $D^*$, $C_0^*$, $C_i^*$ | *None* | *None* | $k_{on}$, $k_{off}$, $k_+$, $k_-$, $k_p$, $\lambda$, $X^*$, $T^*$, $D^*$, $C_0^*$, $C_i^*$ |
| | $T(t)$ | $k_{on}$, $k_{off}$, $k_+$, $k_-$, $k_p$, $\lambda$, $X^*$, $D^*$, $C_0^*$, $C_i^*$ | *None* | $k_{on}$, $k_{off}$, $k_+$, $k_-$, $k_p$, $\lambda$, $X^*$, $D^*$, $C_0^*$, $C_i^*$ | *None* |
| | $k_{on}$ | $k_{off}$, $k_+$, $k_-$, $k_p$, $\lambda$, $X^*$, $T^*$, $D^*$, $C_0^*$, $C_i^*$ | *None* | *None* | $k_{off}$, $k_+$, $k_-$, $k_p$, $\lambda$, $X^*$, $T^*$, $D^*$, $C_0^*$, $C_i^*$ |
| | $k_p$ | $k_{on}$, $k_{off}$, $k_+$, $k_-$, $\lambda$, $X^*$, $T^*$, $D^*$, $C_0^*$, $C_i^*$ | *None* | *None* | $k_{on}$, $k_{off}$, $k_+$, $k_-$, $\lambda$, $X^*$, $T^*$, $D^*$, $C_0^*$, $C_i^*$ |
| | $T(t)$, $k_{on}$, $k_p$ | $k_{off}$, $k_+$, $k_-$, $\lambda$, $X^*$, $D^*$, $C_0^*$, $C_i^*$ | *None* | $k_{off}$, $k_+$, $k_-$, $\lambda$, $X^*$, $D^*$, $C_0^*$, $C_i^*$ | *None* |
| | $T_T$ | $k_{on}$, $k_{off}$, $k_+$, $k_-$, $k_p$, $\lambda$, $X^*$, $D^*$, $C_0^*$, $C_i^*$ | *None* | *None* | $k_{on}$, $k_{off}$, $k_+$, $k_-$, $k_p$, $\lambda$, $X^*$, $D^*$, $C_0^*$, $C_i^*$ |
| | $T_T$, $k_{on}$, $k_p$ | $k_{off}$, $k_+$, $k_-$, $\lambda$, $X^*$, $D^*$, $C_0^*$, $C_i^*$ | *None* | *None* | $k_{off}$, $k_+$, $k_-$, $\lambda$, $X^*$, $D^*$, $C_0^*$, $C_i^*$ |
| **ST** $N=2$ | — | *None* | $\lambda$, $\varphi$, $\sigma$, $k_{eff}$, $k_i$, $S^*$, $T^*$, $T_p^*$, $L$ | *None* | $\lambda$, $\varphi$, $\sigma$, $k_{eff}$, $k_i$, $S^*$, $T^*$, $T_p^*$, $L$ |
| | $T(t)$ | $\lambda$, $\varphi$, $\sigma$, $k_{eff}$, $S^*$, $L$ | $k_i$, $T_p^*$ | $\lambda$, $\varphi$, $\sigma$, $k_i$, $S^*$ | $k_{eff}$, $T_p^*$, $L$ |
| | $k_{\mathrm{eff}}$ | *None* | $\lambda$, $\varphi$, $\sigma$, $k_i$, $S^*$, $T^*$, $T_p^*$, $L$ | *None* | $\lambda$, $\varphi$, $\sigma$, $k_i$, $S^*$, $T^*$, $T_p^*$, $L$ |
| | $T(t)$, $k_{\mathrm{eff}}$ | $\lambda$, $\varphi$, $\sigma$, $k_{eff}$, $S^*$, $L$ | $k_i$, $T_p^*$ | $\lambda$, $\varphi$, $\sigma$, $k_i$, $S^*$, $L$ | $T_p^*$ |
| | $T_T$ | $\lambda$, $\varphi$, $\sigma$, $k_{eff}$, $k_i$, $S^*$, $T^*$, $T_p^*$, $L$ | *None* | $\lambda$, $\varphi$, $\sigma$, $k_i$, $S^*$, $T^*$, $T_p^*$ | $k_{eff}$, $L$ |
| | $T_T$, $k_{\mathrm{eff}}$ | $\lambda$, $\varphi$, $\sigma$, $k_i$, $S^*$, $T^*$, $T_p^*$, $L$ | *None* | $\lambda$, $\varphi$, $\sigma$, $k_i$, $S^*$, $T^*$, $T_p^*$, $L$ | *None* |

**Table S3:** Models' rank according to the number of output configurations that make them identifiable.

| Rank | Model | No. Output configurations | | |
|---|---|---|---|---|
| | | **One output** | **Two outputs (simultaneous)** | **Total** |
| 1 | *KPR-IR* | 5 | 4 | 9 |
| 2 | *KPR-SS* and *KPC* | 3 | 4 | 7 |
| 3 | *KPR-SAC* | 2 | 4 | 6 |
| | *ZU* and *KPZU* | 3 | 3 | 6 |
| 4 | *Occ* | 1 | 4 | 5 |
| 5 | *KPR-LS* | 1 | 2 | 3 |
| 6 | *KPR-1* | 0 | 2 | 2 |
| 7 | *ST* | 1 | 0 | 1 |
| 8 | *KPR-LS-IFF*, *KPR-NF1*, *KPR-NF2* | 0 | 0 | 0 |

**Table S4:** Output configurations' rank according to the number of models that become identifiable under them.

| Rank | Output configuration | No. Identifiable models |
|---|---|---|
| 1 | $R(t) + T(t)$ and $R(t) + T_T$ | 9 |
| 2 | $R(t)$ and $E_{max} + T(t)$ | 7 |
| 3 | $T(t)$ | 6 |
| 4 | $EC_{50} + T(t)$ | 5 |
| 5 | $EC_{50}$ | 4 |
| 6 | $E_{max}$ | 3 |
| 7 | $T_T$ | 2 |

#### 3.2 Sensitivity results

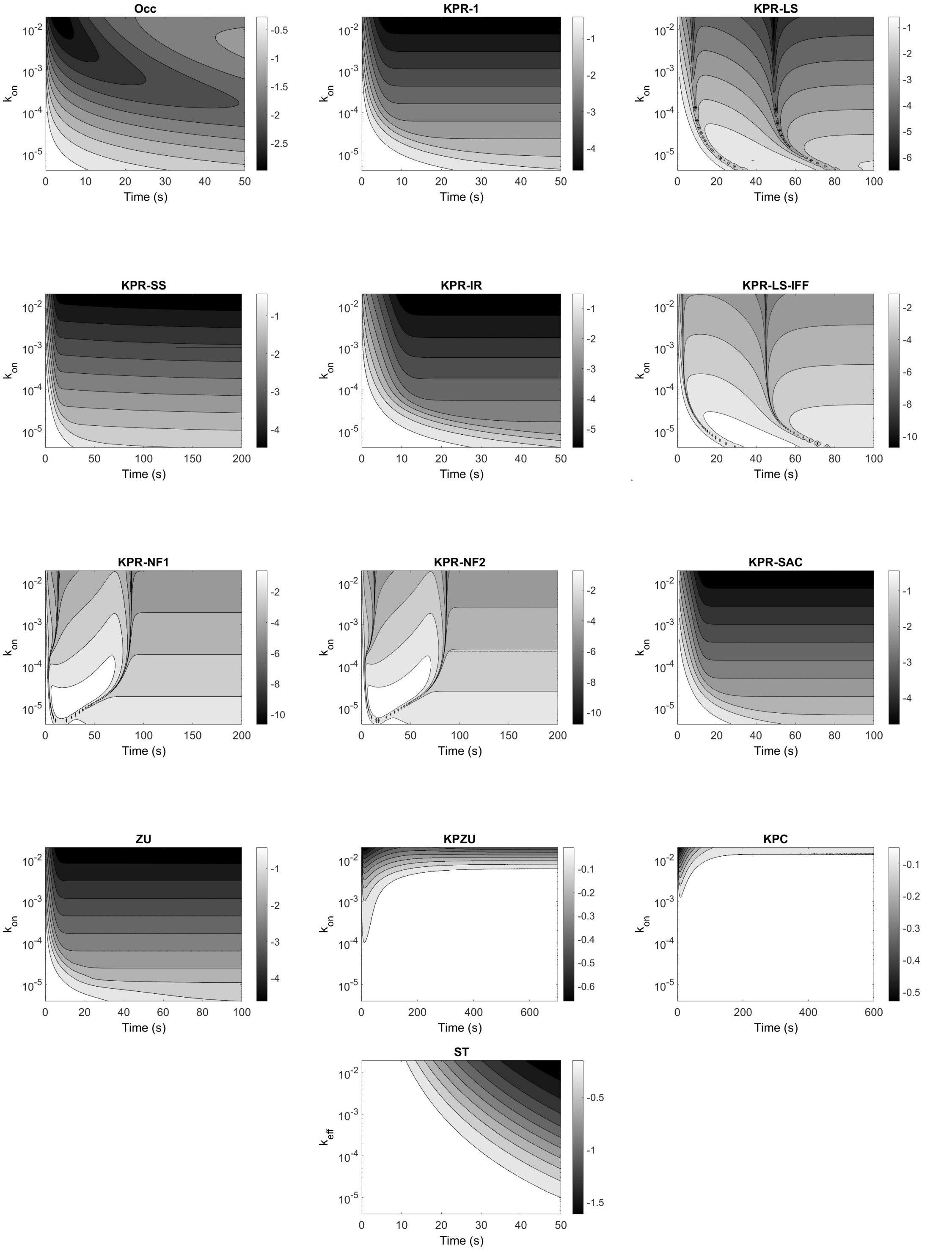


**Figure S4:** Contour map of the sensitivity of the response $R(t)$ to the binding rate $k_{on}$ in all models. The plotted results and the sensitivity gradient bars are in base-10 logarithmic scale. Note that the $ST$ model was analyzed with respect to $k_{eff}$, a compound parameter proportional to $k_{on}$.

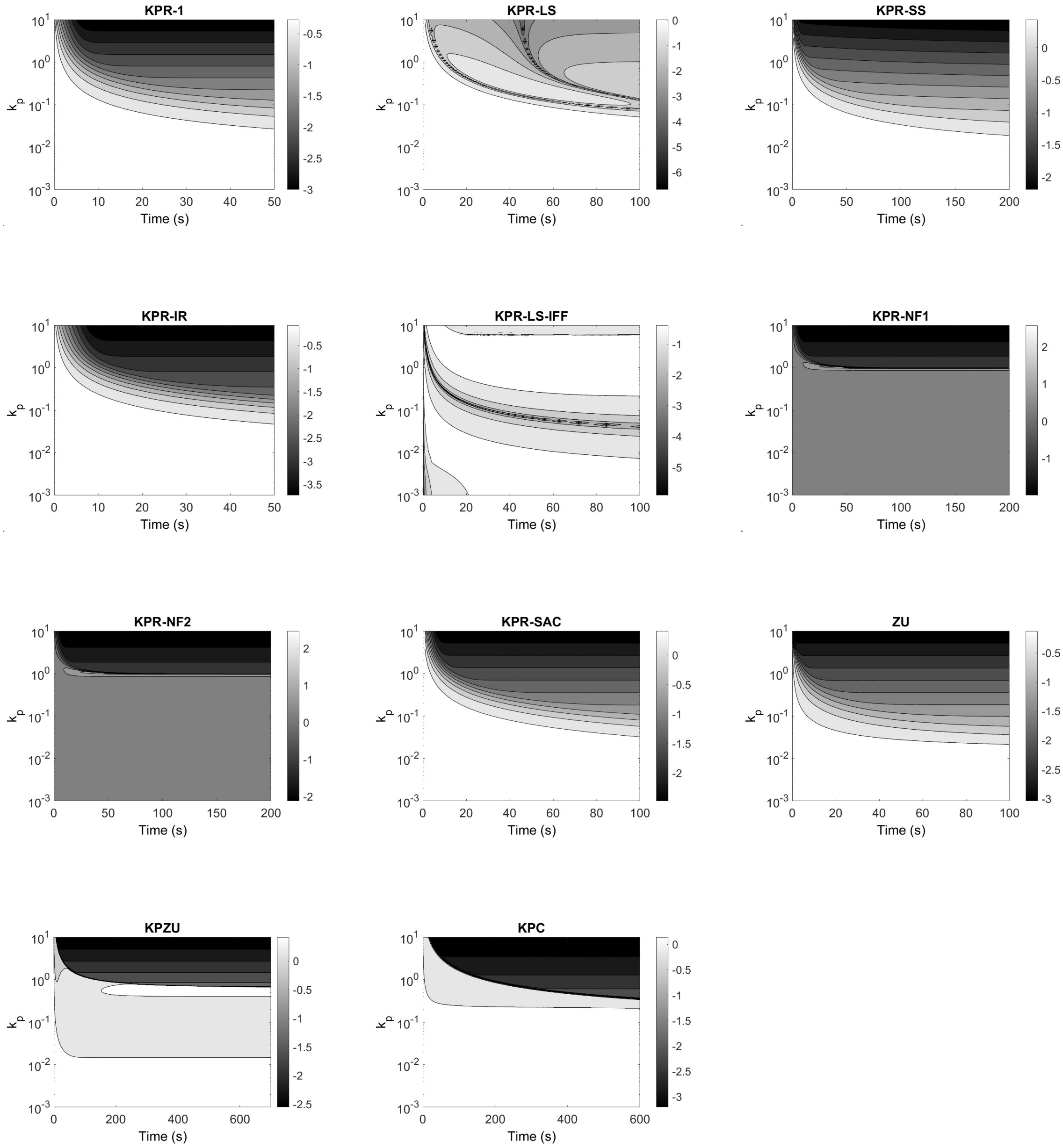


**Figure S5:** Contour map of the sensitivity of the response $R(t)$ to the phosphorylation rate $k_p$ in all models that include this parameter. The plotted results and the sensitivity gradient bars are in base-10 logarithmic scale.

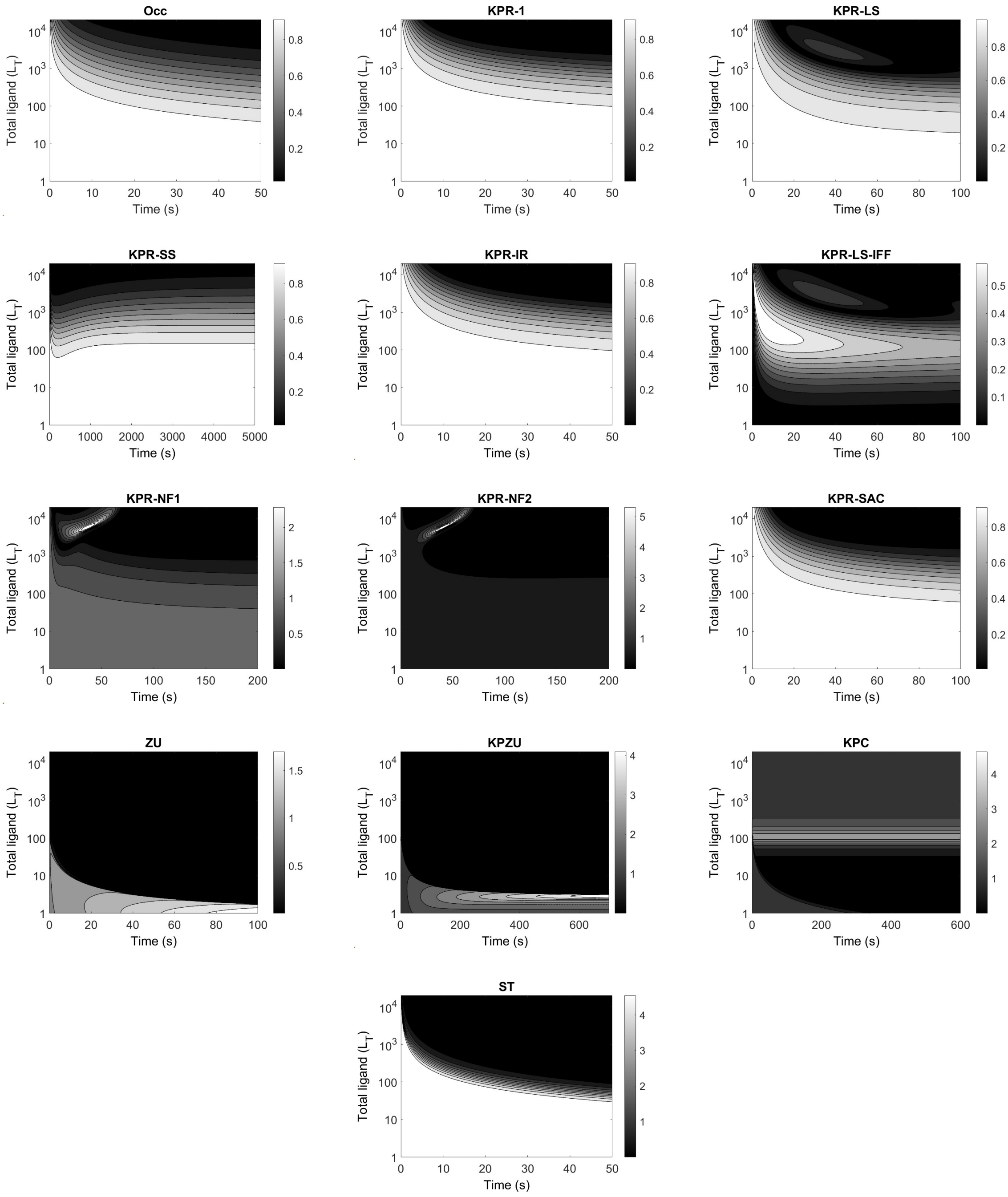


**Figure S6:** Contour map of the sensitivity of the response $R(t)$ to the antigen dose, $L_T$, in all models. The sensitivity gradient bars are in arithmetic scale.

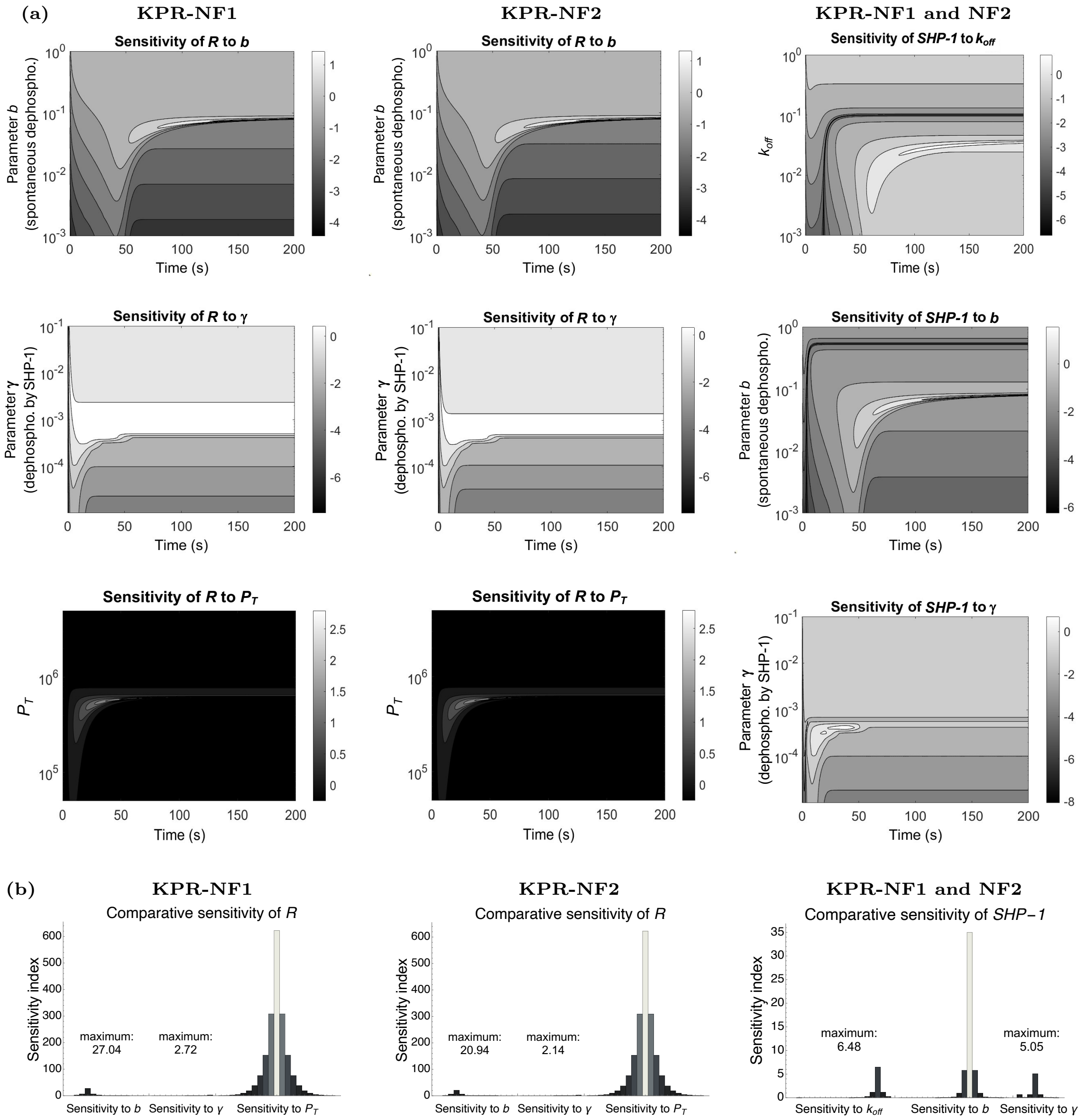


**Figure S7:** Sensitivity analysis for *KPR-NF1* and *KPR-NF2* models with respect to model-specific quantities. **(a)** Density plots with contour lines. Left and middle columns: sensitivity of $R(t)$ to the spontaneous dephosphorylation rate of $C_i$ complexes, $b$ (top), and to the SHP-1–induced dephosphorylation rate of $C_i$ complexes, $\gamma$ (bottom), respectively, in models *KPR-NF1* and *KPR-NF2*. Right column: sensitivity in both models of the amount of active SHP-1 phosphatase, $P(t)$, to parameters $k_{off}$ (top), $b$ (middle), and $\gamma$ (bottom). **(b)** Sensitivity profiles obtained from the graphs in (a). Each histogram is derived from its corresponding graph by taking, at a particular time, the parameter values at each contour line. In each histogram, bars from left to right correspond to the sensitivity levels, from maximum to minimum parameter values. Note that the width of the bars is constant and hence is not related to the difference of values of a parameter between consecutive contour lines. The plotted results and the sensitivity gradient bars in (a) are in base-10 logarithmic scale.

[3] Xabier Rey Barreiro, Jose Faro, and Alejandro F. Villaverde. Response metrics of t-cell activation models. *bioRxiv*, 2025.

[4] Jose Faro, Mario Castro, and Carmen Molina-París. A unifying mathematical framework for experimental tcr-pmhc kinetic constants. *Scientific Reports*, 7(1), April 2017.

[5] Paul François, Guillaume Voisinne, Eric D Siggia, Grégoire Altan-Bonnet, and Massimo Vergassola. Phenotypic model for early t-cell activation displaying sensitivity, specificity, and antagonism. *Proceedings of the National Academy of Sciences*, 110(10):E888–E897, 2013.
